# Assessing the accuracy of octanol-water partition coefficient predictions in the SAMPL6 Part II log *P* Challenge

**DOI:** 10.1101/2020.01.20.913178

**Authors:** Mehtap Işık, Teresa Danielle Bergazin, Thomas Fox, Andrea Rizzi, John D. Chodera, David L. Mobley

**Author notes:** **For correspondence:** (MI). These authors contributed equally to this work (Mehtap Işık, Teresa Danielle Bergazin).

## Abstract

The SAMPL Challenges aim to focus the biomolecular and physical modeling community on issues that limit the accuracy of predictive modeling of protein-ligand binding for rational drug design. In the SAMPL5 log *D* Challenge, designed to benchmark the accuracy of methods for predicting drug-like small molecule transfer free energies from aqueous to nonpolar phases, participants found it difficult to make accurate predictions due to the complexity of protonation state issues. In the SAMPL6 log *P* Challenge, we asked participants to make blind predictions of the octanol-water partition coefficients of neutral species of 11 compounds and assessed how well these methods performed absent the complication of protonation state effects. This challenge builds on the SAMPL6 p*K*_a_ Challenge, which asked participants to predict p*K*_a_ values of a superset of the compounds considered in this log *P* challenge. Blind prediction sets of 91 prediction methods were collected from 27 research groups, spanning a variety of quantum mechanics (QM) or molecular mechanics (MM)-based physical methods, knowledge-based empirical methods, and mixed approaches. There was a 50% increase in the number of participating groups and a 20% increase in the number of submissions compared to the SAMPL5 log *D* Challenge. Overall, the accuracy of octanol-water log *P* predictions in SAMPL6 Challenge was higher than cyclohexane-water log *D* predictions in SAMPL5, likely because modeling only the neutral species was necessary for log *P* and several categories of method benefited from the vast amounts of experimental octanol-water log *P* data. There were many highly accurate methods: 10 diverse methods achieved RMSE less than 0.5 log *P* units. These included QM-based methods, empirical methods, and mixed methods with physical modeling supported with empirical corrections. A comparison of physical modeling methods showed that QM-based methods outperformed MM-based methods. The average RMSE of the most accurate five MM-based, QM-based, empirical, and mixed approach methods based on RMSE were 0.92±0.13, 0.48±0.06, 0.47±0.05, and 0.50±0.06, respectively.

## 1 Introduction

The development of computational biomolecular modeling methodolgoies is motivated by the goal of enabling quantitative molecular design, prediction of properties and biomolecular interactions, and achieving a detailed understanding of mechanisms (chemical and biological) via computational predictions. While many approaches are available for making such predictions, methods often suffer from poor or unpredictable performance, ultimately limiting their predictive power. It is often difficult to know which method would give the most accurate predictions for a target system without extensive evaluation of methods. However, such extensive comparative evaluations are infrequent and difficult to perform, partly because no single group has expertise in or access to all relevant methods and also because of the scarcity of blind experimental data sets that would allow prospective evaluations. In addition, many publications which report method comparisons for a target system constructs these studies with the intention of highlighting the success of a method being developed.

The SAMPL (Statistical Assessment of the Modeling of Proteins and Ligands) Challenges [http://samplchallenges.github.io] provide a forum to test and compare methods with the following goals:

1. Determine prospective predictive power rather than accuracy in retrospective tests.
2. Allow a head to head comparison of a wide variety of methods on the same data.

Regular SAMPL challenges focus attention on modeling areas that need improvement, and sometimes revisit key test systems, providing a crowdsourcing mechanism to drive progress. Systems are carefully selected to create challenges of gradually increasing complexity spanning between prediction objectives that are tractable and that are understood to be slightly beyond the capabilities of contemporary methods. So far, most frequent SAMPL challenges have been on solvation and binding systems. Iterated blind prediction challenges have played a key role in driving innovations in the prediction of physical properties and binding. Here we report on a SAMPL6 log *P* Challenge on octanol-water partition coefficients, treating molecules resembling fragments of kinase inhibitors. This is a follow-on to the earlier SAMPL6 p*K*_a_ Challenge which included the same compounds.

The partition coefficient describes the equilibrium concentration ratio of the neutral state of a substance between two phases:

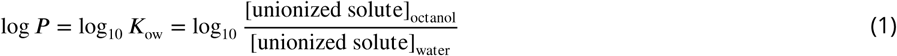

The log *P* challenge examines how well we model transfer free energy of molecules between different solvent environments in the absence of any complications coming from predicting protonation states and p*K*_a_ values. Assessing log *P* prediction accuracy also allows evaluating methods for modeling protein-ligand affinities in terms of how well they capture solvation effects.

### 1.1 SAMPL Challenge History and Motivation

The SAMPL blind challenges aim to focus the field of quantitative biomolecular modeling on major issues that limit the accuracy of protein-ligand binding prediction. Companion exercises such as the Drug Design Data Resource (D3R) blind challenges aim to assess the current accuracy of biomolecular modeling methods in predicting bound ligand poses and affinities on real drug discovery project data. D3R blind challenges serve as an accurate barometer for accuracy. However, due to the conflation of multiple accuracy-limiting problems in these complex test systems it is difficult to derive clear insights into how to make further progress towards better accuracy.

Instead, SAMPL seeks to isolate and focus attention on individual accuracy-limiting issues. We aim to field blind challenges just at the limit of tractability in order to identify underlying sources of error and help overcome these challenges. Working on similar model systems or the same target with new blinded datasets in multiple iterations of prediction challenges maximize our ability to learn from successes and failures. Often, these challenges focus on physical properties of high relevance to drug discovery in their own right, such as partition or distribution coefficients critical to the development of potent, selective, and bioavailable compounds.

The partition coefficient (log *P*) and the distribution coefficient (log *D*) are driven by the free energy of transfer from an aqueous to a nonpolar phase. Transfer free energy of only neutral species are considered for log *P*, whereas both neutral and ionizable species contribute to log *D*. Such solute partitioning models are a simple proxy for the transfer free energy of a drug-like molecule to a relatively hydrophobic receptor binding pocket, in the absence of specific interactions. Protein-ligand binding equilibrium is analogous to partitioning of a small molecule between two environments: protein binding site and aqueous phase. Methods that employ thermodynamic cycles, such as free energy calculations, can therefore use similar strategies for calculating binding affinities and partition coefficients, and given the similarity in technique and environment, we might expect the accuracy on log *P* and log *D* may be related to the accuracy expected from binding calculations, or at least a lower bound for the error these techniques might make in more complex protein-ligand binding phenomena. Evaluating log *P* or log *D* predictions makes it far easier to probe the accuracy of computational tools used to model protein-ligand interactions and to identify sources of error to be corrected. For physical modeling approaches, evaluation of partition coefficient predictions comes with the additional advantage of separating force field accuracy from protonation state modeling challenges.

#### The SAMPL5 log *D* challenge uncovered surprisingly large modeling errors

Hydration free energies formed the basis of several previous SAMPL challenges, but distribution coefficients (log *D*) capture many of the same physical effects—namely, solvation in the respective solvents—and thus replaced hydration free energies in SAMPL5 [1, 2]. This choice was also driven due to a lack of ongoing experimental work with the potential to generate new hydration free energy data for blind challenges. Octanol-water log *D* is also a property relevant to drug discovery, often used as a surrogate for lipophilicity, further justifying its choice for a SAMPL challenge. The SAMPL5 log *D* Challenge allowed decoupled evaluation of small molecule solvation models (captured by the transfer free energy between environments) from other issues, such as the sampling of slow receptor conformational degrees of freedom. This blind challenge generated considerable insight into the importance of various physical effects [1, 2]; see the SAMPL5 special issue (https://link.springer.com/journal/10822/30/11/page/1) for more details.

The SAMPL5 log *D* Challenge used cyclohexane as an apolar solvent, partly to further simplify this challenge by avoiding some complexities of octanol. In particlar, log *D* is typically measured using water-saturated octanol for the nonaqueous phase, which can give rise to several challenges in modeling accuracy such as a heterogeneous environment with potentially micelle-like bubbles [3–6], resulting in relatively slow solute transitions between environments [4, 7]. The precise water content of wet octanol is unknown, as it is affected by environmental conditions such as temperature as well as the presence of solutes, the organic molecule of interest, and salts (added to control pH and ionic strength). Inverse micelles transiently formed in wet octanol create spatial heterogeneity and can have long correlation times in molecular dynamics simulations, potentially presenting a challenge to modern simulation methods[3–6], resulting in relatively slow solute transitions between environments [4, 7].

Performance in the SAMPL5 log *D* Challenge was much poorer than the organizers initially expected—and than would have been predicted based on past accuracy in hydration free energy predictions—and highlighted the difficulty of accurately accounting for protonation state effects [2]. In many SAMPL5 submissions, many participants treated distribution coefficients (log *D*) as if they were asked to predict partition coefficients (log *P*). The difference between log *D* (reflects the transfer free energy at a given pH including the effects of accessing all equilibrium protonation states of the solute in each phase) and log *P* (reflects aqueous-to-apolar phase transfer free energies of the neutral species only) proved particularly important. In some cases, other effects like the presence of a small amount of water in cyclohexane may also have played a role.

Because the SAMPL5 log *D* Challenge highlighted the difficulty in correctly predicting transfer free energies involving protonation states (the best methods obtained an RMSE of 2.5 log units [2]), the SAMPL6 Challenge aimed to further subdivide the assessment of modeling accuracy into two challenges: A small-molecule p*K*_a_ prediction challenge [8] and a log *P* challenge. The SAMPL6 p*K*_a_ Challenge asked participants to predict microscopic and macroscopic acid dissociation constants (p*K*_a_s) of 24 small organic molecules and concluded in early 2018. Details of the challenge are maintained on the GitHub repository (https://github.com/samplchallenges/SAMPL6/). p*K*_a_ prediction proved to be difficult. A large number of methods showed RMSE in the range of 1-2 p*K*_a_units, with only a handful achieving less than 1 p*K*_a_ unit. These results were in line with expectations from the SAMPL5 Challenge about protonation state predictions being one of the major sources of error for log *D*. But the present challenge allows us delve deeper into modeling the solvation of neutral species and focus on log *P*.

#### The SAMPL6 log *P* Challenge focused on small molecules resembling kinase inhibitor fragments

By measuring the log *P* of a series of compounds resembling fragments of kinase inhibitors—a subset of those used in the SAMPL6 p*K*_a_ Prediction Challenge—we sought to assess the limitations of force field accuracy in modeling transfer free energies of drug-like molecules in binding-like processes. This time, the challenge featured octanol as the apolar medium to assess whether wet octanol presented as much of a problem as was previously suspected. Participants are asked to predict the partition coefficient (log *P*) of the neutral species between octanol and water phases. Here we focus on different aspects of the challenge, particularly the staging, analysis, results, and lessons learned. Experimental work for collecting the log *P* values are described elsewhere [9]. One of the goals of this challenge is to encourage prediction of model uncertainties (an estimate of the inaccuracy with which your model predicts the physical property), since the ability to tell when methods will be successful or not would be very useful for increasing the application potential and impact of computational methods.

#### The SAMPL challenges aim to advance predictive quantitative models

The SAMPL challenges have a key focus on lessons learned. In principle, they are a challenge or competition, but we see it as far more important to learn how to improve accuracy than to announce the top-performing methods. To aid in learning as much as possible, this overview paper provides an overall assessment of performance and some analysis of the relative performance of different methods in different categories, provides some insights into lessons we have learned (and/or other participants have learned). Additionally, this work presents our own reference calculations which provide points of comparison for participants (some relatively standard and some more recent, especially in the physical category) and also allow us to provide some additional lessons learned. The data, from all participants and all reference calculations, is made freely available (see Section 6) to allow others to compare methods retrospectively and dig into additional lessons learned.

### 1.2 Common computational approaches for predicting log *P*

Many methods have been developed to predict octanol-water log *P* values of small organic molecules including physical modeling (QM and MM-based methods) and knowledge-based empirical prediction approaches (atom-contribution approaches and QSPR). There are also log *P* prediction methods that combine the strengths of physical and empirical approaches. Here, we briefly highlight some of the major ideas and background behind physical and empirical log *P* prediction methods.

#### 1.2.1 Physical modeling approaches for predicting log *P*

Physical approaches begin with a detailed atomistic model of the solute and its conformation and attempt to estimate partitioning behavior directly from that. Details depend on the approach employed.

##### 1.2.1.1. Quantum mechanical (QM) approaches for predicting log *P*

QM approaches to solvation modeling utilize numerical solution of the Schrödinger equation to estimate solvation free energies (and thereby partitioning) directly from first principles. There are a number of approaches for these calculations, and discussing them is outside the scope of this work. However, it is important to note that direct solution of the underlying equations, especially when coupled with dynamics, becomes impractical for large systems such as molecules in solution. So, several approximations must be made before such approaches can be applied to estimating phase transfer free energies. These typical approximations include assuming the solute has one or a small number of dominant conformations in each phase being considered, and using an implicit solvent model to represent the solvent. The basis set and level of theory can be important choices and can significantly affect accuracy of calculated values. Additionally, protonation or tautomerization state selected as an input can also introduce errors. With QM approaches possible protonation states and tautomers can be evaluated to find the lowest energy state in each solvent. However, if these estimates are erroneous, any errors will propagate into the final transfer free energy and log *P* predictions.

Implicit solvent models can be used, in the context of the present SAMPL, both to represent water and octanol. Such models are often parameterized—sometimes highly so—based on experimental solvation free energy data. This means that such models perform well for solvents (and solute chemistries) where solvation free energy data is abundant (as in the present challenge) but are often less successful when far less training data is available. In this respect, QM methods, by virtue of the solvent model, have some degree of overlap with the empirical methods discussed further below.

Several solvent models are particularly common, and in the present challenge two were particularly commonly employed. One was Marenich, Cramer and Truhlar’s SMD solvation model [10], which derives its electrostatics from the widely used IEF-PCM model and was empirically trained on various solutes/solvents utilizing a total of some 2821 different solvation data points. This has been employed in various SAMPL challenges in the past in the context of calculation of hydration free energies, including the earliest SAMPL challenges [11, 12]. Others in the Cramer-Truhlar series of solvent models were also used, including the 2012 SM12 solvation model, which is based on the generalized born (GB) approximation [13]. Another set of submissions also used the reference interaction site model (RISM) integral equation approach, discussed further below.

The COSMO-RS solvation model is another method utilized in this context which covers a particularly broad range of solvents, typically quite well [14–18]. In the present challenge, a “Cosmoquick” variant was also applied and falls into the “Mixed” method category, as it utilizes additional empirical adjustments. The COSMO-RS implementation of COSMOtherm takes into account conformational effects to some extent; the chemical potential in each phase is computed using the Boltzmann weights of a fixed set of conformers.

In general, while choice of solvation model can be a major factor impacting QM approaches, the neglect of conformational changes means these approaches typically (though not always) neglect any possibility of significant change of conformation on transfer between phases and they simply estimate solvation by the difference in (estimated) solvation free energies for each phase of a fixed conformation. Additionally, solute entropy is often neglected, assuming the single-conformation solvation free energy plays the primary role in driving partitioning between phases. In addition to directly estimating solvation, QM approaches can also be used to drive the selection of the gas- or solution-phase tautomer, and thus can be used to drive the choice of inputs for MM approaches discussed further below.

###### Integral equation-based approaches

Integral equation approaches provide an alternate approach to solvation modeling (for both water and non-water solvents) and have been applied in SAMPL challenges within both the MM and QM frameworks [19–21]. In this particular challenge, however, the employed approaches were entirely QM, and utilized the reference interaction site model (RISM) approach [22–24]. Additionally, as noted above, the IEF-PCM model used by the SMD solvation model (discussed above) is also an integral equation approach. Practical implementation details mean that RISM approaches typically have one to a few adjustable parameters (e.g. four [25]) which are empirically tuned to experimental solvation free energies, in contrast to the SMD and SM-n series of solvation models which tend to have a larger number of adjustable parameters and thus require larger training sets. In this particular SAMPL challenge, RISM participation was limited to embedded cluster EC-RISM methods [19, 22, 26], which combine RISM with a quantum mechanical treatment of the solute.

##### 1.2.1.2. Molecular mechanics (MM) approaches for predicting log *P*

MM approaches to computing solvation and partition free energies (and thus log *P* values), as typically applied in SAMPL, use a force field or energy model which gives the energy (and, usually, forces) in a system as a function of the atomic positions. These models include all-atom fixed charge additive force fields, as well as polarizable force fields. Such approaches typically (though not always) are applied in a dynamical framework, integrating the equations of motion to solve for the time evolution of the system, though Monte Carlo approaches are also possible.

MM-based methods are typically coupled with free energy calculations to estimate partitioning. Often, these are so-called alchemical methods which utilize a non-physical thermodynamic cycle to estimate transfer between phases, though pulling-based techniques which directly model phase transfer are in principle possible [27, 28]. Such free energy methods allow detailed all-atom modeling of each phase, and compute the full free energy of the system, in principle (in the limit of adequate sampling) providing the correct free energy difference given the choice of energy model (“force field”). However, adequate sampling can sometimes prove difficult to achieve.

Key additional limitations facing MM approaches are the accuracy of the force field, the fact that protonation state/tautomer is generally selected as an input and held fixed (meaning that incorrect assignment or multiple relevant states can introduce significant errors), and timescale—simulations only capture motions that are faster than simulation timescale. However, these approaches *do* capture conformational changes on phase transfer, as long as such changes occur on timescales faster than the simulation timescale.

#### 1.2.2 Empirical log *P* predictions

Due to the importance of accurate log *P* predictions, ranging from pharmaceutical sciences to environmental hazard assessment, a large number of empirical models to predict this property have been developed and reviewed [29–31]. An important characteristic of many of these methods is that they are very fast, so even large virtual libraries of molecules can be characterized.

In general, two main methodologies can be distinguished: group- or atom-contribution approaches, also called additive group methods, and quantitative structure-property relationship (QSPR) methods.

##### 1.2.2.1 Atom- and group-contribution approaches

Atom contribution methods, pioneered by Crippen in the late 1980s [32, 33], are the easiest to understand conceptually. These assume that each atom contributes a specific amount to the solvation free energy and that these contributions to log *P* are additive. Using a potentially large number of different atom types (typically in the order of 50-100), the log *P* is the sum of the individual atom types times the number of their occurrences in the molecule:

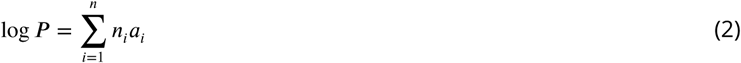

A number of log *P* calculation programs are based on this philosophy, including AlogP [34], AlogP98 [34], and moe_SlogP [35].

The assumption of independent atomic contributions fails for compounds with complex aromatic systems or stronger electronic effects. Thus correction factors and contributions from neighboring atoms were introduced to account for these shortcomings (e.g. in XlogP [36–38] and SlogP [35].

In contrast, in group contribution approaches, log *P* is calculated as a sum of group contributions, usually augmented by correction terms that take into account intramolecular interactions. Thus, the basic equation is

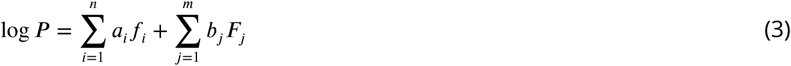

where the first term describes the contribution of the fragments *f*_*i*_ (each occurring *a*_*i*_ times), the second term gives the contributions of the correction factors *F*_*j*_ occurring *b*_*j*_ times in the compound. Group contribution approaches assume that the details of the electronic or intermolecular structure can be better modeled with whole fragments. However, this breaks down when molecular features are not covered in the training set. Prominent examples of group contribution approaches include clogP [39–42], KlogP [43], ACD/logP [44] and KowWIN [45].

clogP is probably one of the most widely used log *P* calculation programs [39–41]. clogP relies on fragment values derived from measured data of simple molecules, e.g., carbon and hydrogen fragment constants were derived from measured values for hydrogen, methane, and ethane. For more complex hydrocarbons, correction factors were defined to minimize the difference to the experimental values. These can be structural correction factors taking into account bond order, bond topology (ring/chain/branched) or interaction factors taking into account topological proximity of certain functional groups, electronic effects through *π*-bonds, or special ortho-effects.

##### 1.2.2.2 QSPR approaches

Quantitative structure-property relationships (QSPR) provide an entirely different category of approaches. In QSPR methods, a property of a compound is calculated from molecular features that are encoded by so-called molecular descriptors. Often, these theoretical molecular descriptors are classified as 0D-descriptors (constitutional descriptors, only based on the structural formula), 1D-descriptors (i.e. list of structural fragments, fingerprints), 2D-descriptors (based on the connection table, topological descriptors), and 3D-descriptors (based on the three-dimensional structure of the compound, thus conformation-dependent). Sometimes, this classification is extended to 4D-descriptors, which are derived from molecular interaction fields (e.g., GRID, CoMFA fields).

Over the years, a large number of descriptors have been suggested, with varying degrees of interpretability. Following the selection of descriptors, a regression model that relates the descriptors to the molecular property is derived by fitting the individual contributions of the descriptors to a dataset of experimental data; both linear and nonlinear fitting is possible. Various machine learning approaches such as random forest models, artificial neural network models, etc. also belong to this category. Consequently, a large number of estimators of this type have been proposed; some of the more well-known ones include MlogP [46] and VlogP [47].

#### 1.2.3 Expectations from different prediction approaches

Octanol-water log *P* literature data abounds, impacting our expectations. Given this abundance of data, in contrast to cyclohexane-water log *D* data, e.g., for the SAMPL5 log *D* Challenge, we expected higher accuracy here. Some sources of public octanol-water log *P* values include DrugBank [48], ChemSpider [49], PubChem, the NCI CACTUS databases [50, 51], and SRC’s PHYSPROP Database [52].

Our expectation was that empirical knowledge-based and other trained methods (implicit solvent QM, mixed methods) would outperform other methods in the present challenge as they are impacted directly by the availability of octanol-water data. Methods well trained to experimental octanol-water partitioning data should typically result in higher accuracy, if fitting is done well. The abundance of octanol-water data may also provide empirical and mixed approaches with an advantage over physical modeling methods. Current molecular mechanics-based methods and other methods not trained to experimental log *P* data ought to do worse in this challenge. Performance of strictly physical modeling based prediction methods might generalize better across other solvent types where training data is scarce, but that will not be tested by this challenge. In principle, molecular mechanics-based methods could also be fitted using octanol-water data as one of the targets for force field optimization, but present force fields have not made broad use of this data in fitting. Thus, top methods are expected to be from empirical knowledge-based, QM-based approaches and combination of QM-based and empirical approaches because of training data availability. These categories are broken out separately for analysis.

## 2 Challenge design and evaluation

### 2.1 Challenge structure

The SAMPL6 Part II Challenge was conducted as a blind prediction challenge on predicting octanol-water partition coefficients of 11 small molecules that resemble fragments of kinase inhibitors. The challenge molecule set was composed of small molecules with limited flexibility (less than 5 non-terminal rotatable bonds) and covers limited chemical diversity. There are six 4-aminoquinazolines, two benzimidazoles, one pyrazolo[3,4-d]pyrimidine, one pyridine, one 2-oxoquinoline substructure containing compounds with log *P* values in the range of 1.95–4.09. Information on experimental data collection is presented elsewhere [9].

The dataset composition was announced several months before the challenge including details of the measurement technique (potentiometric log *P* measurement, at room temperature, using water saturated octanol phase, and ionic strength-adjusted water with 0.15 M KCl [9]), but not the identity of the small molecules. The instructions and the molecule set were released at the challenge start date (Nov 1, 2018), and then submissions were accepted until March 22, 2019.

Following the conclusion of the blind challenge, the experimental data was made public on Mar 25, 2019 and results are first discussed in a virtual workshop (on May 16, 2019) [54] then later in an in person workshop (Joint D3R/SAMPL Workshop, San Diego, Aug 22-23, 2019). The purpose of the virtual workshop was to go over a preliminary evaluation of results, begin considering analysis and lessons learned, and nucleate opportunities for follow up and additional discussion. Part of the goal was to facilitate discussion so that participants can work together to maximize lessons learned in the lead up to an in-person workshop and special issue of a journal. The SAMPL6 log *P* Virtual Workshop video [54] and presentation slides [55] are available, as are organizer presentation slides from the joint D3R/SAMPL Workshop 2019 [56, 57] on the SAMPL Community Zenodo page (https://zenodo.org/communities/sampl/).

A machine-readable submission file format was specified for blind submissions. Participants were asked to report SAMPL6 Molecule IDs, predicted octanol-water log *P* values, the log *P* standard error of the mean (SEM), and model uncertainty. It was mandatory to submit predictions for all these values, including the estimates of uncertainty. The log *P* SEM captures the statistical uncertainty of the predicted method, and the model uncertainty is an estimate of how well prediction and experimental values will agree. Molecule IDs assigned in SAMPL6 p*K* _a_ Challenge were conserved in the challenge for the ease of reference.

Participants were asked to categorize their methods as belonging to one of four method categories — physical, empirical, mixed or other. The following are definitions provided to participants for selecting a method category: Empirical models are prediction methods that are trained on experimental data, such as QSPR, machine learning models, artificial neural networks etc. Physical models are prediction methods that rely on the physical principles of the system such as molecular mechanics or quantum mechanics based methods to predict molecular properties. Methods taking advantage of both kinds of approaches were asked to be reported as “Mixed”. The “other” category was for methods which do not match the previous ones. At the analysis stage, some categories were further refined, as discussed in Section 2.2.

The submission files also included fields for naming the method, listing the software utilized, and a free text method section for the detailed documentation of each method. Only one log *P* value for each molecule per submission and only full prediction sets were allowed. Incomplete submissions – such as for a subset of compounds – were not accepted. We highlighted various factors for participants to consider in their log *P* predictions. These included:

1. There is a significant partitioning of water into the octanol phase. The mole fraction of water in octanol was previously measured as 0.271±0.003 at 25°C [58].
2. The solutes can impact the distribution of water and octanol. Dimerization or oligomerization of solute molecules in one or more of the phases may also impact results [59].
3. log *P* measurements capture partition of neutral species which may consist of multiple tautomers with significant populations or the major tautomer may not be the one given in the input file.
4. Shifts in tautomeric state populations on transfer between phases are also possible.

Research groups were allowed to participate with multiple submissions, which allowed them to submit prediction sets to compare multiple methods or to investigate the effect of varying parameters of a single method. All blind submissions were assigned a 5-digit alphanumeric submission ID, which will be used throughout this paper and also in the evaluation papers of participants. These abbreviations are defined in Table 3.

**Table 1.** Methods used as reference calculations for the MM-based physical methods category. Please see Section 12.1.1 in the Supplementary Information for detailed description of physical reference methods.

| Submission ID | Approach | Force Field | Water Model | Octanol Phase | Number of Replicates |
| --- | --- | --- | --- | --- | --- |
| *REF01* | YANK, DFE protocol | GAFF 1.81 | TIP3P-FB | Wet | 3 |
| *REF02* | YANK, DFE protocol | GAFF 1.81 | TIP3P | Wet | 3 |
| *REF03* | YANK, DFE protocol | GAFF 1.81 | OPC | Wet | 3 |
| *REF04* | YANK, DFE protocol | smirnoff99Frosst 1.0.7 | TIP3P-FB | Wet | 3 |
| *REF05* | YANK, DFE protocol | smirnoff99Frosst 1.0.7 | TIP3P | Wet | 3 |
| *REF06* | YANK, DFE protocol | smirnoff99Frosst 1.0.7 | OPC | Wet | 3 |
| *REF07* | YANK, DFE protocol | GAFF 1.81 | TIP3P | Dry | 3 |
| *REF08* | YANK, DFE protocol | smirnoff99Frosst 1.0.7 | TIP3P | Dry | 3 |

**Table 2.** Methods used as reference calculations for the empirical log *P* prediction category. Please see section 12.1.3 in the Supplementary Information for a detailed description of empirical methods.

| Submission ID | Name | Vendor | Approach | Website |
| --- | --- | --- | --- | --- |
| REF09 | clogP (BioByte) | BioByte | group contributions | www.biobyte.com |
| REF13 | SlogP (MOE) | Chemical Computing Group | atomic contributions | www.chemcomp.com |
| REF11 | logP(ow) (MOE) | Chemical Computing Group | atomic contributions and correction factors | www.chemcomp.com |
| REF10 | h_logP (MOE) | Chemical Computing Group | QSPR, based on extended Hückel theory descriptors | www.chemcomp.com |
| REF12 | MoKa_logP | Molecular Discovery | QSPR, based on Molecular Interaction Field descriptors | www.moldiscovery.com |

**Table 3.**
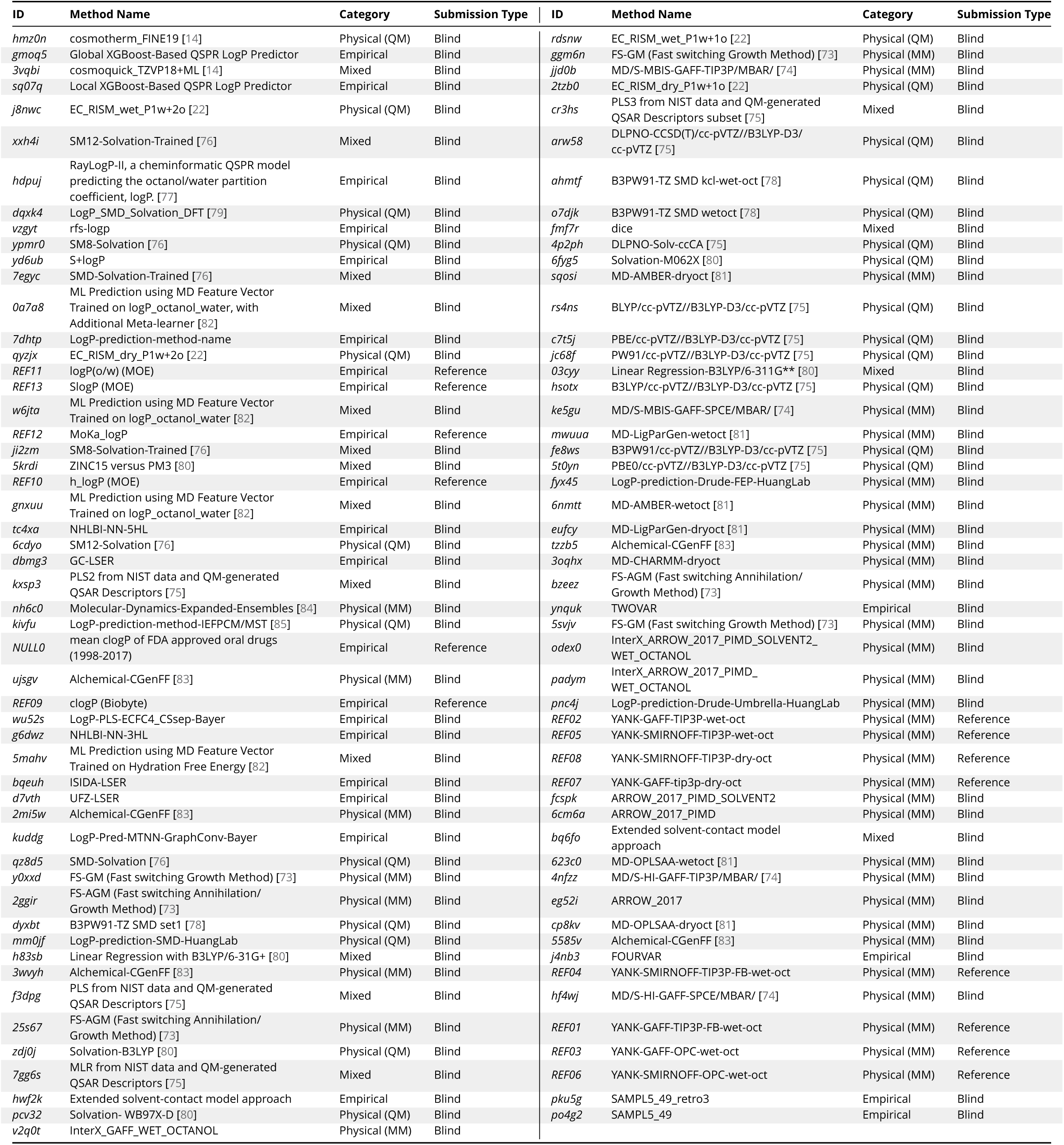
Submission IDs, names, category, and type for all the log *P* participant and reference calculation submissions. Submission IDs of methods are listed in the ID column. Reference calculations are labeled as *REF##*. The method name column lists the names provided by each participant in the submission file. The “type” column indicates if submission was or a post-deadline reference calculation, denoted by “Blind” or “Reference” respectively. The table is ordered by increasing RMSE from top to down and left to right, although many consecutively listed methods are statistically indistinguishable. All calculated error statistics are available in Tables S4 and S5.

### 2.2 Evaluation approach

A variety of error metrics were considered when analyzing predictions submitted to the SAMPL6 log *P* Challenge. Summary statistics were calculated for each submission for method comparison, as well as error metrics of predictions of each method. Both summary statistics and individual error analysis of predictions were provided to participants before the virtual workshop. Details of the analysis and scripts are maintained on the SAMPL6 Github Repository (described in section 6).

There are six error metrics reported: the root-mean-squared error (RMSE), mean absolute error (MAE), mean (signed) error (ME), coefficient of determination (R^2^), linear regression slope (m), and Kendall’s Rank Correlation Coefficient (*τ*). In addition to calculating these performance metrics, 95% confidence intervals were computed for these values using a bootstrapping-over-molecules procedure (with 10000 bootstrap samples) as described elsewhere in a previous SAMPL overview article [60]. Due to the small dynamic range of experimental log *P* values of the SAMPL6 set, it is more appropriate to use accuracy based performance metrics, such as RMSE and MAE, to evaluate methods than correlation-based statistics. This observation is also typically reflected in the confidence intervals on these metrics. Calculated errors statistics of all methods can be found in Tables S4 and S5.

Submissions were originally assigned to four method categories (physical, empirical, mixed, and other) by participants. However, when we evaluated the set of participating methods it became clear that it was going to be more informative to group them using the following categories: **physical (MM), physical (QM), empirical**, and **mixed**. Methods from the “other” group were reassigned to empirical or physical (QM) categories as appropriate. Methods submitted as Physical by participants included quantum mechanical (QM), molecular mechanics-based (MM) and, to a lesser extent, integral equation-based approaches (EC-RISM). We subdivided these submissions into “physical (MM)” and “physical (QM)” categories. Integral equation-based approaches were also evaluated under the Physical (QM) category. The “mixed” category includes methods that physical and empirical approaches are used in combination. Table 3 indicates the final category assignments in the “Category” column.

We created a shortlist of consistently well-performing methods that were ranked in the top 20 consistently according to two error and two correlation metrics: RMSE, MAE, R-Squared, and Kendall’s Tau. These are shown in Table 4.

**Table 4.**
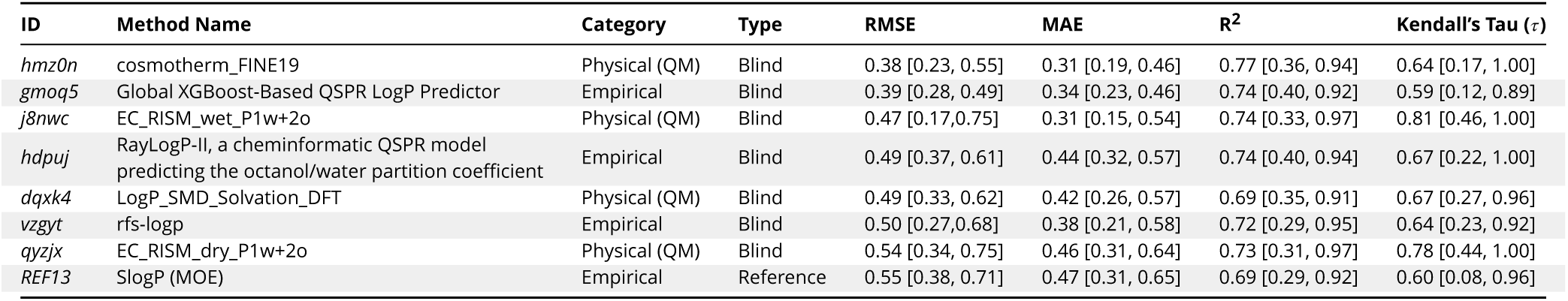
Eight consistently well-performing prediction methods based on consistent ranking within the Top 20 according to various statistical metrics. Submissions were ranked according to RMSE, MAE, R^2^, and *τ*. Many top methods were found to be statistically indistinguishable considering uncertainties of error metrics. Moreover, sorting of methods was influenced significantly by the choice of metric chosen. We assessed top 20 methods according the each metric to determine which methods are always among the top 20 according to all four statistical metrics chosen. A set of consistently well-performing methods were determined: Four QM-based and four empirical methods. Seven of these methods are blind submissions of SAMPL6 Challenge, and one of them (*REF13*) is a non-blind reference calculation. Performance statistics are provided as mean and 95% confidence intervals.

We included null and reference method prediction sets in the analysis to provide perspective for performance evaluations of blind predictions. Null models or null predictions employ a model which is not expected to be useful and can provide a simple point of comparison for more sophisticated methods, as ideally, such methods should improve on predictions from a null model. We created a null prediction set (submission ID *NULL0*) by predicting a constant log *P* value for every compound, based on a plausible log *P* value for drug-like compounds. We also provide reference calculations using several physical (alchemical) and empirical approaches as a point of comparison. The analysis is presented with and without the inclusion of reference calculations in the SAMPL6 GitHub repository. All figures and statistics tables in this manuscript include reference calculations. As the reference calculations were not formal submissions, these were omitted from formal ranking in the challenge, but we present plots in this article which show them for easy comparison. These are labeled with submission IDs of the form *REF##* to allow easy recognition of non-blind reference calculations.

In addition to the comparison of methods we also evaluated the relative difficulty of predicting log *P* of each molecule in the set. For this purpose, we plotted prediction error distributions of each molecule considering all prediction methods. We also calculated MAE for each molecule’s overall predictions as well as for predictions from each category as a whole.

## 3 Methods for reference calculations

Here we highlight the null prediction method and reference methods. We have included several widely-used physical and empirical methods as reference calculations in the comparative evaluation of log *P* prediction methods, in addition to the blind submissions of the SAMPL6 log *P* Challenge. These reference calculations are not formally part of the challenge but are provided as comparison methods. They were collected after the blind challenge deadline when the experimental data was released to the public. For a more detailed description of the methods used in the reference calculations, please refer to Section 12.1.

### 3.1 Physical Reference Calculations

Physical reference calculations were carried out using YANK [61], an alchemical free energy calculation toolkit [62, 63]. YANK implements Hamiltonian replica exchange molecular dynamics (H-REMD) simulations to sample multiple alchemical states and is able to explore a number of different alchemical intermediate functional forms using the OpenMM toolkit for molecular simulation [64–66].

The GAFF 1.81 [67] and SMIRNOFF (smirnoff99Frosst 1.0.7) [68] force fields were combined with three different water models. Water models are important for accuracy in modeling efforts in molecular modeling and simulation. The majority of modeling packages make use of rigid and fixed charge models due to their computational efficiency. To test how different water models can impact predictions, we combined three explicit water models TIP3P [69], TIP3P Force Balance (TIP3P-FB) [70] and the Optimal Point Charge (OPC) model [71] with the GAFF and SMIRNOFF force fields. The TIP3P and TIP3P-FB models are a part of the three-site water model class where each atom has partial atomic charges and a Lennard-Jones interaction site centered at the oxygen atom. The OPC model is a rigid 4-site, 3-charge water model that has the same molecular geometry as TIP3P, but the negative charge on the oxygen atom is located on a massless virtual site at the HOH angle bisector. This arrangement is meant to improve the water molecule’s electrostatic distribution. While TIP3P is one of the older and more common models used, OPC and TIP3P-FB are newer models that were parameterized to more accurately reproduce more of the physical properties of liquid bulk water.

Reference calculations also included wet and dry conditions for the octanol phase using the GAFF and SMIRNOFF force field with TIP3P water. The wet octanol phase was 27% water by mole fraction [58]. The methods used for physical reference calculations are summarized in Table 1.

Physical reference calculations (submission IDs: *REF01-REF08*) were done using a previously untested direct transfer free energy calculation protocol (DFE) which involved calculating the transfer free energy between water and octanol phases (explained in detail in Section 12.1.1), rather than a more typical protocol involving calculating a gas-to-solvent transfer free energy for each phase – an indirect solvation-based transfer free energy (IFE) protocol. In order to check for problems caused by this error, we included additional calculations performed by the more typical IFE protocol. Method details for the IFE protocol are presented in Section 12.1.2 and results are discussed in Section 4.2. However, only reference calculations performed with DFE protocol were included in overall evaluation of the SAMPL6 Challenge presented in Section 4.1, because only these spanned the full range of force fields and solvent models we sought to explore.

### 3.2 Empirical Reference Calculations

As empirical reference models, we used a number of commercial calculation programs, with the permission of the respective vendors, who agreed to have the results included in the SAMPL6 comparison. The programs are summarized in Table 2 and cover several of the different methodologies described in sections 1.2.2 and 12.1.3.

### 3.3 Our null prediction method

This submission set is designed as a null model which predicts the log *P* of all molecules to be equal to the mean clogP of FDA approved oral new chemical entities (NCEs) between the years 1998 and 2017 based on the analysis of Micheal D. Shultz (2019) [72]. We show this null model with submission ID *NULL0*. The mean clogP of FDA approved oral NCEs approved between 1900-1997, 1998-2007, and 2008-2017 were reported 2.1, 2.4, and 2.9, respectively, using StarDrop clogP calculations (https://www.optibrium.com/). We calculated the mean of NCEs approved between 1998 – 2017, which is 2.66, to represent the average log *P* of contemporary drug-like molecules. We excluded the years 1900-1997 from this calculation as the early drugs tend to be much smaller and much more hydrophilic than the ones being developed at present.

## 4 Results and Discussion

### 4.1 Overview of challenge results

A large variety of methods were represented in the SAMPL6 log *P* Challenge. There were 91 blind submissions collected from 27 participating groups in the log *P* challenge (Tables of participants and the predictions they submitted are presented in SAMPL6 GitHub Repository and its archived copy in the Supporting Information.) This represented an increase in interest over the previous SAMPL challenges. In the SAMPL5 Cyclohexane-Water log *D* Challenge, there were 76 submissions from 18 participating groups [2], so participation was even higher this iteration.

Out of blind submissions of the SAMPL6 log *P* Challenge, there were 31 in the physical (MM) category, 25 in the physical (QM) category, 18 in the empirical category, and 17 in the mixed method category (Table 3). We also provided additional reference calculations – five in the empirical category, and eight in the physical (MM) category.

The following sections present detailed performance evaluation of blind submissions and reference prediction methods. Performance statistics of all the methods can be found in S4. Methods are referred to by their submission ID’s which are provided in 3.

#### 4.1.1 Performance statistics for method comparison

Many methods in the SAMPL6 Challenge achieved good predictive accuracy for octanol-water log *P* values. Figure 3 shows the performance comparison of methods based on accuracy with RMSE and MAE. 10 methods achieved an RMSE ≤ 0.5 log *P* units. These methods were QM-based, empirical, and mixed approaches (submission IDs: *hmz0n, gmoq5, 3vqbi, sq07q, j8nwc, xxh4i, hdpuj, dqxk4, vzgyt, ypmr0*). Many of the methods had an RMSE ≤ 1.0 log *P* units. These 40 methods include 34 blind predictions, 5 reference calculations, and the null prediction method.

**Figure 1.**
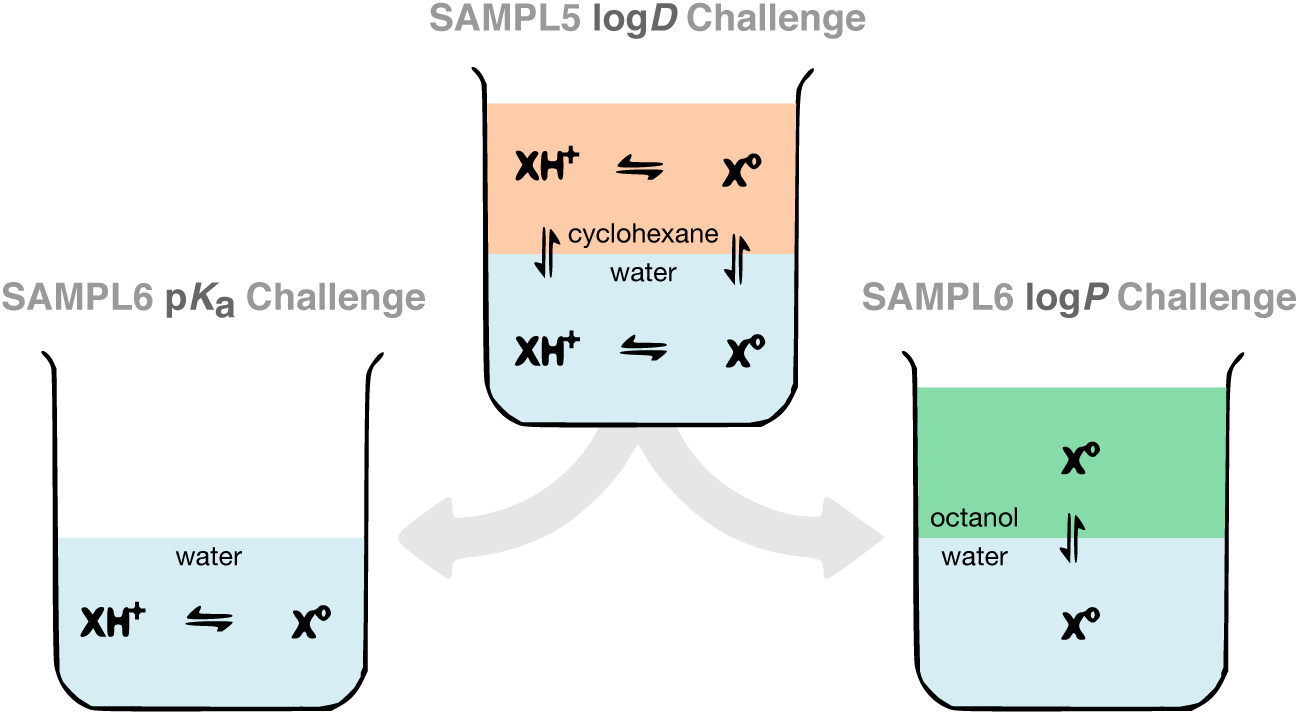
The desire to deconvolute the distinct sources of error contributing to the large errors observed in the SAMPL5 log *D* challenge motivated the separation of p*K*_a_ and log *P* challenges in SAMPL6. The SAMPL6 p*K*_a_ and log *P* challenges aim to evaluate protonation state predictions of small molecules in water and transfer free energy predictions between two solvents, isolating these prediction problems.

**Figure 2.**
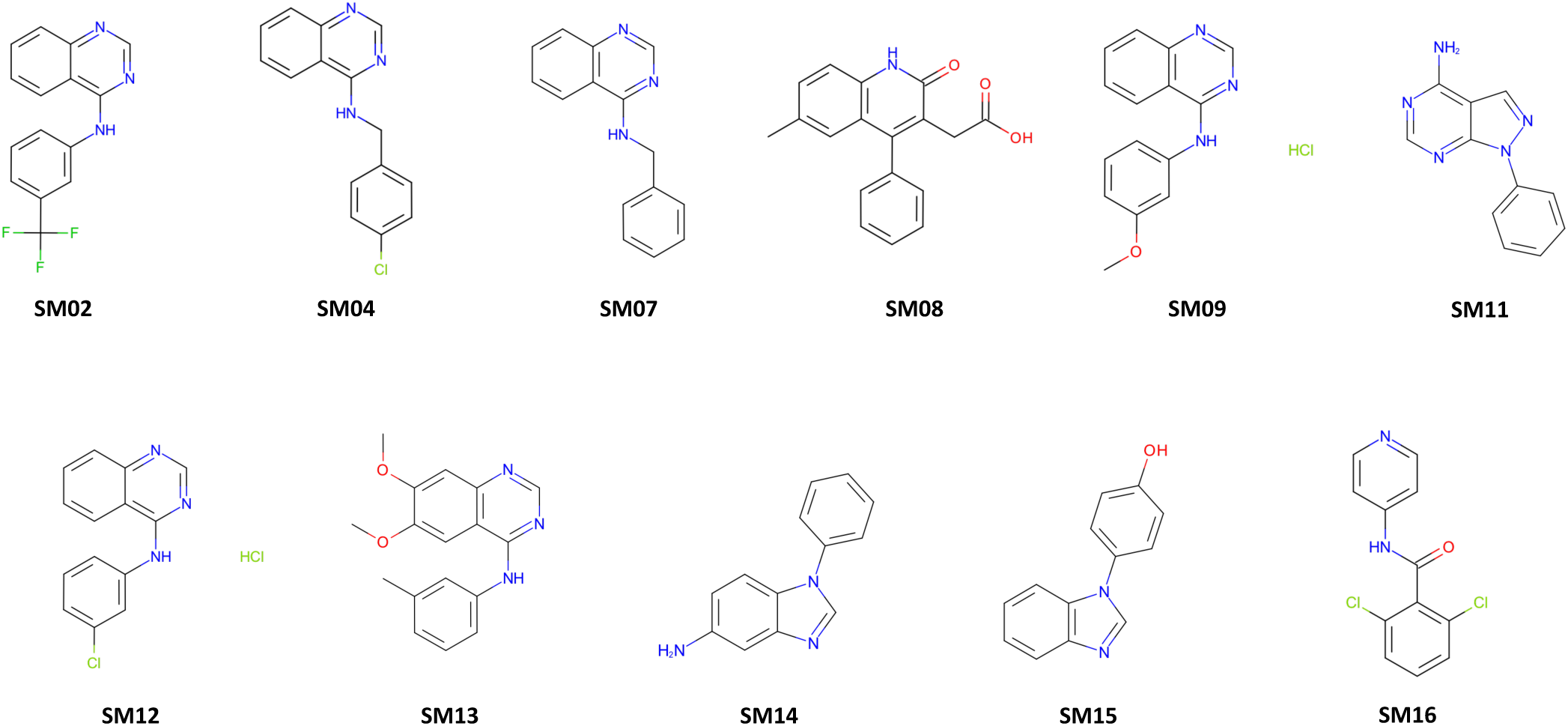
Structures of the 11 protein kinase inhibitor fragments used for the SAMPL6 log *P* Blind Prediction Challenge. These compounds are a subset of the SAMPL6 p*K*_a_ Challenge compound set [8] which were found to be tractable potentiometric measurements with sufficient solubility and p*K*_a_ values far from pH titration limits. Chemical identifiers of these molecules are available in Table S2 and experimental log *P* values are published [9]. Molecular structures in the figure were generated using OEDepict Toolkit [53].

**Figure 3.**
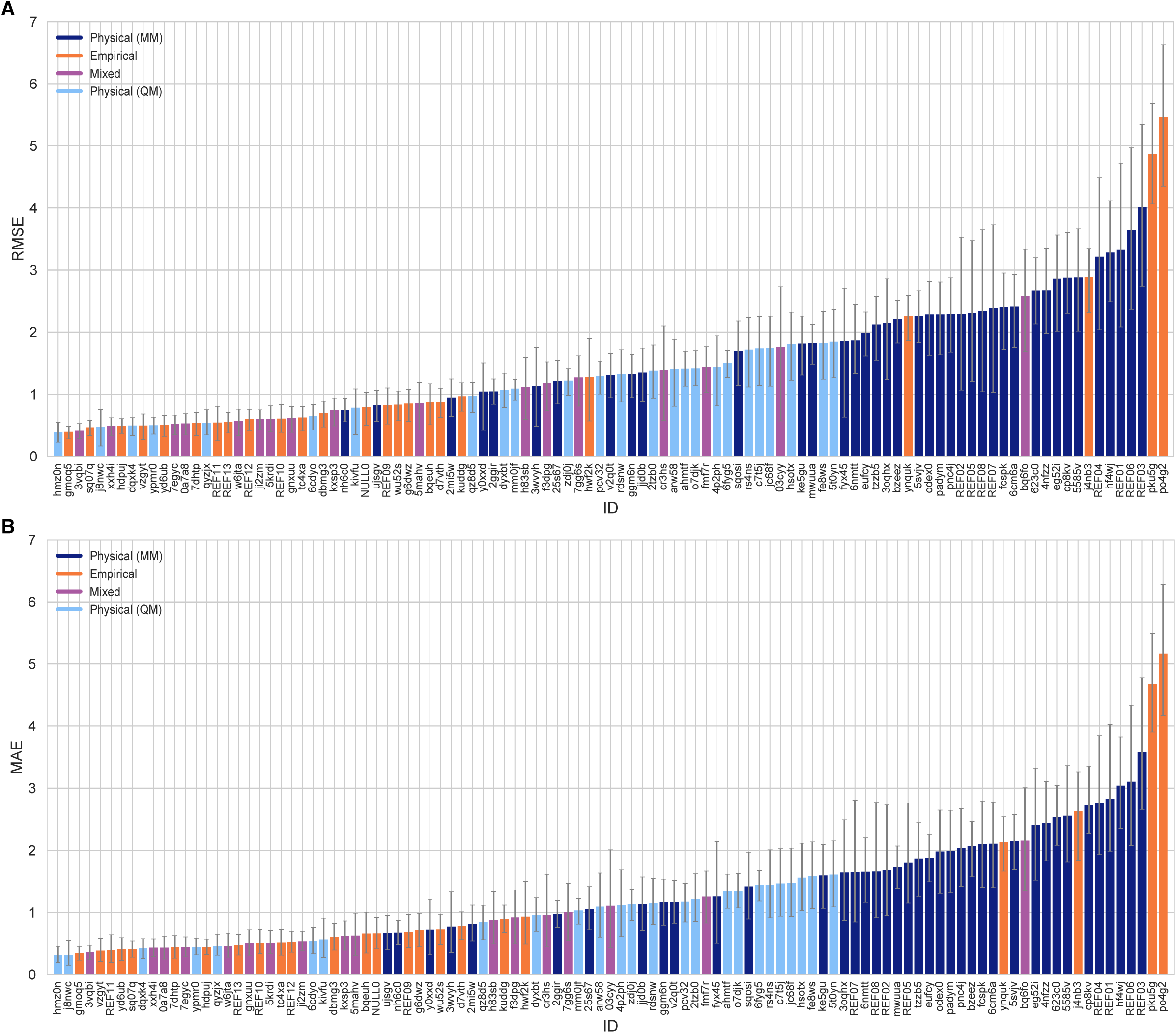
Overall accuracy assessment for all methods participating in the SAMPL6 log *P* Challenge. Both root-mean squared error (RMSE) and mean absolute error (MAE) are shown, with error bars denoting 95% confidence intervals obtained by bootstrapping over challenge molecules. Submission IDs are summarized in Table 3. Submission IDs of the form *REF##* refer to non-blinded reference methods computed after the blind challenge submission deadline, and *NULL0* is the null prediction method; all others refer to blind, prospective predictions.

Correlation-based statistics methods only provide a rough comparison of methods of the SAMPL6 Challenge, given the small dynamic range of the experimental log *P* dataset. Figure 4 shows R^2^ and Kendall’s Tau values calculated for each method, sorted from high to low performance. However, the uncertainty of each correlation statistic is quite high, not allowing a true ranking based on correlation. Methods with R^2^ and Kendall’s Tau higher than 0.5 constitute around 50% of the methods and can be considered as the better half. However, the performance of individual methods is statistically indistinguishable. Nevertheless, it is worth noting that QM-based methods appeared marginally better at capturing the correlation and ranking of experimental log *P* values. These methods comprised the top four based on R^2^ (≥ 0.75; submission IDs: *2tzb0, rdsnw, hmz0n, mm0jf*), and the top six based on Kendall’s Tau, (≥ 0.70; submission IDs: *j8nwc, qyzjx, 2tzb0, rdsnw, mm0jf*, and *6fyg5*). However, due to the small dynamic range and the number of experimental log *P* values of the SAMPL6 set, correlation-based statistics are less informative than accuracy-based performance metrics such as RMSE and MAE.

**Figure 4.**
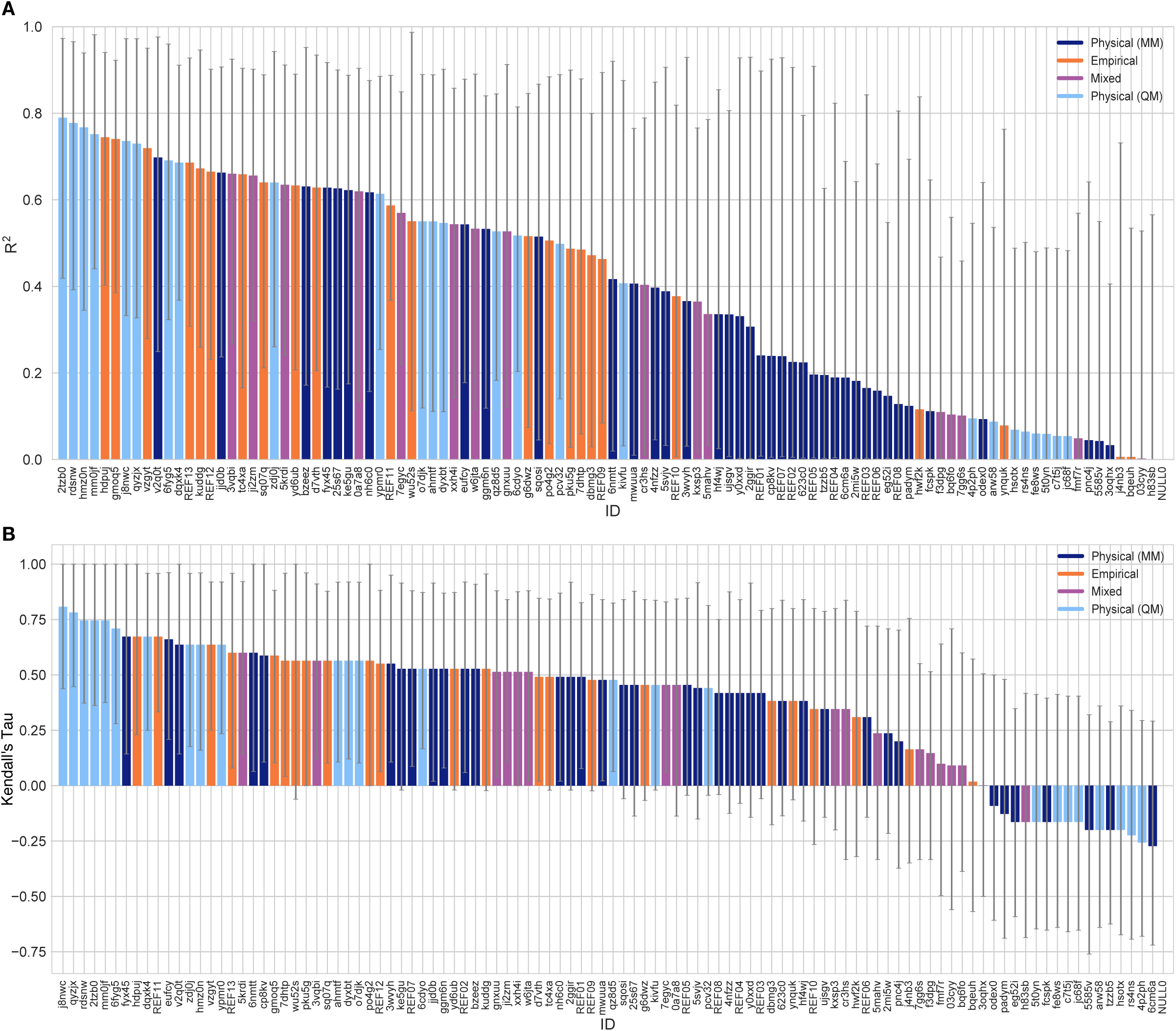
Overall correlation assessment for all methods participating SAMPL6 log *P* Challenge. Pearson’s R^2^ and Kendall’s Rank Correlation Coefficient Tau (*τ*) are shown, with error bars denoting 95% confidence intervals obtained by bootstrapping over challenge molecules. Submission IDs are summarized in Table 3. Submission IDs of the form *REF##* refer to non-blinded reference methods computed after the blind challenge submission deadline, and *NULL0* is the null prediction method; all others refer to blind, prospective predictions. Overall, a large number and wide variety of methods have a statistically indistinguishable performance on ranking, in part because of the relatively small dynamic range of this set and because of the small size of the set. Roughly the top half of methods with Kendall’s Tau > 0.5 fall into this category.

#### 4.1.2 Results from physical methods

One of the aims of the SAMPL6 log *P* Challenge was to assess the accuracy of physical approaches in order to potentially provide direction for improvements which could later impact accuracy in downstream applications like protein-ligand binding. Some MM-based methods used for log *P* predictions use the same technology applied to protein-ligand binding predictions, so improvements made to modeling of partition may in principle carry over. However, prediction of partition between two solvent phases is a far simpler test only capturing some aspects of affinity prediction – specifically, small molecule and solvation modeling – in the absence of protein-ligand interactions and protonation state prediction problems.

Figure 5 shows a comparison of the performance of MM- and QM-based methods in terms of RMSE and Kendall’s Tau. Both in terms of accuracy and ranking ability, QM methods resulted in better results, on average. QM methods using implicit solvation models outperformed MM-based methods with explicit solvent methods that were expected to capture the heterogeneous structure of the wet octanol phase better. Only 3 MM-based methods and 8 QM-based methods achieved RMSE less than 1 log *P* unit. 5 of these QM-based methods showed very high accuracy (RMSE ≤ 0.5 log *P* units). The three MM-based methods with the lowest RMSE were:

**Figure 5.**
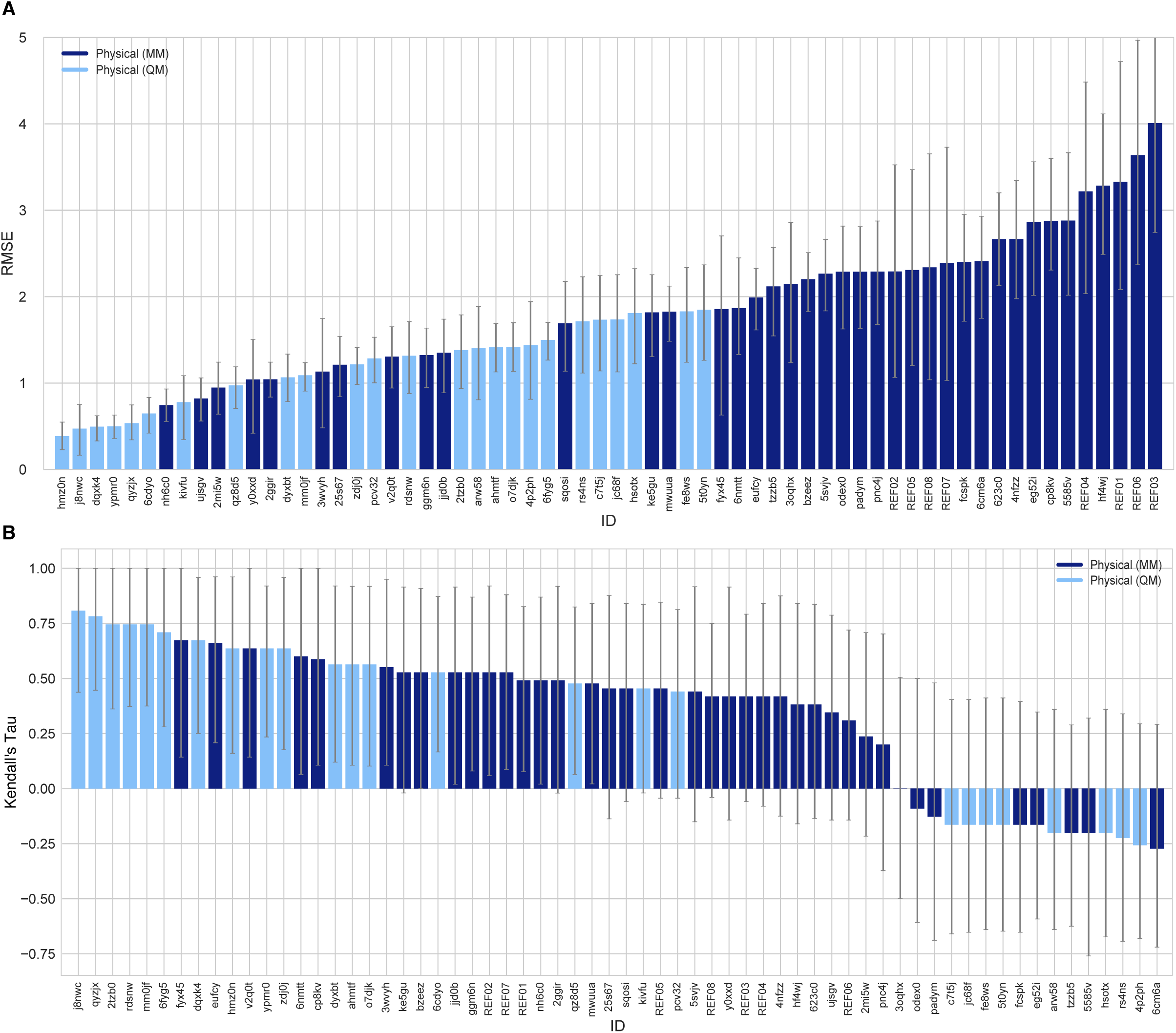
Performance statistics of physical methods. Physical methods are further classified into quantum chemical (QM) methods and molecular mechanics (MM) methods. RMSE and Kendall’s Rank Correlation Coefficient Tau are shown, with error bars denoting 95% confidence intervals obtained by bootstrapping over challenge molecules. Submission IDs are summarized in Table 3. Submission IDs of the form *REF##* refer to non-blinded reference methods computed after the blind challenge submission deadline; all others refer to blind, prospective predictions.

- Molecular-Dynamics-Expanded-Ensembles (*nh6c0*): This submission used an AMBER/OPLS-based force field with manually adjusted parameters (following rules from the participant’s article [86]), modified Toukan-Rahman water model with Non-zero Lennard-Jones parameters [84], and modified Expanded Ensembles (EEMD) method [87] for free energy estimations.
- Alchemical-CGenFF (*ujsgv, 2mi5w*, [83]): These two submissions used Multi-Phase Boltzmann Weighting with the CHARMM Generalized Force Field (CGenFF) [88], and the TIP3P water model [69]. From the brief method descriptions submitted to the challenge we could not identify the difference between these prediction sets.

RMSE values for predictions made with MM-based methods ranged from 0.74 to 4.00 log *P* units, with the average of the better half being 1.44 log *P* units.

Submissions included diverse molecular simulation-based log *P* predictions made using alchemical approaches. These included Free Energy Perturbation (FEP) [89] and BAR estimation [90], Thermodynamic integration (TI) [91], and non-equilibrium switching (NES) [92, 93]. Predictions using YANK [61] Hamiltonian replica exchange molecular dynamics and MBAR [94] were provided as reference calculations.

A variety of combinations of force fields and water models were represented in the challenge. These included CGenFF with TIP3P or OPC3 [95] water models; OPLS-AA [96] with OPC3 and TIP4P [69] water models; GAFF [67] with TIP3P, TIP3P Force Balance [70], OPC [71], and OPC3 water models; GAFF2 [97] with the OPC3 water model; GAFF with Hirshfeld-I [98] and Minimal Basis Set Iterative Stockholder(MBIS) [99] partial charges and the TIP3P or SPCE water models [100]; the SMIRNOFF force field [68] with the TIP3P, TIP3P Force Balance, and OPC water models; and submissions using Drude [101] and ARROW [102] polarizable force fields.

Predictions that used polarizable force fields did not show an advantage over fixed-charged force fields in this challenge. RMSEs for polarizable force field submissions range from 1.85 to 2.86 (submissions with the Drude Force Field were *fyx45, pnc4j*, and those with the ARROW Force Field were *odex0, padym fcspk*, and *6cm6a*).

Predictions using both dry and wet octanol phases were submitted to the log *P* challenge. When submissions from the same participants were compared, we find that including water in the octanol phase only slightly lowers the RMSE (0.05-0.10 log *P* units), as seen in Alchemical-CGenFF predictions (wet: *ujshv, 2mi5w, ttzb5*; dry: *3wvyh*), YANK-GAFF-TIP3P predictions (wet: *REF02*, dry: *REF07*), MD-LigParGen predictions with OPLS and TIP4P (wet: *mwuua*, dry: *eufcy*), and MD-OPLSAA predictions with TIP4P (wet: *623c0*, dry: *cp8kv*). However this improvement in performance with wet octanol phase was not found to be a significant effect on overall prediction accuracy. Methodological differences and choice of force field have a greater impact on prediction accuracy than the composition of the octanol phase.

Refer to Table S1 for a summary of force fields and water models used in MM-based submissions. For additional analysis, we refer the interested reader to the work of Piero Procacci and Guido Guarnieri, who provide a detailed comparison of MM-based alchemical equilibrium and non-equilibrium approaches in SAMPL6 Challenge in their paper [73]. Specifically, in the section “Overview on MD-based SAMPL6 submissions” of their paper, they provide comparisons subdividing submissions based force field (for CGenFF, GAFF1/2, and OPLS-AA).

#### 4.1.3 A shortlist of consistently well-performing methods

Although there was not any single method that performed significantly better than others in the log *P* challenge, we identified a group of consistently well performing methods. There were many methods with good performance when judged based on RMSE, but not many methods consistently showed up at the top according to all metrics. When individual error metrics are considered, many submissions were not different from one another in a statistically significant way, and ranking typically depends on the metric chosen due overlapping confidence intervals. Instead, we identified several consistently well performing methods by looking at several different metrics – two assessing accuracy (RMSE and MAE) and two assessing correlation (Kendall’s Tau and R^2^). We determined those methods which are in the top 20 by each of these metrics. This resulted in a list of eight methods which are consistently well performing. The shortlist of consistently well-performing methods are presented in Table 4.

The resulting eight consistently well-performing methods were QM-based physical models and empirical methods. These eight methods were fairly diverse. Traditional QM-based physical methods included log *P* predictions with COSMO-RS method as implemented in COSMOtherm v19 at the BP//TZVPD//FINE Single Point level (*hmz0n*, [16–18]) and the SMD solvation model with the M06 density functional family (*dqxk4*, [79]). Additionally, two other top QM-based methods seen in this shortlist used EC-RISM theory with wet or dry octanol (*j8nwc* and *qyzjx*) [22]. Several empirical submissions also were among these well-performing methods – specifically, the Global XGBoost-Based QSPR LogP Predictor (*gmoq5*), the RayLogP-II (*hdpuj*) approach, and rfs-logp (*vzgyt*). Among reference calculations, SlogP calculated by MOE software (*REF13*) was the only method that was consistently well performing.

Figure 6 compares log *P* predictions with experimental values for these 8 well-performing methods, as well as one additional method which has an average level of performance. This representative method (*rdsnw*, [22]) is the method with the highest RMSE below the median of all methods (including reference methods).

**Figure 6.**
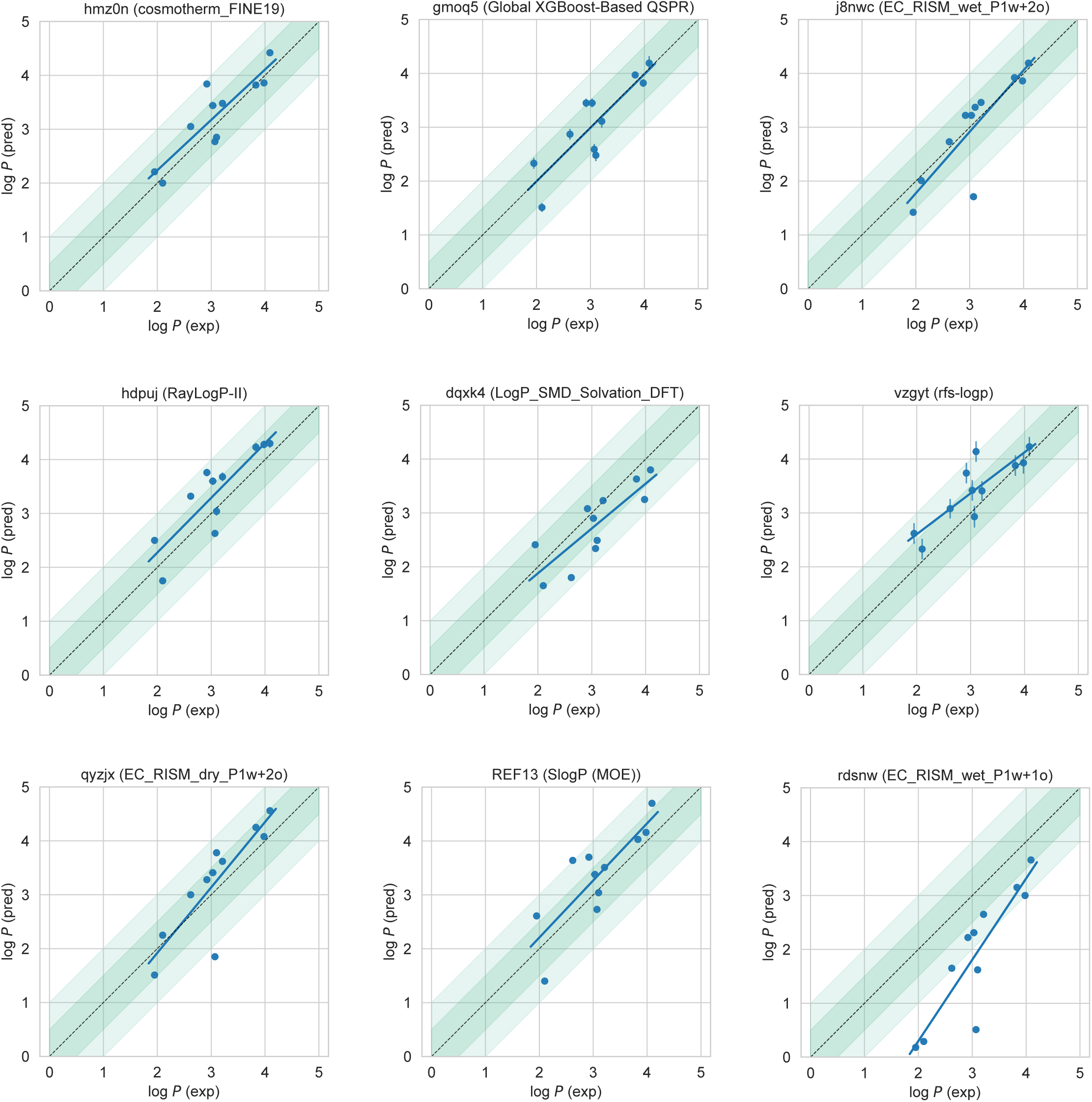
Predicted vs experimental value correlation plots of 8 best-performing methods and one representative average method. Dark and light green shaded areas indicate 0.5 and 1.0 units of error. Error bars indicate standard error of the mean of predicted and experimental values. Experimental log *P* SEM values are too small to be seen under the data points. EC_RISM_wet_P1w+1o method (*rdsnw*) was selected as the representative average method, as it is the method with the highest RMSE below the median.

#### 4.1.4 Difficult chemical properties for log *P* predictions

In addition to comparing method performance, we analyzed the prediction errors for each compound in the challenge set to assess whether particular compounds or chemistries are especially challenging (Figure 7). For this analysis, MAE is a more appropriate statistical value for following global trends, as its value is less affected by outliers than is RMSE.

**Figure 7.**
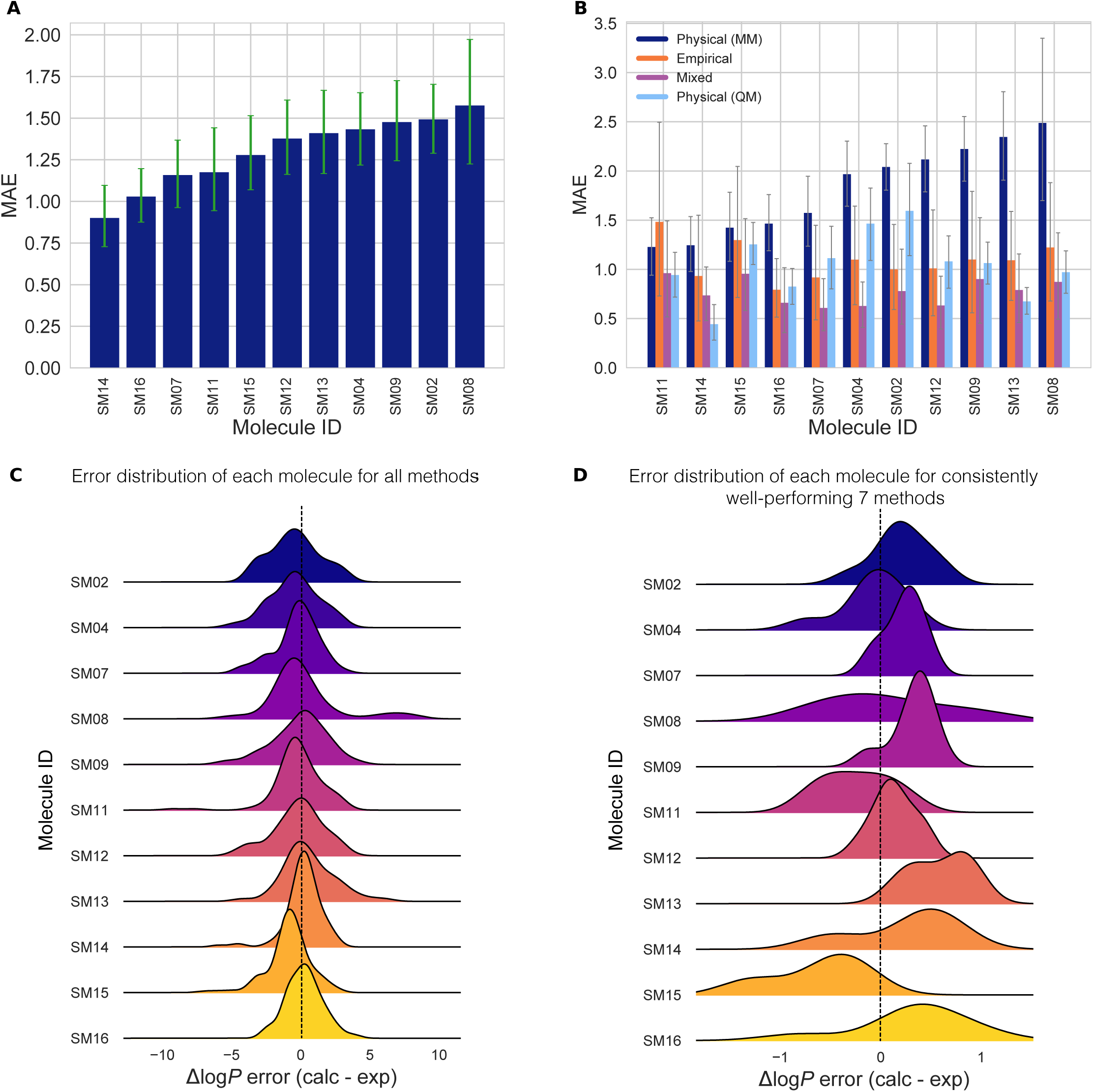
Molecule-wise prediction error distribution plots show how variable the prediction accuracy was for individual molecules across all prediction methods. **(A)** MAE calculated for each molecule as an average of all methods shows relatively uniform MAE across the challenge set. SM14 and SM16 predictions were slightly more accurate than the rest. **(B)** MAE of each molecule broken out by method category shows that for each method category the most challenging molecules were different. Predictions of SM08, SM13, SM09, and SM12 log *P* values were significantly less accurate with Physical (MM) methods than the other method categories. For QM-based methods SM04 and SM02 were most challenging. Largest MAE for Empirical methods were observed for SM11 and SM15. **(C)** Error distribution for each SAMPL6 molecule overall prediction methods. It is interesting to note that most distributions are peaked near an error of zero, suggesting that perhaps a consensus model might outperform most individual models. However, SM15 is more significantly shifted away from zero than any other compound. SM08 has a significant tail showing probability of overestimated log *P* predictions by some methods. **(D)** Error distribution for each molecule calculated for only 7 methods from blind submissions that were determined to be consistently well-performing (*hmz0n, gmoq5, j8nwc, hdpuj, dqxk4, vzgyt, qyzjx*).

Performance on individual molecules shows relatively uniform MAE across the challenge set (Figure 7A). Predictions of SM14 and SM16 were slightly more accurate than the rest of the molecules when averaged across all methods. Prediction accuracy on each molecule, however, is highly variable depending on method category (Figure 7B). Predictions of SM08, SM13, SM09, and SM12 were significantly less accurate with physical (MM) methods than the other method categories by 2 log *P* units in terms of MAE over all methods in each category. These molecules were not challenging for QM-based methods. Discrepancies in predictions of SM08 and SM13 are discussed in Section 4.2. For QM-based methods, SM04 and SM02 were most challenging. The largest MAE for empirical methods was observed for SM11 and SM15.

Figure 7C shows the error distribution for each SAMPL6 molecule over all prediction methods. It is interesting to note that most distributions are peaked near an error of zero, suggesting that perhaps a consensus model might outperform most individual models. However, SM15 is more significantly shifted away from zero than any other compound (ME calculated accross all molecules is −0.88±1.49 for SM15). SM08 had the most spread in log *P* prediction error.

This challenge focused on log *P* of neutral species, rather than log *D* as studied in SAMPL5, which meant that we do not see the same trends where performance is significantly worse for compounds with multiple protonation states/tautomers or where p*K* _a_ values are uncertain. However, in principle, tautomerization can still influence log *P* values. Multiple neutral tautomers can be present at significant populations in either solvent phase, or the major tautomer can be different in each solvent phase. However, this was not expected to be the case for any of the 11 compounds in this SAMPL6 Challenge. We do not have experimental data on the identity or ratio of tautomers, but tautomers other than those depicted in Figure 2 would be much higher in energy according to QM predictions [22] and, thus, very unlikely to play a significant role. Still, for most log *P* prediction methods, it was at least necessary for participants to select the major neutral tautomer. We do not observe statistically worse error for compounds with potential tautomer uncertainties here, suggesting it was not a major factor in overall accuracy, some participants *did* chose to run calculations on tautomers that were not provided in the challenge input files (Figure 11 and Table 5), as we discuss in Section 4.2.

**Figure 8.**
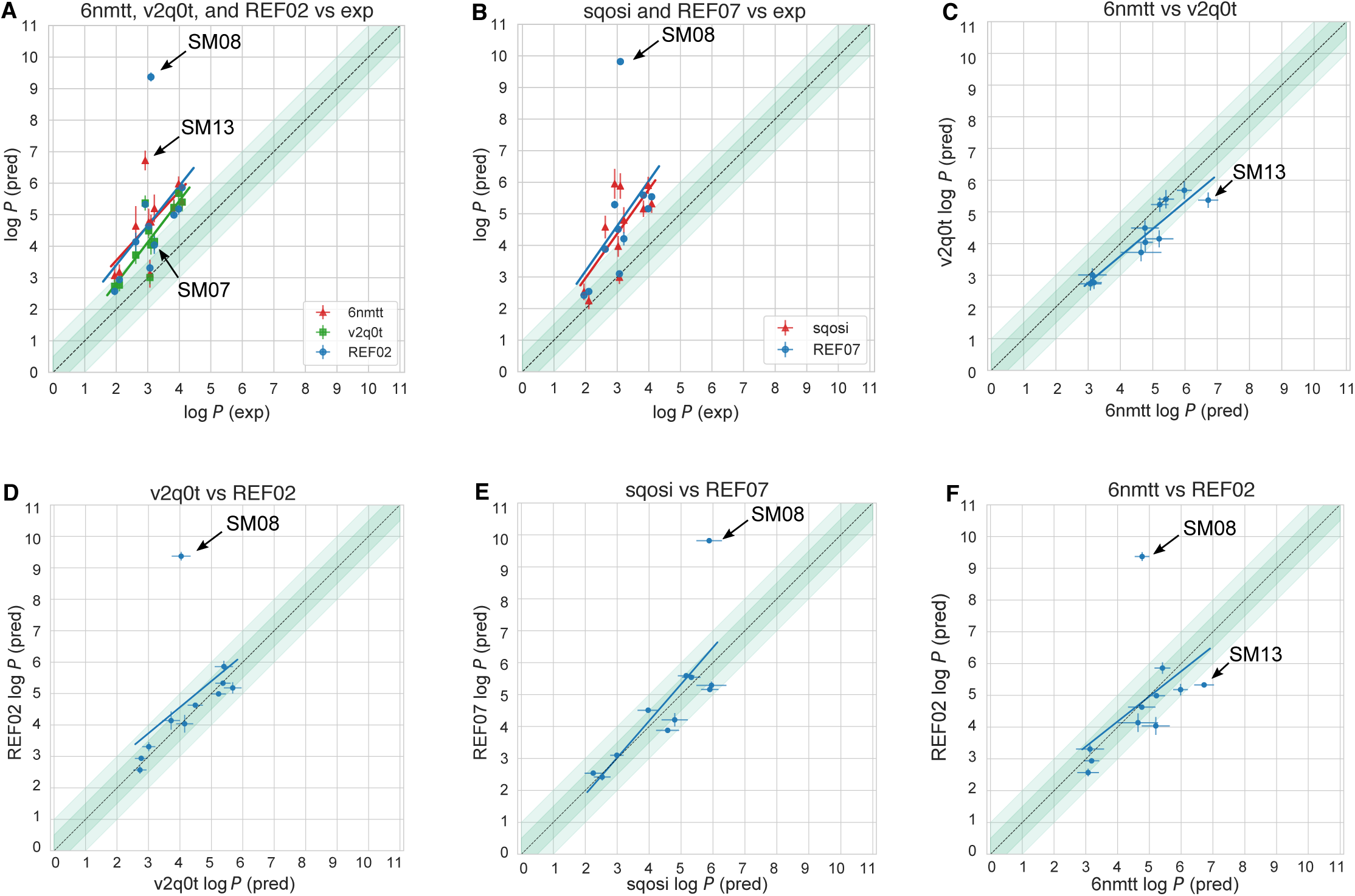
Comparison of independent predictions that use seemingly identical methods (free energy calculations using GAFF and TIP3P water) shows significant systematic deviations between predictions for many compounds. Comparison of the calculated and experimental values for submissions *v2q0t* (InterX_GAFF_WET_OCTANOL), *6nmtt* (MD-AMBER-wetoct), *sqosi* (MD-AMBER-dryoct) and physical reference calculations *REF02* (YANK-GAFF-TIP3P-wet-oct) and *REF07* (YANK-GAFF-TIP3P-dry-oct). (A) compares calculations that used wet octanol, and (B) compares those that used dry octanol. Plots C to F show the methods compared to one another. The dark and light-shaded region indicates 0.5 and 1.0 units of error, respectively.

**Figure 9.**
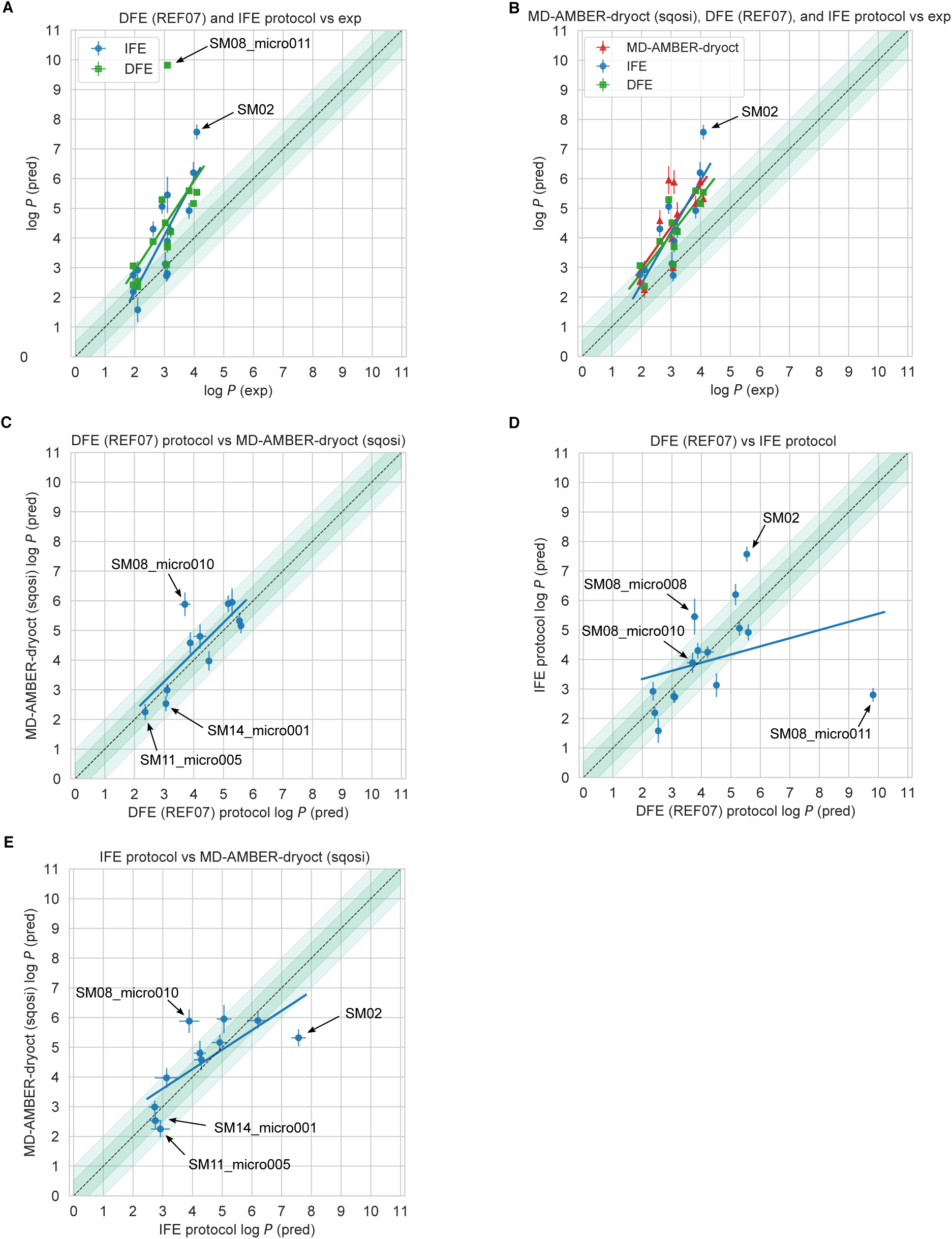
Comparison of predictions that use free energy calculations using GAFF and TIP3P water show deviations between predictions for the challenge molecules and several alternative tautomers and resonance structures. Deviations seem to largely stem from differences in equilibration amount and choice of tautomer. **A** compares reference direct transfer free energy (DFE, *REF07*) and indirect solvation-based transfer free energy (IFE) protocols to experiment for the challenge provided resonance states of molecules and a couple of extra resonance states for SM14 and SM11, and extra tautomers for SM08. **B** compares the same exact tautomers for submission *sqosi* (MD-AMBER-dryoct) and the two reference protocols to experiment. Submission *sqosi* (MD-AMBER-dryoct) used different tautomers than the ones provided in the challenge. **C-E** compares the calculated log *P* between different methods using the same tautomers. All of the predicted values can be found in Table 5.

**Figure 10.**
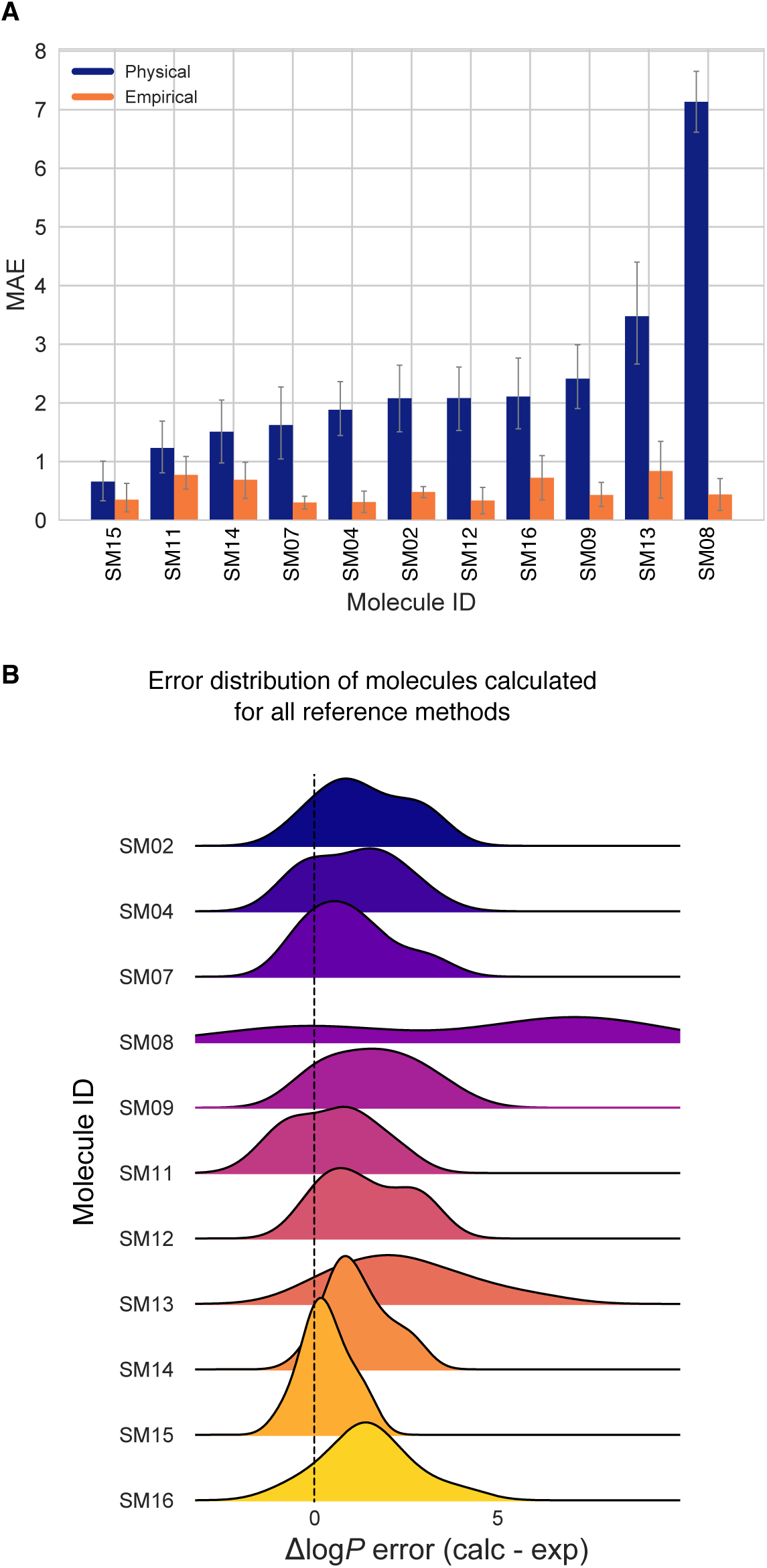
The prediction errors per molecule indicate some compounds were more difficult to predict than others for the reference calculations category. (A) MAE of each SAMPL6 molecule broken out by physical and empirical reference method category. (B) Error distribution for each molecule calculated for the reference methods. SM08 was the most difficult to predict for the physical reference calculations, due to our partial charge assignment procedure.

**Figure 11.**
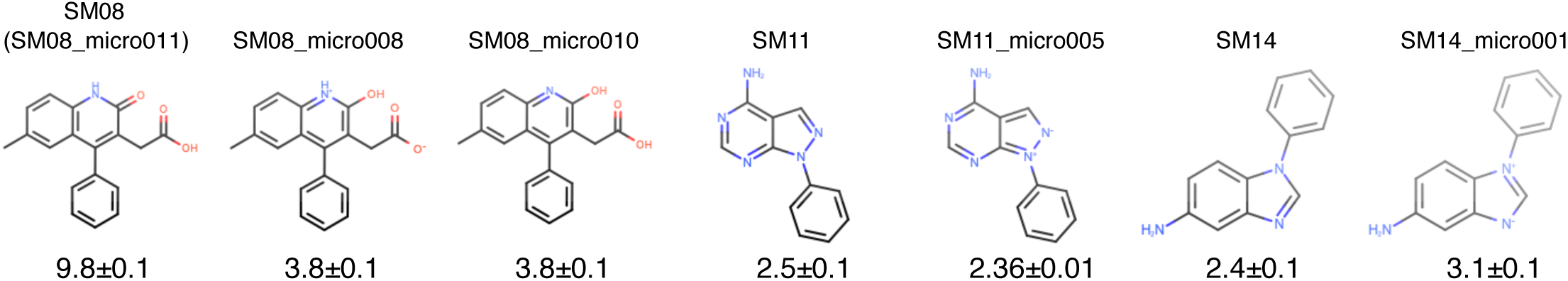
The tautomer and resonance structure choice resulted in discrepancies in the reference calculations. Shown here are calculated values for different input structures using the reference direct transfer free energy method. The uncertainties of the log *P* predictions were calculated as the standard error of the mean (SEM) of three replicate predictions. Structures labelled as SM08, SM11, and SM14 are based on input SMILES provided in SAMPL6 log *P* Challenge instructions. Three microstates shown for SM08 are different tautomers. SM08 (SM08_micro011) and SM08_micro010 are carboxylic acids, while SM08_micro008 is a carboxylate ion. SM08 (SM08_micro011) has a carbonyl group in the ring, while SM08_micro008 and SM08_micro010 have a hydroxyl in the ring. Structures pertaining to SM11 and SM14 are different resonance hybrids of the same tautomer (neutral microstate). Enumeration of all theoretically possible neutral tautomers of SAMPL6 molecules can be found in the SAMPL6 GitHub Repository (https://github.com/samplchallenges/SAMPL6/tree/master/physical_properties/pKa/microstates).

**Table 5.** Predicted log *P* values of free energy calculations of methods using GAFF, TIP3P water, and dry octanol. The methods listed are the reference direct transfer free energy (DFE) protocol, reference indirect solvation-based transfer free energy (IFE) protocol and submission *sqosi* (MD-AMBER-dryoct). Details of the two reference protocols can be found in Section 12.1. log *P* predictions for multiple tautomers (SM08) and resonance structures (SM11 and SM14) are listed, when available. The experimental values are provided for comparison. The same experimental log *P* values are stated for multiple tautomers or resonance structures. Potentiometric log *P* measurements do not provide information about the identity or populations of tautomers.

| Molecule | Indirect Solvation-Based Transfer Free Energy (IFE) Protocol | REF07 Direct Transfer Free Energy (DFE) Protocol | sqosi (MD-AMBER-dryoct) | Experimental |
| --- | --- | --- | --- | --- |
| SM02 | 7.6±0.3 | 5.5±0.1 | 5.3±0.3 | 4.09±0.03 |
| SM04 | 6.2±0.4 | 5.2±0.1 | 5.9±0.3 | 3.98±0.03 |
| SM07 | 4.3±0.2 | 4.2±0.3 | 4.8±0.4 | 3.21±0.04 |
| SM08<sup>1,2</sup> | 2.8±0.2 | 9.8±0.1 | - | 3.10±0.03 |
| SM08_micro008 | 5.5±0.6 | 3.8±0.1 | - | 3.10±0.03 |
| SM08_micro010 | 3.9±0.3 | 3.7±0.2 | 5.9±0.4 | 3.10±0.03 |
| SM09 | 3.1±0.4 | 4.51±0.03 | 4.0±0.3 | 3.0±0.1 |
| SM11<sup>1</sup> | 1.6±0.4 | 2.5±0.1 | - | 2.10±0.04 |
| SM11_micro005 | 2.9±0.3 | 2.36±0.01 | 2.3±0.3 | 2.10±0.04 |
| SM12 | 4.9±0.3 | 5.6±0.1 | 5.2±0.3 | 3.83±0.03 |
| SM13 | 5.1±0.3 | 5.3±0.1 | 6.0±0.5 | 2.92±0.04 |
| SM14<sup>1</sup> | 2.2±0.2 | 2.4±0.1 | - | 1.95±0.03 |
| SM14_micro001 | 2.8±0.2 | 3.1±0.1 | 2.5±0.3 | 1.95±0.03 |
| SM15 | 2.7±0.2 | 3.1±0.1 | 3.0±0.2 | 3.07±0.03 |
| SM16 | 4.3±0.3 | 3.9±0.1 | 4.6±0.4 | 2.62±0.01 |

<sup>1</sup> The tautomer or resonance structure presented as the input SMILES for the SAMPL6 log *P* Challenge.
<sup>2</sup> It corresponds to the microstate SM08\_micro011 of the SAMPL6 pK<sub>a</sub> Challenge.

#### 4.1.5 Comparison to the past SAMPL challenges

Overall, SAMPL6 log *P* predictions were more accurate than log *D* predictions in the SAMPL5 Cyclohexane-Water log *D* Challenge (Figure 3). In the log *D* challenge, only five submissions had an RMSE ≤ 2.5 log units, with the best having an RMSE of 2.1 log *P* units. A rough estimate of expected error for log *P* and log *D* is 1.54 log units. This comes from taking the mean RMSE of the top half of submissions in SAMPL4 Hydration Free Energy Prediction Challenge (1.5 kcal/mol) [60] and assuming the error in each phase is independent and equal to this value, yieding an expected error of 1.54 log *P* units [2]. Here, 64 log *P* challenge methods performed better than this threshold (58 blind predictions, 5 reference calculations, and the null prediction). However, only 10 of them were MM-based methods, with the lowest RMSE of 0.74 observed for method named Molecular-Dynamics-Expanded-Ensembles (*nh6c0*).

Challenge construction and experimental data availability are factors that contributed to the higher prediction accuracy observed in SAMPL6 compared to prior years. The log *P* challenge benefited from having a well-defined protonation state, especially for physical methods. Empirical methods benefited from the wealth of octanol-water training data. Accordingly, empirical methods were among the best performers here. But also, the chemical diversity represented by 11 compounds of the SAMPL6 log *P* challenge is very restricted and lower than the 53 small molecules in the SAMPL5 log *D* Challenge set. This was somewhat consistent with our expectations (discussed in Section 1.2.3)—that empirical, QM (with trained implicit solvation models), and mixed methods would outperform MM methods given their more extensive use of abundant octanol-water data in training (Figure 3).

### 4.2 Lessons learned from physical reference calculations

#### 4.2.1 Comparison of reference calculations did not indicate a single force field or water model with dramatically better performance

As in previous SAMPL challenges, we conducted a number of reference calculations with established methods to provide a point of comparison. These included calculations with alchemical physical methods. Particularly, to see how the choice of water model affects accuracy we included three explicit solvent water models – TIP3P, TIP3P-FB and the OPC model – with the GAFF and SMIRNOFF force fields in our physical reference calculations. Deviations from experiment were significant (RMSE values ranged from 2.3 [1.1, 3.5] to 4.0 [2.7, 5.3] log units) across all the conditions used in the physical reference predictions (Figure 3A). In general, all the water models tend to overestimate the log *P*, especially for the carboxylic acid in the challenge set, SM08, though our calculations on this molecule had some specific difficulties we discuss further below. Relative to the TIP3P-FB and OPC water models, predictions which used TIP3P showed improvement in some of the error metrics, such as lower deviation from experiment with an RMSE range of 2.3 [1.1, 3.5] to 2.34 [1.0, 3.7] log units. The OPC and TIP3P-FB containing combinations had a higher RMSE range of 3.2 [2.0, 4.5] to 4.0 [2.7, 5.3] log units.

Physical reference calculations also included wet and dry conditions for the octanol phase using the GAFF and SMIRNOFF force field with TIP3P water. The wet octanol phase was composed of 27% water and dry octanol was modeled as pure octanol (0% water content). For reference calculations with the TIP3P water model the GAFF, and SMIRNOFF force fields using wet or dry octanol phases resulted in statistically indistinguishable performance. With GAFF, the dry octanol (*REF07*) RMSE was 2.4 [1.0, 3.7]. The wet octanol (*REF02*) RMSE was 2.3 [1.1, 3.5]. With SMIRNOFF, the dry octanol *REF08* RMSE was 2.4 [1.0, 3.7], with a wet octanol (*REF05*) RMSE of 2.3 [1.2, 3.5] (Table S4 and S5).

While water model and force field may have significantly impacted differences in performance across methods in some cases in this challenge, we have very few cases – aside from these reference calculations – where submitted protocols differed *only* by force field or water model, making it difficult to know the origin of performance differences for certain.

#### 4.2.2 Different simulation protocols lead to different results between “equivalent” methods that use the same force field and water model

Several participants submitted predictions from physical methods which are equivalent to those used in our reference calculations and use the same force field and water model, which in principle ought to give identical results given adequate simulation time. There were three submissions which used the GAFF force field, TIP3P water model, and wet octanol phase: *6nmtt* (MD-AMBER-wetoct), *v2q0t* (InterX_GAFF_ WET_OCTANOL), and *REF02* (YANK-GAFF-TIP3P-dry-oct). As can be seen in Figure 3A, *v2q0t* (InterX_GAFF_ WET_OCTANOL) showed the best accuracy with an RMSE of 1.31 [0.94, 1.65]. *6nmtt* (MD-AMBER-wetoct) and *REF02* (YANK-GAFF-TIP3P-dry-oct) had higher RMSE values of 1.87 [1.33, 2.45] and 2.29 [1.07, 3.53], respectively. Two methods that used GAFF force field, TIP3P water model and wet octanol phase are *sqosi* (MD-AMBER-dryoct) and *REF07* (YANK-GAFF-TIP3P-dry-oct). These two also have an RMSE difference of 0.7 log *P* units. Although, in terms of overall accuracy there are differences, Figure 8 shows that in terms of individual predictions, submissions using the same force field and water model largely agree for most compounds.

Some discrepancies are observed for molecules SM13 and SM07, but are largest for SM08. For SM13 and SM07, method *v2q0t* (InterX_GAFF_ WET_OCTANOL) performs over 1 log *P* unit better than *6nmtt* (MD-AMBER-wetoct). The rest of the predictions for these two methods differ by no more than about 1 log *P* unit, with the majority of the molecules differing by about 0.5 log *P* units or less from each other. Comparing *6nmtt* (MD-AMBER-wetoct) vs *REF02* (YANK-GAFF-TIP3P-wet-oct) (Figure 8A), there is a substantial difference in the predicted values for molecules SM08 (4.6 log unit difference), SM13 (1.4 log unit difference), and SM07 (1.2 log unit difference). Method *v2q0t* (InterX_GAFF_ WET_OCTANOL) and *6nmtt* (MD-AMBER-wetoct) perform about 5 log *P* units better than *REF02* (YANK-GAFF-TIP3P-wet-oct) for molecule SM08. Besides SM08, predictions from *v2q0t* (InterX_GAFF_ WET_OCTANOL) and *REF02* (YANK-GAFF-TIP3P-wet-oct) differ by 0.5 log *P* units or less from each other. In dry octanol, *REF07* (YANK-GAFF-TIP3P-dry-oct) performs about 4 log *P* units worse than *sqosi* (MD-AMBER-dryoct) for SM08 (Figure 8B).

Submissions *6nmtt* (MD-AMBER-wetoct), *sqosi* (MD-AMBER-dryoct) and *v2q0t* (InterX_GAFF_WET_OCTANOL) used GAFF version 1.4 and the reference calculations used version 1.81, though GAFF differences are not expected to play a significant role here (i.e. only the valence parameters differ).

#### 4.2.3 Selected small molecule state differences may have caused divergence between otherwise equivalent methods

In several of these approaches, users selected their own starting conformation, protonation state and tautomer, rather than those provided in the SAMPL6 challenge, so the differences here could possibly be attributed to differences in tautomer or resonance structures. Submissions *6nmtt* (MD-AMBER-wetoct) and *sqosi* (MD-AMBER-dryoct) used different tautomers for SM08 and different resonance structures for SM11 and SM14 (microstates SM08_micro010, SM11_micro005, SM14_micro001 from the previous SAMPL6 p*K*_a_ Challenge). We will discuss possible differences due to tautomer choice below in Section 4.2.6. The majority of the calculated log *P* values in *6nmtt* (MD-AMBER-wetoct), *sqosi* (MD-AMBER-dryoct), *v2q0t* (InterX_GAFF_WET_OCTANOL), *REF02* (YANK-GAFF-TIP3P-wet-oct), and *REF07* (YANK-GAFF-TIP3P-dry-oct) show the molecules having a greater preference for octanol over water than the experimental measurements (Figure 8A, B). Methods *6nmtt* (MD-AMBER-wetoct) and *REF02* (YANK-GAFF-TIP3P-wet-oct) overestimate log *P* more than *v2q0t* (InterX_GAFF_WET_OCTANOL) (Figure 8A). Method *REF07* (YANK-GAFF-TIP3P-dry-oct) overestimates log *P* slightly more than *sqosi* (MD-AMBER-dryoct) (Figure 8B).

Three equivalent wet octanol methods and 2 equivalent dry octanol methods gave dissimilar results, and specific molecules were identified that show the major differences in predicted values (Figure 8C-F). GAFF and the TIP3P water model were used in all of these cases, but different simulation setups and codes were used, as well as different equilibration protocols and production methods. Submissions *6nmtt* (MD-AMBER-wetoct) and *sqosi* (MD-AMBER-dryoct), which come from the same group, used 10 ps NPT, 15 ns additional equilibration with MD, and Thermodynamic integration for production in their setup. Submission *v2q0t* (InterX_GAFF_WET_OCTANOL) used 200 ns of molecular dynamics to pre-equilibrate octanol systems, 10 ns of Temperature replica exchange in equilibration, and Isothermal-isobaric ensemble based molecular dynamics simulations in production. The reference calculations (*REF02* and *REF07*) were equilibrated for about 500 ns and used Hamiltonian replica exchange in production. Reference calculations performed with the IFE protocol and MD-AMBER-dryoct (*sqosi*) method used shorter equilibration times than the DFE protocol (*REF07*).

#### 4.2.4 DFE and IFE protocols led to indistinguishable performance, except for SM08 and SM02

The direct transfer free energy (DFE) protocol was used for the physical reference calculations (*REF01-REF08*). Because the DFE protocol implemented in YANK [61] (which was also used in our reference calculations (*REF01-REF08*)) was relatively untested (see Section 12.1.1 for more details), we wanted to ensure it had not dramatically affected performance, so we compared it to the indirect solvation-based transfer free energy protocol (IFE) [103] protocol. The DFE protocol directly computed the transfer free energy between solvents without any gas phase calculation, whereas the IFE protocol (used in the blind submissions and some additional reference calculations labeled IFE) computed gas-to-solution solvation free energies in water and octanol separately and then subtracted to obtain the transfer free energy. The IFE protocol calculates the transfer free energy as the difference between the solvation free energy of the solute going from the gas to the octanol phase, and the hydration free energy going from the gas to the water phase. These protocols ought to yield equivalent results in the limit of sufficient sampling, but may have different convergence behavior.

Figure 9A shows calculations from our two different reference protocols using the DFE and IFE methods. We find that the two protocols yield similar results, with the exception of two molecules. Molecule SM08 is not substantially overestimated using the IFE protocol, where it is with the DFE protocol, and SM02 is largely overestimated by IFE, but not DFE (Figure 9A). The DFE (*REF07*)and IFE protocol both tend to overestimate the molecules’ preference for octanol over water than in experiment, with the DFE protocol overestimating it slightly more. Figure 9D shows comparison of predicted log *P* values of the same tautomers by the DFE (*REF07*) and IFE protocols. The DFE and IFE protocols are almost within statistical error of one another, with the largest discrepancies coming from SM02 and SM08. The DFE and IFE protocols are in better agreement for some tautomers of SM08 more than others. They agree better on the predicted values for SM08_micro008 and SM08_micro010 than for SM08_micro011.

In the SAMPL6 blind submissions, there was a third putatively equivalent method to our reference predictions with the DFE protocol (*REF07*) and IFE protocol: *sqosi* (MD-AMBER-dryoct). It is identical in chosen force field, water model, and composition of octanol phase, however, different tautomers and resonance states for some molecules were used. All three predictions used free energy calculations with GAFF, TIP3P water, and a dry octanol phase. Additionally, *sqosi* (MD-AMBER-dryoct) also used the more traditional indirect solvation free energy protocol. We chose to investigate the differences in these equivalent approaches approaches by comparing predictions using matching tautomers and resonance structures (Figure 9). Figure 9B shows comparison of these three methods using predictions made with DFE and IFE protocols using identical tautomer and resonance input states as *sqosi* (MD-AMBER-dryoct): SM08_micro010, SM11_micro005, and SM14_micro001 (structures can be found in Figure 11. Except SM02, there is general agreement between these predictions. Figure 9C, other than the SM08_micro010 tautomer, predictions of DFE (*REF07*) and *sqosi* (MD-AMBER-dryoct) largely agree. Figure 9E highlights SM02 and SM08_micro010 predictions as the major differences between our predictions with IFE protocol and *sqosi* (MD-AMBER-dryoct).

Only results from the DFE protocol were assigned submission numbers (of the form *REF##*) and presented in the overall method analysis in Section 4.1. More details of the solvation and transfer free energy protocol can be found in section 12.1.

#### 4.2.5 SM08 and SM13 were the most challenging for physical reference calculations

For the physical reference calculations category, some of the challenge molecules were harder to predict than others (Figure 10). Overall, the chemical diversity in the SAMPL6 Challenge dataset was limited. This set has 6 molecules with 4-amino quinazoline groups and 2 molecules with a benzimidazole group. The experimental values have a narrow dynamic range from 1.95 to 4.09 and the number of heavy atoms ranges from 16 to 22 (with the average being 19), and the number of rotatable bonds ranges from 1 to 4 (with most having 3 rotatable bonds). SM13 had the highest number of rotatable bonds and number of heavy atoms. This molecule was overestimated in the reference calculations. As noted earlier, molecule SM08, a carboxylic acid, was predicted poorly across all reference calculations. The origin of problems with molecule SM08 are discussed below in Section 4.2.6.

SM08 is a carboxylic acid and can potentially form an internal hydrogen bond. This molecule was greatly overestimated in the physical reference calculations. When this one molecule in the set is omitted from the analysis, log *P* prediction accuracy improves. For example, the average RMSE and R^2^ values across all of the physical reference calculations when the carboxylic acid is included are 2.9 (2.3–4.0 RMSE range) and 0.2 (0.1–0.2 R^2^ range), respectively. Excluding this molecule gives an average RMSE of 2.1 (1.3–3,3 RMSE range) and R^2^ of 0.57 (0.3–0.7 R^2^ range), which is still considerably worse than best-performing methods.

#### 4.2.6 Choice of tautomer, resonance state, and assignment of partial charges impact log *P* predictions appreciably

Some physical submissions selected alternate tautomers or resonance structures for some compounds. Figure 11 shows three tautomers of SM08, and two alternative resonance structures of SM11 and SM14, all of which were considered by some participants. The leftmost structure of the alternate structure group of each molecule depicts the structure provided to participants.

Because some participants chose alternate structures, we explored how much variation in the selected input structures impacted the results. Particularly, for molecules SM14, SM11 and SM08, both AMBER MD protocols (submissions *sqosi* (MD-AMBER-dryoct) and *6nmtt* (MD-AMBER-wetoct)) used the SM14_micro001 microstate for SM14, SM11_micro005 for SM11 and SM08_micro010 for SM08, rather than the input structures provided as SMILES in the SAMPL6 log *P* Challenge instructions (See Figure 11 for depictions). The reference calculations and the submission from InterX (*v2q0t* (InterX_GAFF_WET_OCTANOL)) used the exact input structures provided as input SMILES for the challenge. Below, we refer to these several submissions as the MD-AMBER and InterX submissions.

To assess whether the choice of tautomer or resonance structure was important, we performed direct transfer free energy (DFE) and indirect solvation-based transfer free energy (IFE) [103] calculations for these alternate structures (please refer to section 4.2.4 for an explanation of the DFE and IFE methods). We —and the other participants utilizing these MM-based methods— assumed that the tautomers and resonance structures are fixed on transfer between phases and we did not do any assessments of how such populations might shift between octanol and water. Table 5 and Figure 9B compare log *P* calculations starting from the same input structures across the three methods: *sqosi* (MD-AMBER-dryoct), *REF07* (YANK-GAFF-TIP3P-dry-oct) which used the DFE protocol, and an additional set of calculations with the IFE protocol (using YANK, the GAFF force field, and TIP3P water just like *REF07*). The DFE protocol prediction set presented in Figure 9A is the same as *REF07* (YANK-GAFF-TIP3P-dry-oct), but includes extra tautomers for SM08, and extra resonance structures for SM11 and SM14.

From our comparison of our reference calculations and those with the InterX and MD-AMBER, we find that the choice of input tautomer has a significant effect on log *P* predictions. Particularly, within the traditional IFE method, our results indicate up to 2.7 log units variation between log *P* values for different tautomers of SM08 (between SM08_micro011, SM08_micro08 and SM08_micro10) (Table 5). Our exploration of these issues was prompted by the fact that the MD-AMBER protocols had utilized different tautomers than those initially employed in our physical reference calculations.

We also find that the choice of resonance structure affects calculated values, though less strongly so than the choice of tautomer. Within the IFE method we find 1.3 log units of variation between log *P* values calculated with different resonance structures of SM11 (SM11 and SM11_micro005) and 0.6 log units of variation between resonance structures of SM14 (SM14 and SM14_micro001) (Table 5).

We also find that the partial charge assignment procedure can also dramatically impact log *P* values for carboxylic acids (Table S7). Particularly, our calculations with the DFE and IFE protocols employed different partial charge assignment procedures as an unintentional feature of the protocol difference, as we detail below, and this impacted calculated log *P* values by up to 6.7 log units for SM08 (specifically SM08_micro011, the carboxylic acid in the set) compared to experiment. Particularly, the DFE protocol utilized antechamber for assigning AM1-BCC charges, whereas the IFE protocol used OpenEye’s quacpac. Antechamber utilizes the provided conformer (in this case, the *anti* conformation) for each molecule, whereas quacpac’s procedure computes charges for carboxylic acids in the *syn* conformation because this has been viewed as the relevant conformation, and because of concerns that the *anti* conformation might result in unusually large and inappropriate charges. Thus, because of this difference, the DFE and IFE protocols used dramatically different partial charges for these molecules (Table S7). Our results for SM08_micro011 (likely the dominant state) indicate that indeed, the conformer used for charging plays a major role in assigned charges and the resulting log *P* values (Table S7, Figure S2). We find our DFE protocol, which used the *anti* conformation for charging, overestimates the log *P* by about 6.7 log units, whereas the IFE protocol which used the *syn* conformation only overestimates it by about 0.3 log units. With the IFE method, we calculated a log *P* of 2.8 ± 0.2 for SM08, whereas with DFE method we obtained a value of 9.8 ± 0.1 (Table 5).

### 4.3 Lessons learned from empirical reference calculations

Empirical methods are fast and can be applied to large virtual libraries (100 000 cmps/min/CPU). This is in contrast to physical methods, which are often far more computationally demanding. Most of the empirical methods are among the top performers, with the exception of a few approaches that use descriptors and/or pre-factors that do not yield accurate log *P* predictions. Most empirical methods obtain RMSE and MAE values below 1 log *P* unit. The best empirical method achieved RMSE and MAE below 0.5 (*gmoq5*, Global XGBoost-Based QSPR LogP Predictor). In all these cases, using a relatively large training set (>1000-10000 compounds) seems to be key.

The exact choice of method or descriptors seems to be less critical. Predictions based on atom or group contributions perform as well as those using either a small set of EHT-derived descriptors or a large set of diverse descriptors, sometimes additionally including fingerprint descriptors. A possible explanation could be that log *P* is, to first order, primarily an additive property so that empirical methods can do well since a wealth of octanol-water data is available for training. This is also reflected in the success of the simple methods summing up atom contributions. This approach may become problematic, however, when a functional group is present that was underrepresented or missing in the training set. In such cases, higher are expected.

As is true for the physical methods, empirical methods depend on the tautomeric state of the compound. Here we have observed that clogP is particularly sensitive. clogP shifts of more than one log unit upon change of the tautomer are not uncommon. h_logP is much less sensitive to tautomers with shifts usually below 0.5 log *P* units. This is also true for molecule SM08, for which different tautomeric forms are possible (as seen in Figure 11). For the pyridone form of SM08 (SM08_micro011), clogP predicts a log *P* of 2.17, whereas the hydroxy-pyridine form (SM08_micro010) yields a log *P* of 3.63. For h_logP, the respective values are 3.09 and 3.06.

Despite the small training sets of the MOE models, good prediction for kinase inhibitor fragments and the extra compounds was achieved. This is possibly because the training set for this model was biased towards drug-like compounds, with substantial similarity to the SAMPL6 Challenge set.

Other studies have found that some empirical methods tend to overestimate log *P* when molecular weight increases [104, 105]. In this challenge, this was less of an issue as molecular size remained relatively constant.

According to in-house experience at Boehringer-Ingelheim, different experimental log *P* measurement methods produce values that are correlated with one another with an R^2^ value of around 0.7 (T. Fox, P. Sieger, unpublished results), indicating that experimental methods themselves can disagree with one another significantly. This is especially true when it comes to more approximate methods of estimating log *P* experimentally, such as HPLC-based methods [42, 106]. A dataset composed of 400 compounds from Boehringer-Ingelheim measured both with GLpKa and HPLC assays covering a range from 0-7 log *P* units had R^2^ of 0.56, though in some cases these methods may have higher correlations with potentiometric approaches [107]. Thus, if an empirical model is trained on log *P* data from one particular method, testing it on data collected via another method may not yield performance as high as expected.

Here, all of the analyzed empirical reference methods achieved absolute error <2.0, and often <1.5 calculated for each molecule in the SAMPL6 log *P* Challenge set. This is a sign of more consistent accuracy of the predictions across different molecules compared to physical methods. However, it is difficult to draw general conclusions given the small size of the data set, and many hypotheses being based on only one example.

### 4.4 Performance of reference methods on additional molecules

To broaden the analysis with a larger set with more chemical diversity and larger dynamic range of log *P* values, an extra set of 27 compounds were included in the analysis of reference calculations (Figure 12). These compounds had literature experimental log *P* values collected using the same method as the SAMPL6 dataset. This set is composed of substituted phenols, substituted quinolines, barbiturate derivatives and other pharmaceutically relevant compounds [108].

**Figure 12.**
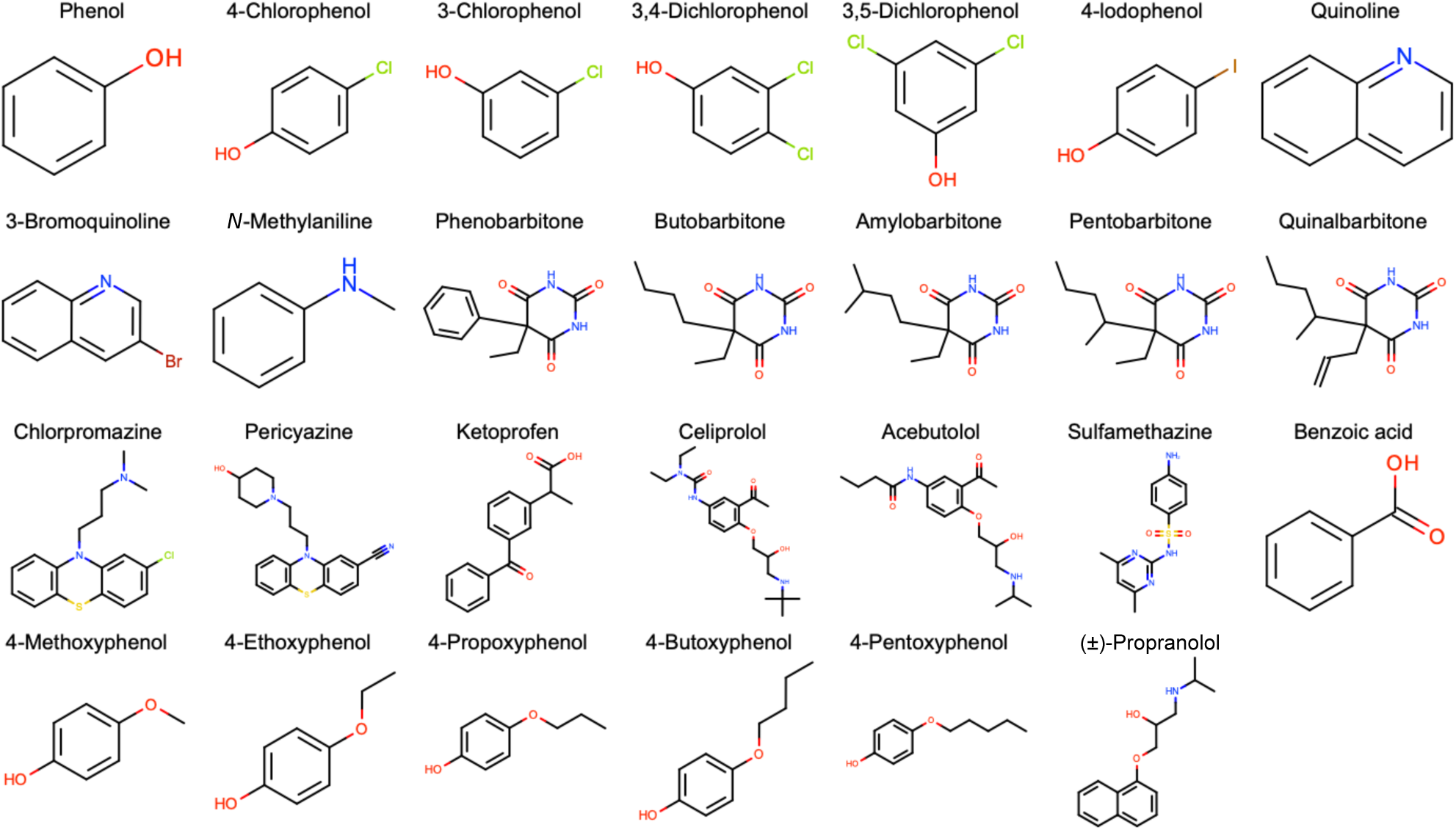
Structures of the 27 additional molecules that were included in follow-up assessment of the reference methods. These molecules were not included in the statistics overview.

This set of molecules is larger and more diverse than the SAMPL6 challenge set, spanning a range of 4.5 log units compared to the challenge set which had a range of 2.1 log units. For this set, the number of rotatable bonds ranges from 0 to 12, with an average of 3 per compound. The number of heavy atoms ranges from 7 to 27, and the average per compound is 14. Most of the worst-performing compounds for the physical reference calculations had a higher number of heavy atoms – celiprolol (27), acebutolol (24) and pericyazine (26). Celiprolol and acebutolol both have the highest number of rotatable bonds in the set, 12 and 11 respectively. Chlorpromazine, pericyazine, and sulfamethazine all contain sulfur. Sulfur can in some cases pose particular challenges for force fields, especially hypervalent sulfur [109], which may account for the poor performance of pericyazine, chlorpromazine, and sulfamethazine. Pericyazine, one of the worst performing compounds, is also the only molecule in the set that has a nitrile.

In the physical reference calculations, the mean absolute errors are below 1 log *P* unit for dry octanol conditions and below 1 log *P* units for wet octanol conditions (Table 6). The calculated log *P* values had an average RMSE of 1.4 (RMSE range of 1.3 [0.9, 1.6] to 1.5 [1.0, 2.0]), and an average R^2^ of 0.5 (with a correlation range of 0.5 [0.1, 0.8] to 0.6 [0.3, 0.8]). Physical methods are on par with empirical ones for the smaller, less flexible compounds, but in general are worse, especially for compounds with long flexible hydrophilic tails. The exception is chlorpromazine, but the smaller error seen in this molecule might be due to an error compensation caused by the presence of the sulfur atom since force fields have challenges with sulfur-containing compounds. [109].

**Table 6.** Statistics of the physical and empirical reference method predictions on the extra test of molecules. Methods were ranked according to increasing RMSE in this table. Performance statistics of MAE, R^2^, and Kendall’s Tau are also provided. Mean and 95% confidence intervals of all statistics are presented.

| ID | Method Name | Category | Type | RMSE | MAE | $R^2$ | Kendall's Tau ($\tau$) |
| --- | --- | --- | --- | --- | --- | --- | --- |
| EXT09 | clogP (Biobyte) | Empirical | Reference | 0.23 [0.16, 0.29] | 0.17 [0.12, 0.23] | 0.94 [0.86, 0.98] | 0.85 [0.75, 0.93] |
| EXT12 | MoKa_logP | Empirical | Reference | 0.28 [0.20, 0.35] | 0.22 [0.16, 0.28] | 0.91 [0.82, 0.97] | 0.83 [0.73, 0.91] |
| EXT11 | logP(o/w) (MOE) | Empirical | Reference | 0.32 [0.22, 0.41] | 0.24 [0.17, 0.33] | 0.90 [0.81, 0.95] | 0.80 [0.66, 0.91] |
| EXT10 | h_logP (MOE) | Empirical | Reference | 0.43 [0.34, 0.51] | 0.35 [0.26, 0.45] | 0.83 [0.62, 0.93] | 0.74 [0.55, 0.90] |
| EXT13 | SlogP (MOE) | Empirical | Reference | 0.59 [0.48, 0.69] | 0.49 [0.36, 0.61] | 0.71 [0.41, 0.87] | 0.55 [0.33, 0.73] |
| EXT08 | YANK-SMIRNOFF-TIP3P-dry-oct | Physical (MM) | Reference | 1.26 [0.88, 1.60] | 0.97 [0.69, 1.28] | 0.56 [0.16, 0.83] | 0.50 [0.23, 0.73] |
| EXT07 | YANK-GAFF-TIP3P-dry-oct | Physical (MM) | Reference | 1.27 [0.69, 1.74] | 0.88 [0.57, 1.26] | 0.55 [0.19, 0.88] | 0.60 [0.34, 0.81] |
| EXT02 | YANK-GAFF-TIP3P-wet-oct | Physical (MM) | Reference | 1.38 [0.94, 1.78] | 1.03 [0.70, 1.40] | 0.58 [0.26, 0.83] | 0.58 [0.35, 0.78] |
| EXT05 | YANK-SMIRNOFF-TIP3P-wet-oct | Physical (MM) | Reference | 1.50 [0.96, 1.98] | 1.11 [0.75, 1.52] | 0.50 [0.13, 0.81] | 0.54 [0.29, 0.75] |

Empirical methods are more stable in the sense that there are no gross outliers found in the extended set. For the empirical reference calculations, the absolute errors for the 27 extra compounds are all below 1 log *P* unit. For clogP, most compounds have errors below 0.4 log *P* unit, with only (±)-propanolol a bit higher. Compound 3,5-dichlorophenol and 3,4-dichlorophenol consistently had a slightly higher error; there is no obvious correlation between method performance and size or complexity of the compounds. Figure 13A shows that 3,5-dichlorophenol, 3,4-dichlorophenol, and (±)-propranolol were the most challenging compounds for empirical reference methods. The MAE calculated for these three molecules as the average of five methods (*EXT09, EXT10, EXT11, EXT12, EXT13*) was higher than 0.5 log *P* units (Table 6). RMSE overall compounds between 0.23 for clogP and 0.59 for MOE_SlogP, significantly below the best physical model. This is mirrored in the Kendall tau values, where the best empirical method (clogP) achieves 0.85, whereas the best physical methods are comparable to the worst empirical method with a value of 0.55.

**Figure 13.**
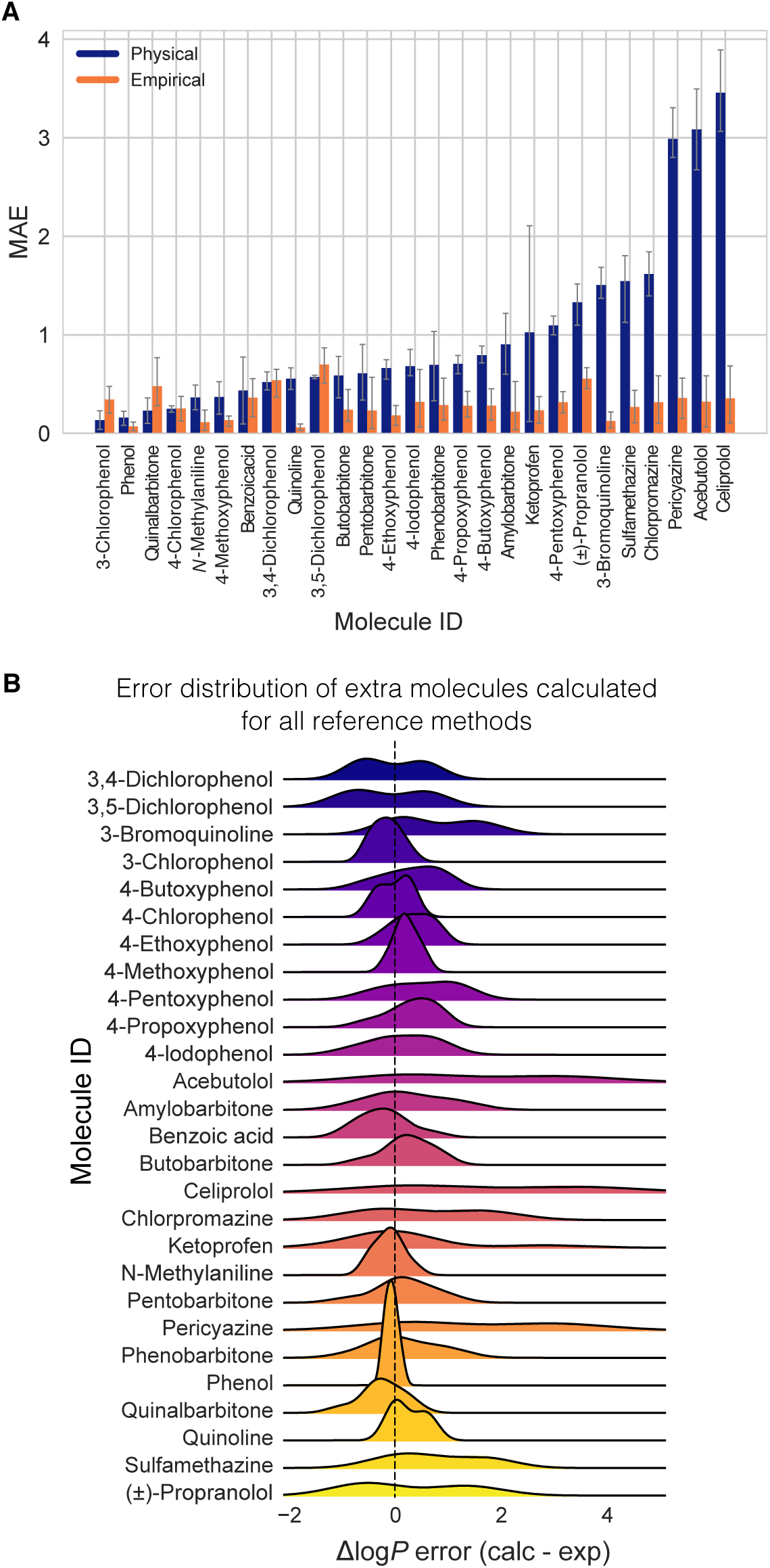
Distribution of reference method calculation errors by molecule on our extra set shows that a few of the molecules were more challenging than others. (A) MAE of each of the extra molecules broken out by physical and empirical reference method category. Majority of molecules have mean absolute errors below 1 log *P* unit for physical reference calculations. All of the mean absolute errors are well below 1 log *P* unit for empirical reference calculations. (B) Error distribution for each molecule calculated for the reference methods. A couple molecules have a significant tail showing probability of overestimated log *P* predictions.

When prediction performance of empirical prediction methods for this dataset and for the SAMPL6 Challenge set are compared, we observe better prediction accuracy for this set, with an RMSE range of 0.2 [0.2, 0.3] to 0.6 [0.5,0.7] for the extra molecules and 0.5 [0.2, 0.8] to 0.8 [0.5, 1.1] for the challenge molecules. The average R^2^ was 0.6 (with a correlation range of 0.38 [0.01, 0.82] to 0.7 [0.3, 0.9]). This may be due to SAMPL6 compounds being more challenging, or it may be that these extra molecules appear in the training sets used in developing empirical methods.

### 4.5 Take-away lessons from the SAMPL6 Challenge

Empirical and QM-based prediction methods represented in SAMPL6 Challenge in general performed better than MM-based methods. Ten empirical and QM-based methods achieved an RMSE < 0.5 log *P* units. The lowest RMSE observed for MM-based physical methods was 0.74 and the average RMSE of the better half of MM-based methods was 1.44 log *P* units. However, the RMSE of the best two MM-based methods was similar to the null model, which simply guessed that all compounds had a constant, typical log *P* value.

For MM approaches, prediction accuracy varied based on methodological choices such as simulation method, equilibration protocol, free energy estimation method, force field, and water model. Only a small number of MM-based physical models achieved an accuracy similar to the null model, which had an RMSE of 0.8. Some MM methods outperformed the null model, but such performance was variable across approaches and not clearly linked to a single choice of force field, water model, etc. Polarizable force fields also did not provide an advantage for log *P* predictions, possibly due to solute simplicity and the absence of formal charge, or because other sources of error dominated.

Analysis of the MM-based reference calculations highlighted equilibration and charging protocols, sampling challenges, identification of the dominant neutral tautomer, and selection of input resonance states as confounding factors. Comparison of equivalent calculations from independent participants (identical methods such as free energy calculations using GAFF and TIP3P water with different setups and code) showed significant systematic deviations between predicted values for some compounds. The comparison of identical methods also showed that the tautomer and resonance state choice for some molecules resulted in discrepancies in calculated log *P* values. In one case, conformation selected for a carboxylic acid before charging was important. We have also noticed differences in in equilibration protocols, which could be particularly important for the octanol phase, though the present challenge does not conclusively demonstrate that differences in equilibration made a significant difference.

Fast empirical methods showed greater consistency of prediction accuracy across test molecules compared to physical methods. Most of the empirical methods were among the better-performing methods. The size of the training sets seems to be more important for accuracy than the exact methods and descriptors used in each model. Although not observed in the SAMPL6 Challenge set, empirical methods may experience problems if a functional group is underrepresented in training sets. Just like the physical methods, the choice of tautomer makes a difference. For example, shifts greater than 1 log unit in the calculated log *P* of different tautomers are common.

Performance in the SAMPL6 log *P* challenge was generally better than in the SAMPL5 log *D* Challenge. The change of partition solvent from cyclohexane to octanol, absence of protonation state effects, and smaller chemical diversity represented in the challenge are likely reasons. In the SAMPL5 log *D* Challenge, only five submissions had an RMSE below 2.5 log units, while here, 10 methods achieved an RMSE ≤ 0.5 log *P* units and many of the submissions had an RMSE ≤ 1.0 log *P* units. The design of the SAMPL6 log *P* Challenge removed some of the factors confounding accuracy in the earlier log *D* challenge, namely p*K*_a_ prediction and cyclohexane (a challenging solvent for empirical methods).

Compared to expected accuracy for partition coefficients based on SAMPL4 Challenge performance, many QM-based methods were better while only a small number of MM-based methods achieved slightly better results. In SAMPL4, the top-performing hydration free energy predictions had an error of about 1.5 kcal/mol, which would yield an expected error here (assuming independent errors/no error cancellation) of about 1.54 log units [2], if log *P* values were estimated from a difference in solvation free energies. Many physical methods achieved roughly this accuracy or slightly better.

Partition coefficient predictions can also serve, for physical calculations, as a model system that reflects how well solvation effects can be captured by the same techniques developed for protein-ligand binding predictions – where solvation also plays a role in calculations. Relative binding free energy calculations tend to achieve errors, in the best-case scenario, in the 1–2 kcal/mol range [110], or about 1.03–2.06 log units if similar accuracy were achieved here for solvation in each phase (with independent errors). Many methods did better than 2 log *P* units of error in this challenge, which is in agreement with the expectation that partition coefficients present an easier model system compared to protein-ligand binding affinities.

Performance of empirical methods far surpassed these thresholds taking advantage of the available octanol-water experimental data, however, these empirical techniques are specifically oriented towards predicting partitioning and cannot be applied to the binding problem.

### 4.6 Suggestions for the design of future challenges

In the SAMPL6 Challenge, the log *P* focus proved helpful to allow a focus on modeling of solvation effects without the complexities of modeling different protonation states present in a log *D* challenge. Challenges which focus on specific aspects of modeling help isolate methodological problems, making challenges like log *P* and log *D* modeling particularly helpful. We believe the largest benefits to the field will be achieved from iterated challenges, as seen from the progress achieved in predicting hydration free energies over multiple SAMPL challenges [60].

As MM-based physical methods struggled with octanol-water log *P* predictions in SAMPL6, we recommend additional SAMPL iterations focused on log *P* with larger datasets and more chemical diversity to facilitate progress. The conclusions of SAMPL6 p*K*_a_ and log *P* Challenges indicate that, if this had been posed as a log *D* challenge rather than a log *P* challenge, larger p*K*_a_ prediction errors would have masked underlying issues in predicting equilibrium partitioning of neutral solutes between solvent phases. The fact that performance for physical methods was still relatively poor illustrates the potential benefit of future log *P* challenges.

For near-term challenges, we would like to keep the level of difficulty reasonable by keeping the focus on smaller and fragment-like compounds and limiting the number of non-terminal rotatable bonds (maximum of 6) similar to SAMPL5. The SAMPL5 Challenge suggested that molecules with many rotatable bonds still pose challenges for contemporary methods, suggesting this is a criterion for difficulty. However, in later challenges we hope to gradually increase the difficulty of the compounds considered to provide a more diverse set that includes more difficult compounds including varying numbers of rotatable bonds.

Ideally, a more diverse combination of functional groups in the compounds should be included in future sets, with improved chemical diversity posing more challenges and also helping provide additional lessons learned. For example, a dataset could include matched molecular pairs which differ by only a single functional group, helping to isolate which functional groups pose particular challenges. Current MM-based methods are known to often have difficulty modeling sulfonyl and sulfonamide groups, but a challenge utilizing matched molecular pairs could reveal other such challenging functional groups. In addition, expanding partition coefficient challenges with a diverse set of solvent phases would be beneficial for improving solute partitioning models.

The statistical power of the SAMPL6 log *P* Challenge for comparative method evaluation was limited due to the narrow experimental data set with only 2 log *P* units of dynamic range and 11 data points, both of which were driven by limitations of the experimental methodology chosen for this challenge [9]. Future log *P* challenges would benefit from larger blind datasets with a broader dynamic range. We recommend at least a log *P* range of 1–5. The potentiometric log *P* measurement method used for the collection for SAMPL6 data was rather low throughput, requiring method optimization for each molecule. High-throughput log *D* measurement methods performed at pHs that would ensure neutral states of the analytes may provide a way to collect larger datasets of log *P* measurements. However, this approach poses some challenges. First, it is necessary to measure p*K*_a_ values of the molecules first. Second, partitioning measurements need to be done at a pH that guarantees that the compound has neutral charge, in which case solubility will be lower than if it is charged and may become a limitation for the experiment.

SAMPL6 log *P* Challenge molecules were not expected to have multiple tautomers affecting log *P* predictions (based on QM predictions). The choice of the challenge set also ensured participants did not have to calculate contributions of multiple relevant tautomerization states or shifts in tautomerization states during transfer between phases. However, participants still had to select a major tautomer for each compound. To evaluate the tautomer predictions in the future, experimental measurement of tautomer populations in each solvent phase would provide valuable information. However, such experimental measurements are difficult and low throughput. If measuring tautomers is not a possibility, the best approach may be to exclude compounds that present potential tautomerization issues from the challenge, unless the challenge focus is specifically on tautomer prediction.

Overall, for future solute partitioning challenges, we would like to focus on fragment-like compounds, matched molecular pairs, larger dynamic range, larger set size, and functional group diversity.

## 5 Conclusion

Several previous SAMPL challenges focused on modeling solvation to help address this key accuracy-limiting component of protein-ligand modeling. Thus, the SAMPL0 through SAMPL4 challenges included hydration free energy prediction as a component, followed by cyclohexane-water distribution coefficient in SAMPL5.

Here, a community-wide blind partition coefficient prediction challenge was fielded for the first time, and participants were asked to predict octanol-water partition coefficients for small molecules resembling fragments of kinase inhibitors. As predicting log *D* in the previous challenge was quite challenging due to issues with p*K*_a_ prediction, the present challenge focused on log *P*, avoiding these challenges and placing it at roughly the right level of complexity for evaluating contemporary methods and issues they face regarding the modeling of small molecule solvation in different liquid phases. The set of molecules selected for the challenge were small and relatively rigid fragment-like compounds without tautomerization issues which further reduces the difficulty of the prospective prediction challenge.

Participation in the challenge was much higher than in SAMPL5, and included submissions from many diverse methods. A total of 27 research groups participated, submitting 91 blind submissions in total. The best prospective prediction performance observed in SAMPL6 log *P* Challenge came from QM-based physical modeling methods and empirical knowledge-based methods, with 10 methods achieving an RMSE below 0.5 log *P* units. On the other hand, only a small number of MM-based physical models achieved an accuracy similar to the null model (which predicted a constant, typical log *P* value), which had an RMSE of 0.8. Empirical predictions showed performance which was less dependent on the compound/dataset than physical methods in this study. For empirical methods, the size and chemical diversity of the training set employed in developing the method seems to be more important than the exact methods and descriptors employed. We expected many of the empirical methods to be the top performers, given the wealth of octanol-water log *P* training data available, and this expectation was borne out.

Better prediction performance was seen for octanol-water log *P* challenge than the SAMPL5 cyclohexane-water log *D* challenge. In addition to absence of p*K* _a_ prediction problem for the partition system, the molecules in the SAMPL6 log *P* Challenge were considerably less diverse than in the SAMPL5 log *D* Challenge, which may have also affected relative performance in the two challenges. Physical methods fared slightly better in this challenge than previous cyclohexane-water log *D* challenge, likely because of the elimination of the need to consider protonation state effects. However, MM-based physical methods with similar approaches did not necessarily agree on predicted values, with occasionally large discrepancies resulting from apparently relatively modest variations in protocol.

All information regarding the challenge structure, experimental data, blind prediction submission sets, and evaluation of methods is available in the SAMPL6 GitHub Repository to allow follow up analysis and additional method testing.

Overall, high participation and clear lessons learned pave the way forward for improving solute partitioning and biomolecular binding models for structure-based drug design.

## Supporting information

SAMPL6-supplementary-documents.tar.gz

## 0.2 Abbreviations

SAMPL: Statistical Assessment of the Modeling of Proteins and Ligands
log *P*: log_10_ of the organic solvent-water partition coefficient (*K*_*ow*_) of neutral species
log *D*: log_10_ of organic solvent-water distribution coefficient (*D*_*ow*_)
p*K*_a_: −log_10_ of the acid dissociation equilibrium constant
SEM: Standard error of the mean
RMSE: Root mean squared erro
MAE: Mean absolute error
*τ*: Kendall’s rank correlation coefficient (Tau)
R^2^: Coefficient of determination (R-Squared)
QM: Quantum Mechanics
MM: Molecular Mechanics

## 6 Code and Data Availability

All SAMPL6 log *P* Challenge instructions, submissions, experimental data and analysis are available at https://github.com/samplchallenges/SAMPL6/tree/master/physical_properties/logP. An archive copy of SAMPL6 GitHub Repository log *P* Challenge directory is also available in the Supplementary Documents bundle (*SAMPL6-supplementary-documents.tar.gz*). Some useful files from this repository are highlighted below.

- Table of participants and their submission filenames: https://github.com/samplchallenges/SAMPL6/blob/master/physical_properties/logP/predictions/SAMPL6-user-map-logP.csv
- Table of methods including submission IDs, method names, participant assigned method category, and reassigned method categories: https://github.com/samplchallenges/SAMPL6/blob/master/physical_properties/logP/predictions/SAMPL6-logP-method-map.csv
- Submission files of prediction sets: https://github.com/samplchallenges/SAMPL6/tree/master/physical_properties/logP/predictions/submission_files
- Python analysis scripts and outputs: https://github.com/samplchallenges/SAMPL6/blob/master/physical_properties/logP/analysis_with_reassigned_categories/
- Table of performance statistics calculated for all methods: https://github.com/samplchallenges/SAMPL6/blob/master/physical_properties/logP/analysis_with_reassigned_categories/analysis_outputs_withrefs/StatisticsTables/statistics.csv

## 7 Overview of Supplementary Information

### Contents of Supplemantary Information

- **Detailed methods section**:
  1. Physical reference calculations -Direct Transfer Free Energy Approach
  2. Physical reference calculations -Indirect Solvation-Based Transfer Free Energy Approach
  3. Empirical reference calculations
- **Table S1** Method details of log *P* predictions with MM-based physical methods.
- **Table S2** SMILES and InChI identifiers of SAMPL6 log *P* Challenge molecules.
- **Table S3** SMILES and InChI identifiers of extra molecules included in the evaluation of reference methods.
- **Table S4 and Table S5** Evaluation statistics calculated for all methods.
- **Table S6** Comparison of force field parameters of the TIP3P, TIP3P-FB and OPC water models
- **Table S7** Comparison of the charges assigned to the syn and anti conformation of SM08_micro011 in the DFE protocol
- **Figure S1:** Varying the amount of water in the octanol phase has no significant effect on the predicted log *P* in reference calculations.
- **Figure S2** 2D and 3D structures of SM08_micro011 with the carboxylic acid in “anti” and “syn” conformation.
- **Figure S3** For the DFE method, the starting conformation impacts the number of C-O dihedral transitions for SM08_micro011.

### Additional supplementary files

*SAMPL6-supplementary-documents.tar.gz* file includes:

- An archive copy of the log *P* Challenge directory of SAMPL6 GitHub Repository (*SAMPL6-repository-logP-directory.zip*)
- SAMPL6 log *P* Challenge Instructions (*logP_challenge_instructions.md*)
- Table S1 in CSV format (*SI-table-MM-method-details.csv*)
- Table S2 in CSV format (*SAMPL6-logP-chemical-identifiers-table.csv*)
- Table S3 in CSV format (*extra-chemical-identifiers-table.csv*)
- Table S4 and Table S5 in CSV format (*statistics.csv*)
- The free energy and enthalpy values of each phase in triplicate and comparisons of calculated solvation free energies across trials for the physical reference calculations (*analysis-of-physical-reference-calculations.zip*)
- Scripts related to the physical reference calculations (*physical-reference-calculation-scripts.zip*)

## 8 Author Contributions

Conceptualization, MI, JDC, DLM; Methodology, MI, TDB, DM, JDC; Software, MI, TDB, AR; Formal Analysis, MI, TDB; Investigation, MI, TDB, DLM, TF; Resources, JDC, DLM; Data Curation, MI, TDB; Writing-Original Draft, MI, TDB, DLM, TF; Writing - Review and Editing, MI, TDB, DLM, TF, JDC, AZ; Visualization, MI, TDB; Supervision, DLM, JDC; Project Administration, MI; Funding Acquisition, DLM, JDC, MI, TDB.

## 9 Acknowledgments

We would like to thank OpenEye, especially Gaetano Calabró, for help with Orion, and for constructing the Orion workflows partially utilized here. We would like to thank experimental collaborators Timothy Rhodes (ORCID: 0000-0001-7534-9221), Dorothy Levorse, and Brad Sherborne (ORCID: 0000-0002-0037-3427).

MI and JDC acknowledge support from the Sloan Kettering Institute. JDC acknowledges partial support from NIH grant P30 CA008748. MI, TDB, JDC, and DLM gratefully acknowledge support from NIH grant R01GM124270 supporting the SAMPL Blind Challenges. MI acknowledges support from a Doris J. Hutchinson Fellowship during the collection of experimental data. TDB acknowledges support from the ACM SIGHPC/Intel Fellowship. DLM appreciates financial support from the National Institutes of Health (1R01GM108889-01) and the National Science Foundation (CHE 1352608). We acknowledge contributions from Caitlin Bannan who provided feedback on experimental data collection and structure of log *P* challenge from a computational chemist’s perspective. MI and JDC are grateful to OpenEye Scientific for providing a free academic software license for use in this work. TF thanks BioByte, MOE, and Molecular Discovery for allowing us the include log *P* predictions calculated by their software in this work as empirical reference calculations.

## 10 Disclaimers

The content is solely the responsibility of the authors and does not necessarily represent the official views of the National Institutes of Health.

## 12 Supplementary Information

### 12.1 Detailed methods

#### 12.1.1 Physical reference calculations - Direct Transfer Free Energy Approach

log *P* can be estimated directly from the transfer free energy of a solute moving from the organic to the aqueous layer. Specifically, we calculate the transfer free energy from the difference in solvation free energy into octanol and hydration free energy. log *P* is directly proportional to the difference between the solvation free energy for the solute into each solvent

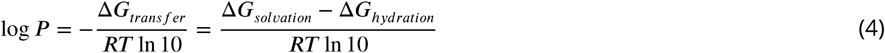

where Δ*G*_*transfer*_ is the transfer free energy, Δ*G*_*soluation*_ is the solvation free energy of the solute going from the gas to the octanol phase, Δ*G*_*hydration*_ is the hydration free energy going from the gas to the water phase, *R* is the gas constant (8.314 J / mol · K) and *T* is the temperature (298.15 K).

The direct transfer free energy (DFE) protocol that was used for the physical reference calculations (*REF01*-REF08, *EXT02, EXT05, EXT07, EXT08*) directly computed the transfer free energy between solvents without any gas phase calculation, whereas the IFE protocol (discussed in Section 12.1.2) computed gas-to-solution phase solvation free energies in water and octanol separately and then subtracted to obtain the transfer free energy.

To explore how solvent mixing would effect predicted values, water was included in the octanol phase for the majority of the reference calculations. A portion of the calculations treated the octanol and water phase as completely immiscible for comparison. The experimental mole fraction of water in octanol was measured as 0.2705 [58]. The solutions and calculations modeled each phase at infinite dilution, with only a single solute molecule in each solvent.

The initial input files were made using the Solvation Toolkit (https://github.com/MobleyLab/SolvationToolkit), which converts SMILES strings to parameterized molecules and builds topology and coordinate files for use in molecular dynamics software packages. Solvation Toolkit is a driver utility that utilizes the OpenEye toolkits (version 2018.10.1) for cheminformatics (specifically file conversion and handling of molecular identities), and OEChem for reading and writing files. AmberTools [111] was used to parameterize systems with the General AMBER Force Field for organic molecules (version 2017.1.81) and water was parameterized with the TIP3P water model, AM1-BCC charges were assigned via Antechamber, Packmol (version 18.169) [112] was used to build boxes, and lastly AMBER topology and coordinate files were made with LEaP.

The SMILES string and the mole fraction of each compound in the system were used as input. The “wet” octanol systems were generated using a mole fraction of 0.7295 for octanol and 0.2705 for water, producing systems with about 200 octanol molecules and 74 water molecules, depending on the solute size. The “dry” octanol systems had no water component and about 211 octanol molecules. All of the water systems had 1497 molecules. The box dimensions were about 40×40 Å in all cases.

The following equilibration stages were carried out using the GAFF forcefield (version 2017.1.81), the TIP3P water model and OpenMM (version 7.3.1) [66, 113], a molecular simulation toolkit.

For minimization, an energy tolerance of 10 kilojoules/mole was used and the systems were minimized until convergence was reached. A Langevin integrator was used with a 0.5 fs timestep. Minimization was followed by 100 ps of NVT using a Langevin integrator and 1.0 fs timestep, 100 ps of NPT using a Langevin integrator and 2.0 fs timestep, and lastly 500 ns of NPT using a Langevin integrator and 2.0 fs timestep.

Three independent equilibrations were run starting from water and octanol phase systems of the initial setup, in order to obtain three different sets of starting coordinates for replicate transfer free energy calculations with YANK (version 0.24.0 [63]). The protocol for creating systems with different force field and/or water model conditions (1) is detailed below.

Following equilibration, the resulting systems were saved to PDBs. For each solvent system, a ParmEd Structure was created using the topology and positions from the equilibrated PDB, with parameters coming from the original GAFF/TIP3P OpenMM System. The ParmEd structure of the system was split into individual components or structures and then used to create newly parameterized OpenMM systems. The water was parameterized with either the TIP3P, TIP3P-FB or OPC water model, and the solute and solvent were parameterized with the SMIRNOFF force field (smirnoff99Frosst version 1.0.7) or remained parameterized with GAFF. In just the OPC case, a dummy atom was added to the water component structure. After parameterization, the OpenMM systems of the solute-octanol and water were converted back to ParmEd structures which maintained their new parameters. The final OpenMM system was created using the particle mesh Ewald (PME) method for periodic boundary conditions, an error tolerance of 1e-4 and a cutoff for nonbonded interactions was set to 11 Å.

The resulting OpenMM Systems were saved as XMLs for use later on. Prior to using YANK, the new systems were briefly equilibrated using the same setup described previously, excluding the 500 ns of NPT. The final equilibrated PDB and system XML files were used as input files for solvation and transfer free energy calculations with YANK [61], a toolkit that uses Hamiltonian replica exchange and can compute solvation free energies. For the YANK simulations, hydrogen mass repartitioning (HMR) was used to allow a 3 fs timestep. HMR works by slowing down the fastest motions in the simulation by reallocating mass from the connected heavy atom to the hydrogens [114]. The temperature was set to 298.15 K (the experimental temperature), the pressure to 1.0 atm, and an anisotropic dispersion cutoff of 12.0 Å was used. There were 5000 iterations total and 335 steps per iteration. The overall length of the YANK simulations were 5 ns for each replica.

In the octanol and water phase the electrostatic interactions of the solute with the solvent were scaled off through a *λ* (lambda) parameter using the following lambda values where *λ* = [1.00, 0.75, 0.50, 0.25, 0.00, 0.00, 0.00, 0.00, 0.00, 0.00, 0.00, 0.00, 0.00, 0.00, 0.00, 0.00, 0.00, 0.00, 0.00, 0.00], and steric interactions were scaled using *λ* = [1.00, 1.00, 1.00, 1.00, 1.00, 0.95, 0.90, 0.80, 0.70, 0.60, 0.50, 0.40, 0.35, 0.30, 0.25, 0.20, 0.15, 0.10, 0.05, 0.00].

The direct transfer free energy was obtained from YANK [61], where the Δ*G*_*transfer*_ was equivalent to Δ*G*_*octanol*_ − Δ*G*_*water*_. This was then converted to log *P* using Equation 4. The uncertainties of the log *P* predictions were calculated as the standard error of the mean (SEM) of three replicate predictions. The SEM was estimated as 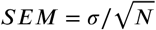 where *σ* is the sample standard deviation and *N* is the size of the sample (in this case the number of replicate predictions made). The model uncertainty was reported as 1.6 log units, based on similar previous work [103].

#### 12.1.2 Physical Reference Calculations - Indirect Solvation-Based Transfer Free Energy Approach

We ran an additional set of reference calculations using a more traditional indirect solvation-based transfer free energy method to see how it would compare to the direct transfer free energy method (described in Section 12.1.1). Specifically, the IFE protocol calculates the transfer free energy as the difference between the solvation free energy of the solute going from the gas to the octanol phase, and the hydration free energy going from the gas to the water phase. The direct transfer free energy method that was run with YANK had not computed gas-to-water transfer free energies as previous work had done when computing log *P* and log *D* values and, while in principle this should be an unimportant methodological detail, we wanted to assess whether this choice had negatively impacted results. Thus, we ran the indirect solvation-based transfer free energy protocol described below.

The set of indirect solvation-based alchemical free energy calculations were run using OpenEye’s Orion cloud computing platform, also with the YANK software but with an alternate, fully automated workflow. The Orion workflow utilizes a very similar approach to that utilized above, except that it employs solvation free energy calculations for each molecule in each phase, rather than computing transfer free energies. Details of equilibration and simulation length are also different, as described below – with the largest difference being equilibration protocol.

On Orion, the input for each calculation was the target solute (SMILES) and the target solvent (SMILES), along with the temperature (298.15 K) and a guessed initial density for each solution (here 1.0 g/mL for solutes in water, 0.83 g/mL for octanol, to match experiment roughly). These settings are used on Orion, to prepare initial simulations via an internal Orion workflow based on that used in SolvationToolkit. The GAFF version used during parameterization on Orion was 1.8. In this Orion workflow, we also tested several additional potential tautomers for some molecules. For each molecule, we conducted a solvation free energy calculation of the solute in pure water and another in octanol. After parameterization, equilibration stages were run with OpenMM (version 7.2.2.dev-32bc79a) and free energy calculations were done with YANK (version 0.23.7 [115]). A cutoff for nonbonded interactions was set to 9 Å, electrostatic interactions were computed using PME, bonds involving hydrogen were constrained and HMR was used to allow for a 4 fs timestep.

The equilibration was carried out with OpenMM on Orion. The first step was 200 ps of NVT simulation with the solute heavy atoms harmonically restrained with 2.0 kcal/(mol·Å^2^) spring constants. The second step of equilibration was 200 ps of NPT simulation with harmonic heavy atom restraints with a 0.1 kcal/(mol·Å^2^) spring constant. These equilibrated structures were then used in YANK [61] simulations. The length of the YANK simulations were 5 ns for each replica. The electrostatic and steric interactions of the solute with the solvent were scaled using the same *λ* parameters listed in the transfer free energy protocol previously. The OpenEye workflow was also different in that it employed the ELF10 AM1-BCC charging engine (https://docs.eyesopen.com/toolkits/python/quacpactk/OEProtonClasses/OEAM1BCCELF10Charges.html, https://docs.eyesopen.com/applications/quacpac/theory/molcharge_theory.html), and only *syn* conformers of neutral carboxylic acids were retained for charging because, in OpenEye’s view, *anti* comformers result in incorrect charges dominated by strong internal interactions which are not well suited for MM applications. The only carboxylic acid studied was SM08, but the modification of the charging procedure in this case (relative to that employed in our direct solvation free energy approach) appears to have significantly impacted employed partial charges, likely for the better, as performance on SM08 was markedly different with this protocol.

#### 12.1.3 Empirical reference calculations

For all empirical calculations, the compounds were stripped of counter ions and neutralized. The pyridone tautomer of SM08 was used, as given, and as it is assumed to be the most stable tautomer.

The MOE/logP(o/w) model, the MOE/h_logP model, and the MOE/S_logP model are all available within the graphical modeling program MOE (MOE, available from the Chemical Computing Group, Montreal, www.chemcomp.com). The MOE/logP(o/w) model is based on 95 atom types, plus a few corrections for geminal halogens, 1-4 aromatic nitrogens, ethylene-glycol ethers, alkane carbons, and amino acids. The individual contributions were obtained from fitting to a data set of 1827 measurements, yielding an R^2^ of 0.931 and an RMSE of 0.393 (P. Labute, logP(o/w) model, unpublished).

The MOE/h_logP model uses 8 2D-descriptors derived from Extended Huckel Theory (the descriptors used are the sum of atomic EHT donor and acceptor strengths, the sum over log(1 + pi-bond order), the sum over log(1+ d-orbital bond order), the Gerber ring number and Gerber atomic surface area [116], and the number of hydrogens and number of hydrophobic carbons (carbons with no heteroatom within 3 bonds). The contributions of these descriptors were obtained by fitting to 1836 molecules yielding a model with an R^2^ of 0.084 and an RMSE of 0.59 (P. Labute, MOE h_mr, h_logP, and h_logS models, unpublished). The MOE program is available from the Chemical Computing Group, Montreal (www.chemcomp.com). The MOE/S_logP model is described in this reference [35]. In this model, 68 different atom types were defined based on element and nearest neighbors, e.g. 27 different carbon types or 14 different nitrogen types. Then the atomic contributions were determined by fitting to a training set of almost 10000 molecules.

The MoKa/logP methodology [MoKa-3.2.2, Molecular Discovery Ltd, London, www.moldiscovery.com] builds on a similar approach as the corresponding p*K*_a_ prediction [117]. The procedure starts by calculating molecular interaction fields based on the GRID force field on a large number of molecular fragments. The 3D energy fields of these fragments are then stored and used to recompute any molecule as a summation of appropriate 3D fragments. Therefore any molecule can be quickly approximated by 3D fields describing polar and hydrophobic interaction with water and n-octanol. From these fields, VolSurf descriptors are computed and used in a training scheme using a database of about 20000 compounds. From the training model, a final model is computed to make external predictions (G. Cruciani, personal communication).

#### 12.2 Supplementary figures and tables

**Table S1.**
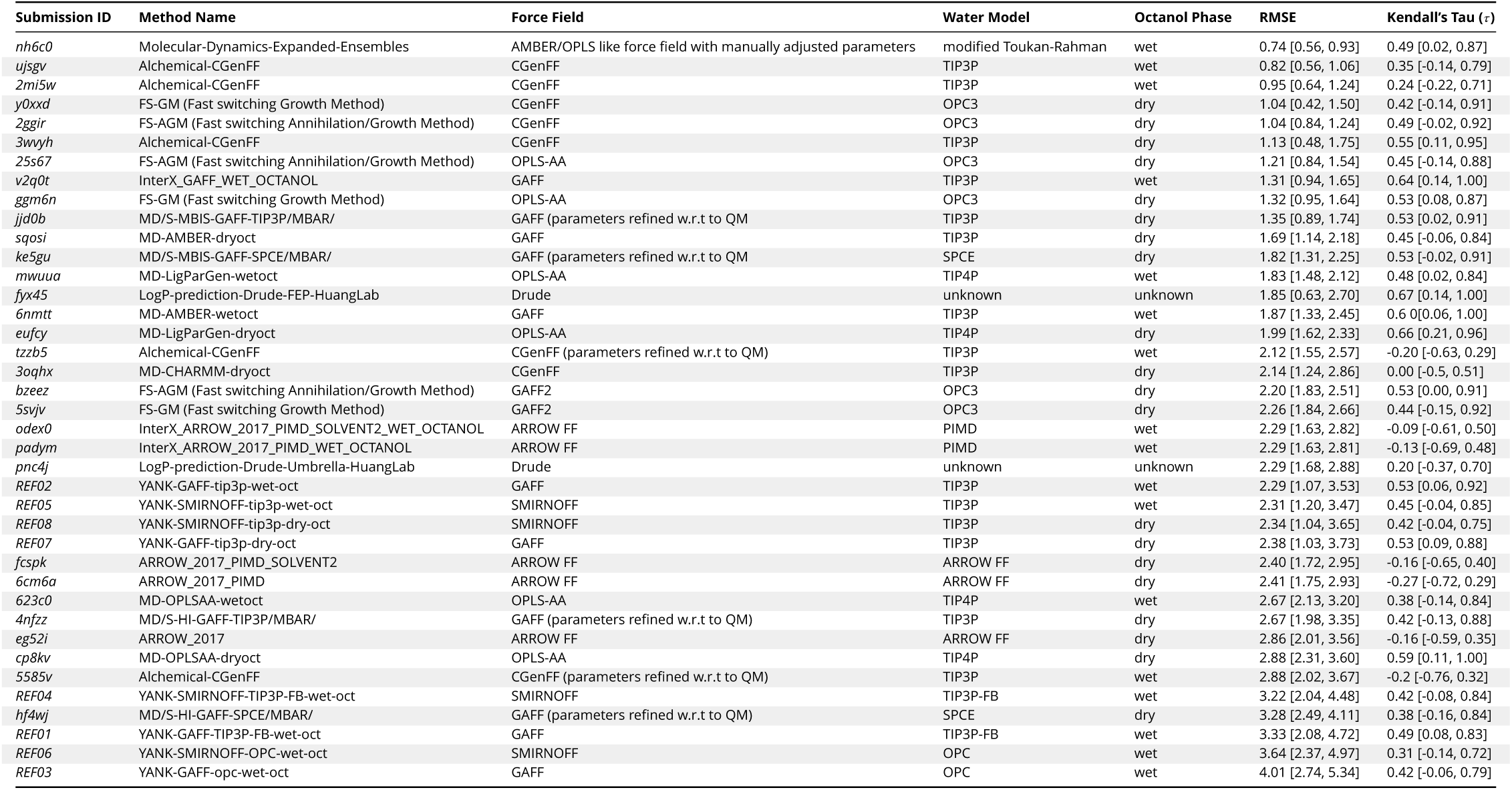
Method details of log *P* predictions with MM-based physical methods. Force fields, water models, and octanol phase choice are reported. A dry octanol phase indicates the octanol phase was treated as consisting of pure octanol. A wet octanol phase indicates the octanol phase was treated as a mixture of octanol and water. RMSE and Kendall’s Tau values are reported as mean and 95% confidence intervals. A CSV version of this table can be found in *SAMPL6-supplementary-documents.tar.gz*.

**Table S2.**
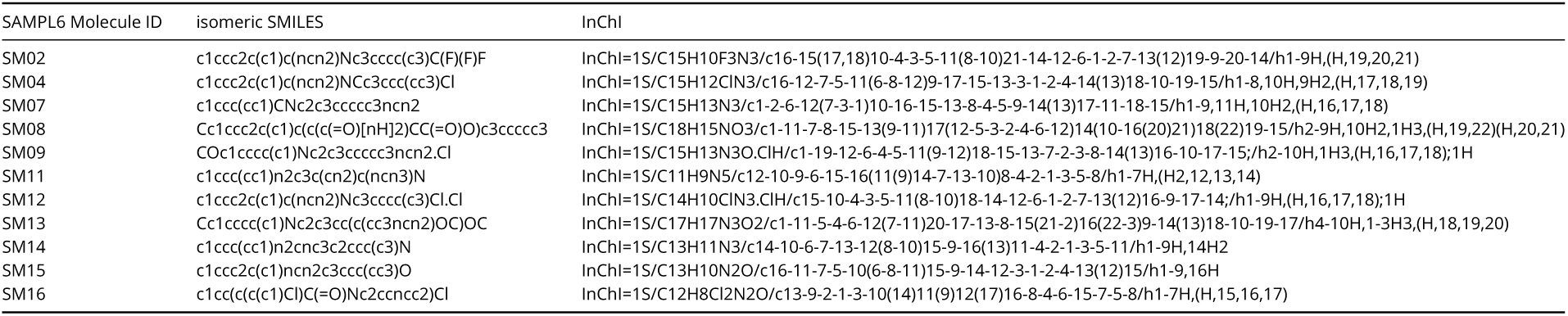
SMILES and InChI identifiers of SAMPL6 log *P* Challenge molecules. Experimental log *P* values can be found in a separate paper reporting measurements [9]. A CSV version of this table can be found in *SAMPL6-supplementary-documents.tar.gz*.

**Table S3.**
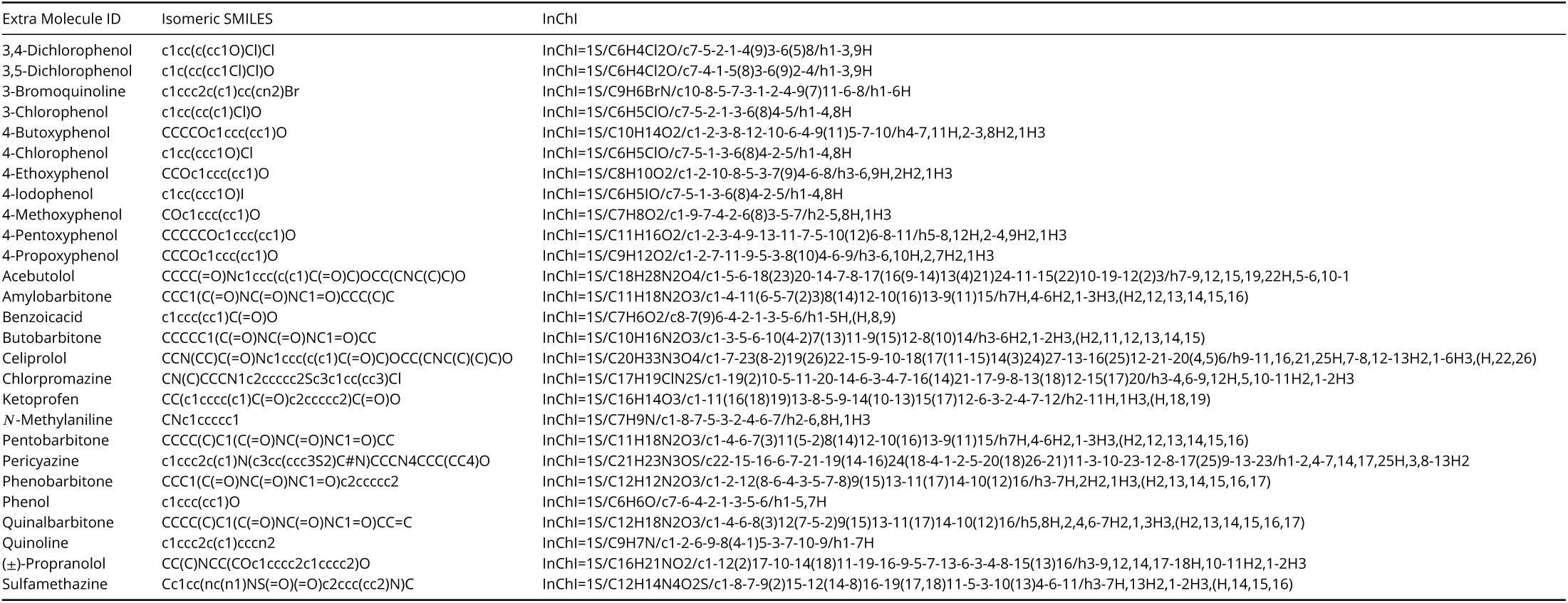
SMILES and InChI identifiers of extra molecules included in the evaluation of reference methods. A CSV version of this table can be found in *SAMPL6-supplementary-documents.tar.gz*. Experimental log *P* values can be found in a separate paper reporting measurements [108] and in machine readable format in https://github.com/samplchallenges/SAMPL6/blob/master/physical_properties/logP/analysis_of_extra_molecules/logP_experimental_values.csv.

**Table S4.**
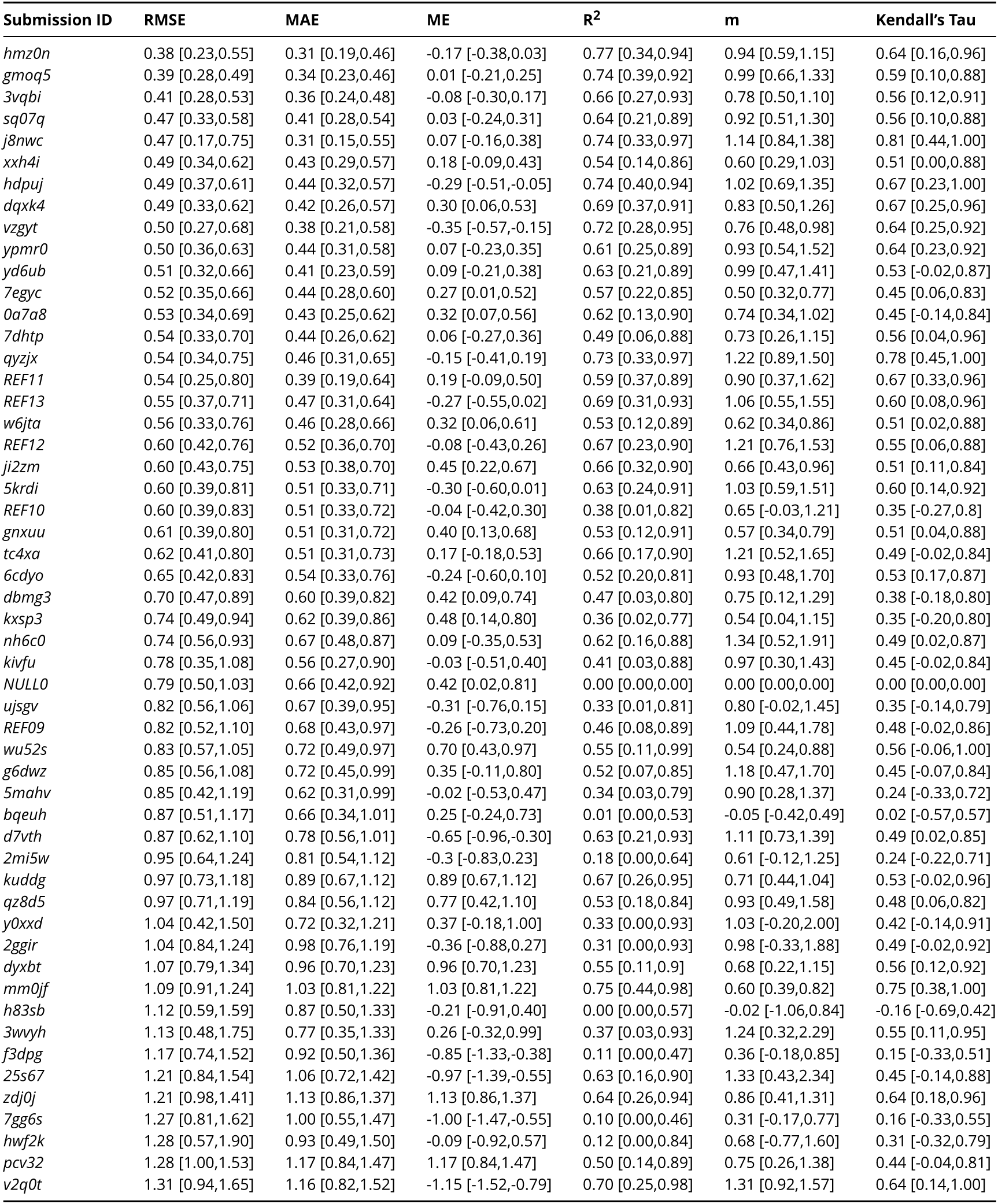
Evaluation statistics calculated for all methods. Methods are represented via their SAMPL6 submission IDs which can be cross referenced with Table 3 for method details. There are six error metrics reported: the root-mean-squared error (RMSE), mean absolute error (MAE), mean (signed) error (ME), coefficient of determination (R^2^), linear regression slope (m), and Kendall’s Rank Correlation Coefficient (*τ*). This table is ranked by increasing RMSE. A CSV version of this table can be found in *SAMPL6-supplementary-documents.tar.gz*.

**Table S5.**
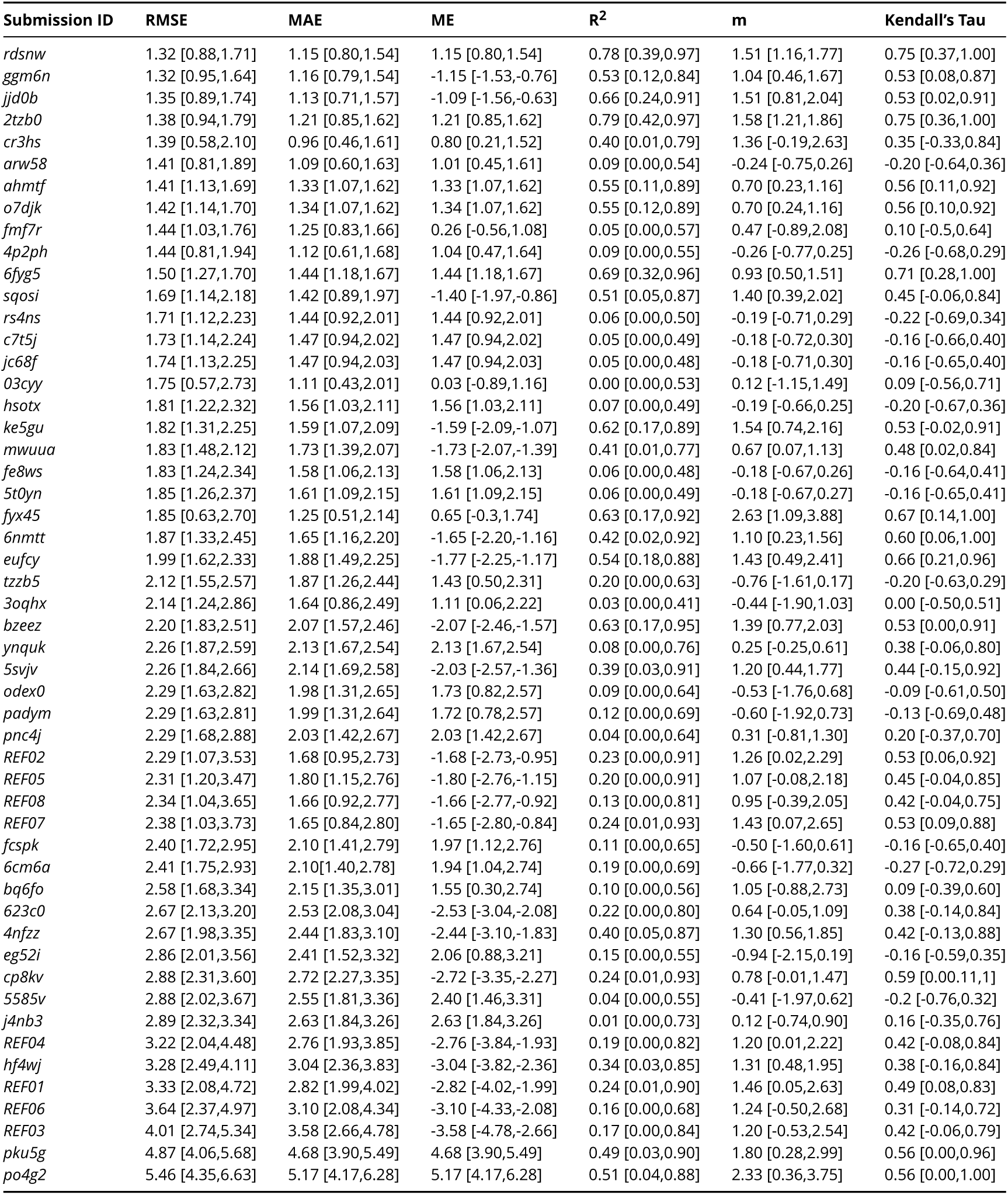
[Table S4 continued.] Evaluation statistics calculated for all methods. Methods are represented via their SAMPL6 submission IDs which can be cross referenced with Table 3 for method details. There are six error metrics reported: the root-mean-squared error (RMSE), mean absolute error (MAE), mean (signed) error (ME), coefficient of determination (R^2^), linear regression slope (m), and Kendall’s Rank Correlation Coefficient (*τ*). This table is ranked by increasing RMSE. A CSV version of this table can be found in *SAMPL6-supplementary-documents.tar.gz*.

**Figure S1.**
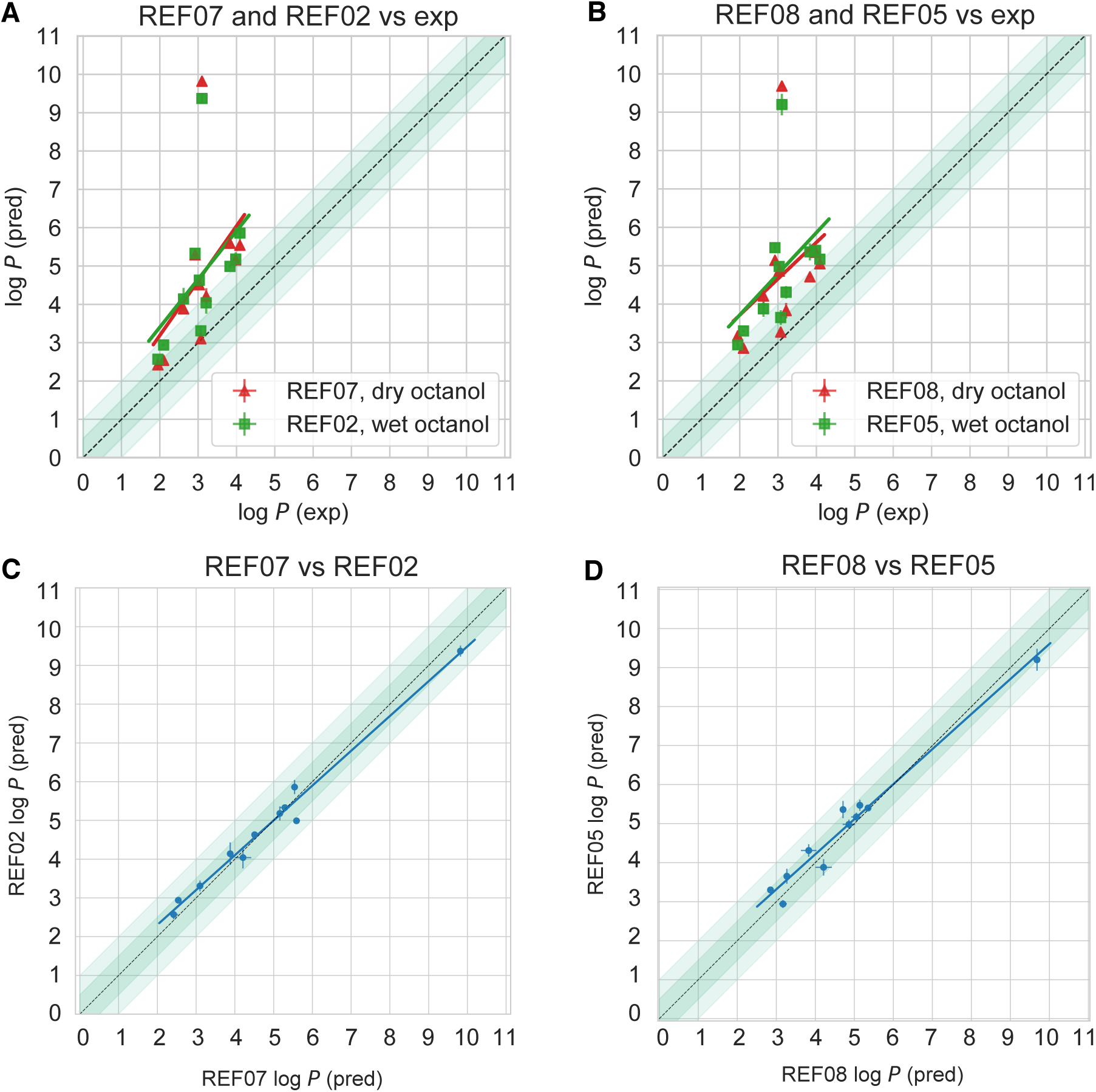
Varying the amount of water in the octanol phase has no significant effect on the predicted log *P* in reference calculations, as discussed in section 4.2.1. Comparison of predicted log *P* values to the experimental values using wet (27% water) and dry octanol phases and the (A) GAFF and (B) SMIRNOFF force field, from non-blinded reference calculations performed for this paper, shows no statistically significant difference in performance of methodologies. Comparison of the calculated log *P* using dry and wet octanol phases for (C) the GAFF force field and (D) the SMIRNOFF force field shows a small systematic difference.

**Table S6.**
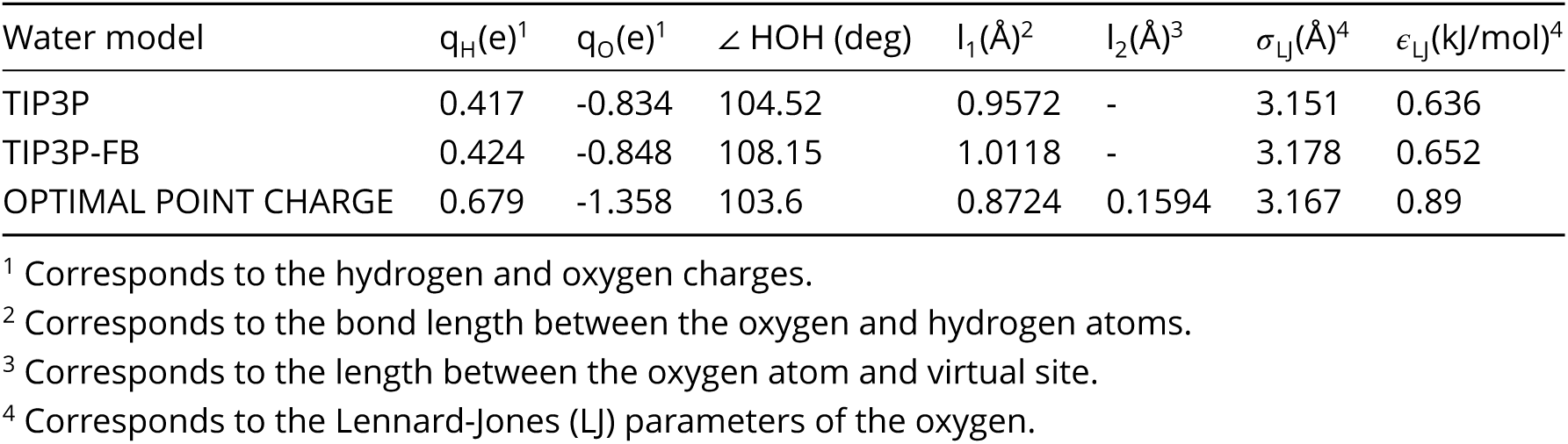
Comparison of force field parameters of the TIP3P, TIP3P-FB and OPC water models.

**Table S7.** Comparison of the charges assigned to the syn and anti conformation of SM08_micro011 in the DFE protocol.

| anti conformation |  |  |  | syn conformation |  |  |  |
| --- | --- | --- | --- | --- | --- | --- | --- |
| atom number | atom type | atom name | charge | atom number | atom type | atom name | charge |
| 1 | C1 | C1 | 0.1252 | 1 | C1 | C1 | -0.1205 |
| 2 | C2 | C2 | -0.0073 | 2 | C1 | C2 | -0.1322 |
| 3 | C2 | C3 | 0.0000 | 3 | C1 | C3 | -0.1322 |
| 4 | C2 | C4 | 0.0000 | 4 | C1 | C4 | -0.1112 |
| 5 | C2 | C5 | -0.0074 | 5 | C1 | C5 | -0.1112 |
| 6 | C2 | C6 | -0.0118 | 6 | C1 | C6 | -0.0742 |
| 7 | C2 | C7 | 0.0000 | 7 | C1 | C7 | -0.1856 |
| 8 | C2 | C8 | 0.0236 | 8 | C1 | C8 | -0.0653 |
| 9 | C2 | C9 | -0.0196 | 9 | C1 | C9 | -0.0839 |
| 10 | C2 | C10 | 0.3597 | 10 | C1 | C10 | -0.1199 |
| 11 | O1 | O1 | -0.2755 | 11 | C1 | C11 | -0.1160 |
| 12 | N1 | N1 | -0.1461 | 12 | C1 | C12 | 0.0947 |
| 13 | C1 | C11 | 0.0310 | 13 | C1 | C13 | 0.1024 |
| 14 | C2 | C12 | 0.3291 | 14 | C1 | C14 | -0.1731 |
| 15 | O1 | O2 | -0.1890 | 15 | C1 | C15 | 0.6987 |
| 16 | O2 | O3 | -0.2911 | 16 | C1 | C16 | 0.6458 |
| 17 | C2 | C13 | -0.0118 | 17 | C2 | C17 | -0.0498 |
| 18 | C2 | C14 | 0.0000 | 18 | C2 | C18 | -0.0780 |
| 19 | C2 | C15 | 0.0000 | 19 | N1 | N1 | -0.4371 |
| 20 | C2 | C16 | 0.0000 | 20 | O1 | O1 | -0.6386 |
| 21 | C2 | C17 | 0.0000 | 21 | O1 | O2 | -0.5474 |
| 22 | C2 | C18 | 0.0000 | 22 | O2 | O3 | -0.6145 |
| 23 | H1 | H1 | -0.0393 | 23 | H1 | H1 | 0.1352 |
| 24 | H1 | H2 | -0.0393 | 24 | H1 | H2 | 0.1377 |
| 25 | H1 | H3 | -0.0393 | 25 | H1 | H3 | 0.1377 |
| 26 | H2 | H4 | 0.0000 | 26 | H1 | H4 | 0.1459 |
| 27 | H2 | H5 | 0.0000 | 27 | H1 | H5 | 0.1459 |
| 28 | H2 | H6 | 0.0000 | 28 | H1 | H6 | 0.1381 |
| 29 | H3 | H7 | 0.0865 | 29 | H1 | H7 | 0.1406 |
| 30 | H1 | H8 | -0.0393 | 30 | H1 | H8 | 0.1469 |
| 31 | H1 | H9 | -0.0393 | 31 | H2 | H9 | 0.0455 |
| 32 | H4 | H10 | 0.2010 | 32 | H2 | H10 | 0.0455 |
| 33 | H2 | H11 | 0.0000 | 33 | H2 | H11 | 0.0455 |
| 34 | H2 | H12 | 0.0000 | 34 | H2 | H12 | 0.1028 |
| 35 | H2 | H13 | 0.0000 | 35 | H2 | H13 | 0.1028 |
| 36 | H2 | H14 | 0.0000 | 36 | H3 | H14 | 0.3362 |
| 37 | H2 | H15 | 0.0000 | 37 | H4 | H15 | 0.4428 |

**Figure S2.**
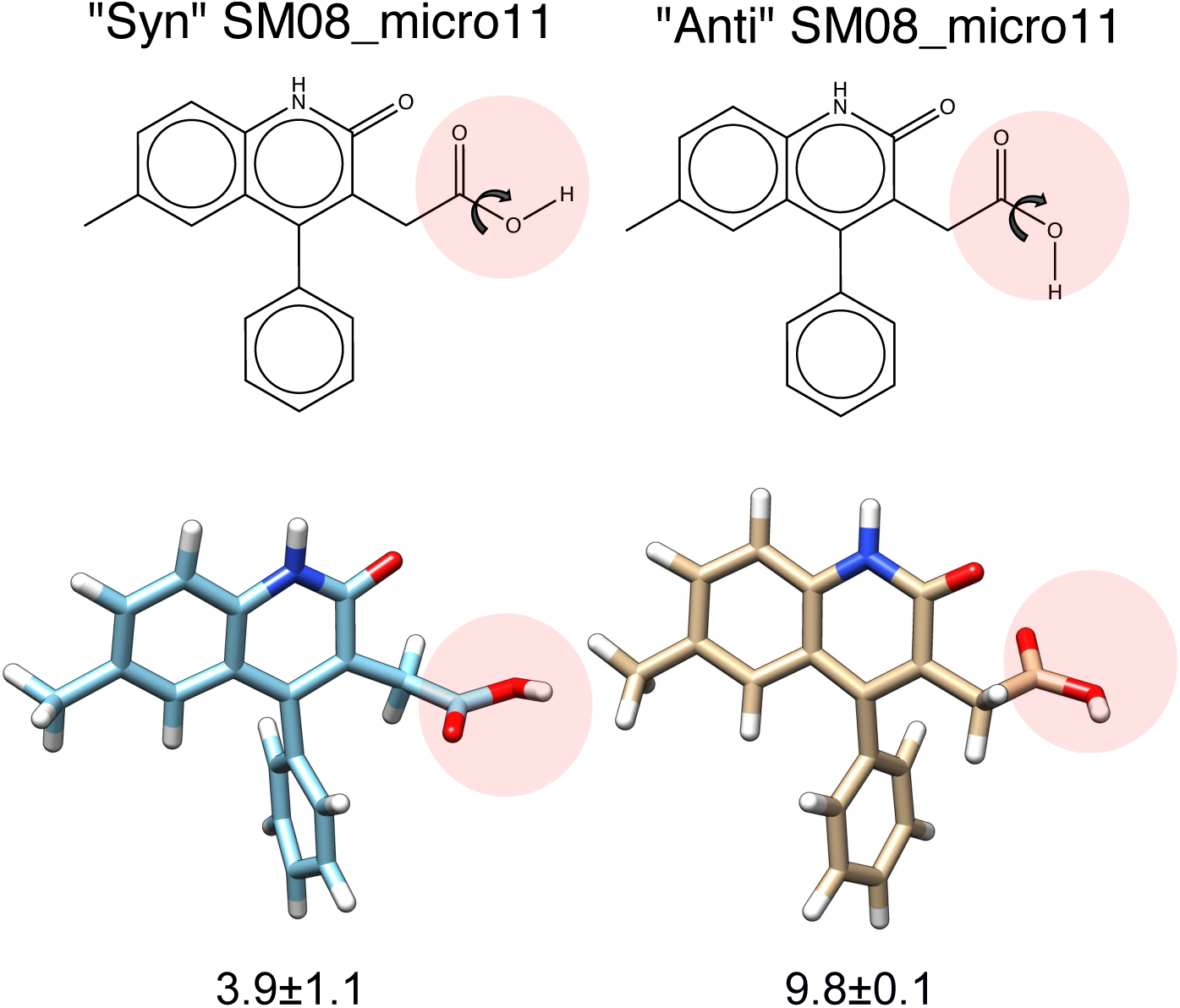
Shown here are the 2- and 3D structures of SM08_micro011 with the carboxylic acid in “anti” and “syn” conformation. The dihedral angle is indicated by the arrow around the carbon and oxygen atom. The calculated log *P* is included for comparison. The charges pertaining to each conformation are listed in Figure S7 and transition data is available in Figure S3

**Figure S3.**
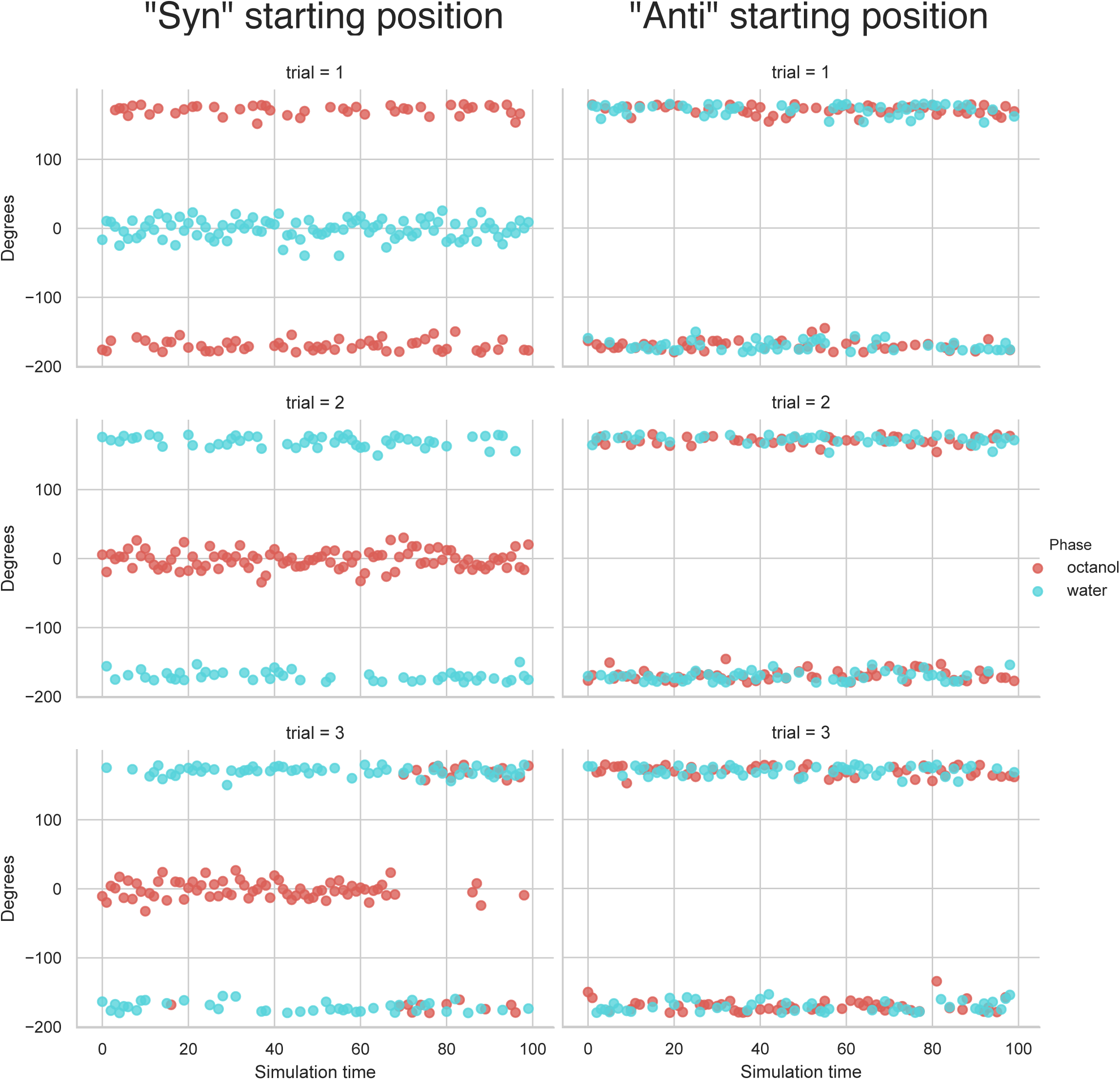
For the DFE method, the starting conformation impacts the number of C-O dihedral transitions for SM08_micro011, influencing sampling. Here is the transition data for the C-O dihedral in Figure S2, with charges listed in Table S7, for the DFE method (run in triplicate). In the “anti starting position” the torsion remains “anti” throughout the simulation, while the “syn starting position” allows transitions.

## References

[1] Rustenburg AS, Dancer J, Lin B, Feng JA, Ortwine DF, Mobley DL, Chodera JD. Measuring Experimental Cyclohexane-Water Distribution Coefficients for the SAMPL5 Challenge. Journal of Computer-Aided Molecular Design. 2016 Nov; 30(11):945–958. doi: 10.1007/s10822-016-9971-7.

[2] Bannan CC, Burley KH, Chiu M, Shirts MR, Gilson MK, Mobley DL. Blind Prediction of Cyclohexane–Water Distribution Coefficients from the SAMPL5 Challenge. Journal of Computer-Aided Molecular Design. 2016 Nov; 30(11):927–944. doi: 10.1007/s10822-016-9954-8.

[3] Kollman PA. Advances and Continuing Challenges in Achieving Realistic and Predictive Simulations of the Properties of Organic and Biological Molecules. Accounts of Chemical Research. 1996 Jan; 29(10):461–469. doi: 10.1021/ar9500675.

[4] Best SA, Merz KM, Reynolds CH. Free Energy Perturbation Study of Octanol/Water Partition Coefficients: Comparison with Continuum GB/SA Calculations. The Journal of Physical Chemistry B. 1999 Jan; 103(4):714–726. doi: 10.1021/jp984215v.

[5] Chen B, Siepmann JI. Microscopic Structure and Solvation in Dry and Wet Octanol. The Journal of Physical Chemistry B. 2006 Mar; 110(8):3555–3563. doi: 10.1021/jp0548164.

[6] Lyubartsev AP, Jacobsson SP, Sundholm G, Laaksonen A. Solubility of Organic Compounds in Water/Octanol Systems. A Expanded Ensemble Molecular Dynamics Simulation Study of Log P Parameters. The Journal of Physical Chemistry B. 2001 Aug; 105(32):7775–7782. doi: 10.1021/jp0036902.

[7] DeBolt SE, Kollman PA. Investigation of Structure, Dynamics, and Solvation in 1-Octanol and Its Water-Saturated Solution: Molecular Dynamics and Free-Energy Perturbation Studies. J Am Chem Soc. 1995 May; 117(19):5316–5340. doi: 10.1021/ja00124a015.

[8] Isik M, Levorse D, Rustenburg AS, Ndukwe IE, Wang H, Wang X, Reibarkh M, Martin GE, Makarov AA, Mobley DL, Rhodes T, Chodera JD. pKa Measurements for the SAMPL6 Prediction Challenge for a Set of Kinase Inhibitor-like Fragments. Journal of Computer-Aided Molecular Design. 2018 Oct; 32(10):1117–1138. doi: 10.1007/s10822-018-0168-0.

[9] Isik M, Levorse D, Mobley DL, Rhodes T, Chodera JD. Octanol–Water Partition Coefficient Measurements for the SAMPL6 Blind Prediction Challenge. Journal of Computer-Aided Molecular Design. 2019 Dec; doi: 10.1007/s10822-019-00271-3.

[10] Marenich AV, Cramer CJ, Truhlar DG. Universal Solvation Model Based on Solute Electron Density and on a Continuum Model of the Solvent Defined by the Bulk Dielectric Constant and Atomic Surface Tensions. J Phys Chem B. 2009 May; 113(18):6378–6396. doi: 10.1021/jp810292n.

[11] Marenich AV, Cramer CJ, Truhlar DG. Performance of SM6, SM8, and SMD on the SAMPL1 Test Set for the Prediction of Small-Molecule Solvation Free Energies. J Phys Chem B. 2009 Apr; 113(14):4538–4543. doi: 10.1021/jp809094y.

[12] Ribeiro RF, Marenich AV, Cramer CJ, Truhlar DG. Prediction of SAMPL2 Aqueous Solvation Free Energies and Tautomeric Ratios Using the SM8, SM8AD, and SMD Solvation Models. J Comput Aided Mol Des. 2010 Apr; 24(4):317–333. doi: 10.1007/s10822-010-9333-9.

[13] Marenich AV, Cramer CJ, Truhlar DG. Generalized Born Solvation Model SM12. J Chem Theory Comput. 2013 Jan; 9(1):609–620. doi: 10.1021/ct300900e.

[14] Loschen C, Reinisch J, Klamt A. COSMO-RS Based Predictions for the SAMPL6 logP Challenge. Journal of Computer-Aided Molecular Design. 2019 Nov; doi: 10.1007/s10822-019-00259-z.

[15] Klamt A, Eckert F, Reinisch J, Wichmann K. Prediction of Cyclohexane-Water Distribution Coefficients with COSMO-RS on the SAMPL5 Data Set. Journal of Computer-Aided Molecular Design. 2016 Nov; 30(11):959–967. doi: 10.1007/s10822-016-9927-y.

[16] Klamt A, Eckert F, Diedenhofen M. Prediction of the Free Energy of Hydration of a Challenging Set of Pesticide-Like Compounds †. The Journal of Physical Chemistry B. 2009 Apr; 113(14):4508–4510. doi: 10.1021/jp805853y.

[17] Klamt A. Conductor-like Screening Model for Real Solvents: A New Approach to the Quantitative Calculation of Solvation Phenomena. The Journal of Physical Chemistry. 1995 Feb; 99(7):2224–2235. doi: 10.1021/j100007a062.

[18] Klamt A, Jonas V, Bürger T, Lohrenz JCW. Refinement and Parametrization of COSMO-RS. The Journal of Physical Chemistry A. 1998 Jun; 102(26):5074–5085. doi: 10.1021/jp980017s.

[19] Tielker N, Tomazic D, Heil J, Kloss T, Ehrhart S, Güssregen S, Schmidt KF, Kast SM. The SAMPL5 Challenge for Embedded-Cluster Integral Equation Theory: Solvation Free Energies, Aqueous pK a, and Cyclohexane–Water Log D. Journal of Computer-Aided Molecular Design. 2016 Nov; 30(11):1035–1044. doi: 10.1007/s10822-016-9939-7.

[20] Tielker N, Eberlein L, Güssregen S, Kast SM. The SAMPL6 Challenge on Predicting Aqueous pKa Values from EC-RISM Theory. Journal of Computer-Aided Molecular Design. 2018 Oct; 32(10):1151–1163. doi: 10.1007/s10822-018-0140-z.

[21] Luchko T, Blinov N, Limon GC, Joyce KP, Kovalenko A. SAMPL5: 3D-RISM Partition Coefficient Calculations with Partial Molar Volume Corrections and Solute Conformational Sampling. J Comput Aided Mol Des. 2016 Nov; 30(11):1115–1127. doi: 10.1007/s10822-016-9947-7.

[22] Tielker N, Tomazic D, Eberlein L, Güssregen S, Kast SM. The SAMPL6 challenge on predicting octanol-water partition coefficients from ECRISM theory. Journal of Computer-Aided Molecular Design. 2020; (SAMPL6 Part II Special Issue).

[23] Beglov D, Roux B. An Integral Equation To Describe the Solvation of Polar Molecules in Liquid Water. J Phys Chem B. 1997 Sep; 101(39):7821–7826. doi: 10.1021/jp971083h.

[24] Kovalenko A, Hirata F. Three-Dimensional Density Profiles of Water in Contact with a Solute of Arbitrary Shape: A RISM Approach. Chemical Physics Letters. 1998 Jun; 290(1-3):237–244. doi: 10.1016/S0009-2614(98)00471-0.

[25] Tielker N, Eberlein L, Güssregen S, Kast SM. The SAMPL6 Challenge on Predicting Aqueous pKa Values from EC-RISM Theory. J Comput Aid Mol Des. 2018 Oct; 32(10):1151–1163. doi: 10.1007/s10822-018-0140-z.

[26] Kloss T, Heil J, Kast SM. Quantum Chemistry in Solution by Combining 3D Integral Equation Theory with a Cluster Embedding Approach. The Journal of Physical Chemistry B. 2008 Apr; 112(14):4337–4343. doi: 10.1021/jp710680m.

[27] Darve E, Pohorille A. Calculating free energies using average force. The Journal of Chemical Physics. 2001; 115(20):9169–9183. https://doi.org/10.1063/1.1410978, doi: 10.1063/1.1410978.

[28] Comer J, Gumbart JC, Hénin J, Lelièvre T, Pohorille A, Chipot C. The Adaptive Biasing Force Method: Everything You Always Wanted To Know but Were Afraid To Ask. The Journal of Physical Chemistry B. 2015 Jan; 119(3):1129–1151. doi: 10.1021/jp506633n.

[29] Nieto-Draghi C, Fayet G, Creton B, Rozanska X, Rotureau P, de Hemptinne JC, Ungerer P, Rousseau B, Adamo C. A General Guidebook for the Theoretical Prediction of Physicochemical Properties of Chemicals for Regulatory Purposes. Chemical Reviews. 2015 Dec; 115(24):13093–13164. doi: 10.1021/acs.chemrev.5b00215.

[30] Mannhold R, Poda GI, Ostermann C, Tetko IV. Calculation of Molecular Lipophilicity: State-of-the-Art and Comparison of LogP Methods on More than 96,000 Compounds. Journal of Pharmaceutical Sciences. 2009 Mar; 98(3):861–893. doi: 10.1002/jps.21494.

[31] D Eros LOKTNGA I Kovesdi, Keri G. Reliability of logP Predictions Based on Calculated Molecular Descriptors: A Critical Review. Current Medicinal Chemistry. 2002; 9(20):1819–1829. http://www.eurekaselect.com/node/64184/article, doi: 10.2174/0929867023369042.

[32] Ghose AK, Crippen GM. Atomic Physicochemical Parameters for Three-Dimensional Structure-Directed Quantitative Structure-Activity Relationships I. Partition Coefficients as a Measure of Hydrophobicity. Journal of Computational Chemistry. 1986; 7(4):565–577. doi: 10.1002/jcc.540070419.

[33] Ghose AK, Crippen GM. Atomic Physicochemical Parameters for Three-Dimensional-Structure-Directed Quantitative Structure-Activity Relationships. 2. Modeling Dispersive and Hydrophobic Interactions. J Chem Inf Comput Sci. 1987 Feb; 27(1):21–35. doi: 10.1021/ci00053a005.

[34] Ghose AK, Viswanadhan VN, Wendoloski JJ. Prediction of Hydrophobic (Lipophilic) Properties of Small Organic Molecules Using Fragmental Methods: An Analysis of ALOGP and CLOGP Methods. The Journal of Physical Chemistry A. 1998 May; 102(21):3762–3772. doi: 10.1021/jp980230o.

[35] Wildman SA, Crippen GM. Prediction of Physicochemical Parameters by Atomic Contributions. J Chem Inf Comput Sci. 1999 Sep; 39(5):868–873. doi: 10.1021/ci990307l.

[36] Wang R, Fu Y, Lai L. A New Atom-Additive Method for Calculating Partition Coefficients. Journal of Chemical Information and Computer Sciences. 1997 May; 37(3):615–621. doi: 10.1021/ci960169p.

[37] Wang R, Gao Y, Lai L. Calculating Partition Coefficient by Atom-Additive Method. Perspect Drug Disc. 2000; 19(1):47–66. doi: 10.1023/A:1008763405023.

[38] Cheng T, Zhao Y, Li X, Lin F, Xu Y, Zhang X, Li Y, Wang R, Lai L. Computation of Octanol-Water Partition Coefficients by Guiding an Additive Model with Knowledge. Journal of Chemical Information and Modeling. 2007 Nov; 47(6):2140–2148. doi: 10.1021/ci700257y.

[39] Leo A J, Hoekman D. Calculating Log P(Oct) with No Missing Fragments;The Problem of Estimating New Interaction Parameters. Perspectives in Drug Discovery and Design. 2000 Jun; 18(1):19–38. doi: 10.1023/A:1008739110753.

[40] Leo A J. Calculating log Poct from structures. Chemical Reviews. 1993; 93(4):1281–1306.

[41] Leo A. The Octanol–Water Partition Coefficient of Aromatic Solutes: The Effect of Electronic Interactions, Alkyl Chains, Hydrogen Bonds, and Ortho-Substitution. J Chem Soc, Perkin Trans 2. 1983; (6):825–838. doi: 10.1039/P29830000825.

[42] Sangster J. Octanol-water partition coefficients of simple organic compounds. Journal of Physical and Chemical Reference Data. 1989; 18(3):1111–1229.

[43] Klopman G, Li JY, Wang S, Dimayuga M. Computer Automated Log P Calculations Based on an Extended Group Contribution Approach. Journal of Chemical Information and Modeling. 1994 Jul; 34(4):752–781. doi: 10.1021/ci00020a009.

[44] Petrauskas AA, Kolovanov EA. ACD/Log P Method Description. Persect Drug Discov. 2000 Sep; 19(1):99–116. doi: 10.1023/A:1008719622770.

[45] Meylan WM, Howard PH. Atom/Fragment Contribution Method for Estimating Octanol–Water Partition Coefficients. Journal of Pharmaceutical Sciences. 1995 Jan; 84(1):83–92. doi: 10.1002/jps.2600840120.

[46] Moriguchi I, Hirono S, Liu Q, Nakagome I, Matsushita Y. Simple Method of Calculating Octanol/Water Partition Coefficient. Chem Pharm Bull. 1992; 40(1):127–130. doi: 10.1248/cpb.40.127.

[47] Gombar VK, Enslein K. Assessment of N-Octanol/Water Partition Coefficient: When Is the Assessment Reliable? J Chem Inf Comput Sci. 1996 Jan; 36(6):1127–1134. doi: 10.1021/ci960028n.

[48] Wishart DS, Knox C, Guo AC, Shrivastava S, Hassanali M, Stothard P, Chang Z, Woolsey J. DrugBank: A Comprehensive Resource for in Silico Drug Discovery and Exploration. Nucleic Acids Res. 2006 Jan; 34(Database issue):D668–672. doi: 10.1093/nar/gkj067.

[49] Pence HE, Williams A. ChemSpider: An Online Chemical Information Resource. J Chem Educ. 2010 Nov; 87(11):1123–1124. doi: 10.1021/ed100697w.

[50] NCI Open Database, August 2006 Release;. https://cactus.nci.nih.gov/download/nci/.

[51] Enhanced NCI Database Browser 2.2;. https://cactus.nci.nih.gov/ncidb2.2/.

[52] SRC’s PHYSPROP Database;. https://www.srcinc.com/what-we-do/environmental/scientific-databases.html.

[53] OEDepict Toolkit 2017.Feb.1;. OpenEye Scientific Software, Santa Fe, NM. http://www.eyesopen.com.

[54] Mobley DL, Isik M, Paluch A, Loschen C, Tielker N, Vöhringer-Martinez E, Nikitin A, The SAMPL6 LogP Virtual Workshop. Zenodo; 2019. https://doi.org/10.5281/zenodo.3518862, doi: 10.5281/zenodo.3518862.

[55] Mobley DL, Isik M, Paluch A, Loschen C, Tielker N, Vöhringer-Martinez E, Nikitin A, The SAMPL6 LogP Virtual Workshop GitHub Repository for Presentation Slides; 2019. https://github.com/choderalab/SAMPL6-logP-challenge-virtual-workshop.

[56] Mobley DL, SAMPL: Its present and future, and some work on the logP challenge. Zenodo; 2019. https://doi.org/10.5281/zenodo.3376196, doi: 10.5281/zenodo.3376196.

[57] Isik M, SAMPL6 Part II Partition Coefficient Challenge Overview. Zenodo; 2019. https://doi.org/10.5281/zenodo.3386592, doi: 10.5281/zenodo.3386592.

[58] Lang BE. Solubility of Water in Octan-1-Ol from (275 to 369) K. Journal of Chemical & Engineering Data. 2012 Aug; 57(8):2221–2226. doi: 10.1021/je3001427.

[59] Leo A, Hansch C, Elkins D. Partition Coefficients and Their Uses. Chem Rev. 1971 Dec; 71(6):525–616. doi: 10.1021/cr60274a001.

[60] Mobley DL, Wymer KL, Lim NM, Guthrie JP. Blind Prediction of Solvation Free Energies from the SAMPL4 Challenge. Journal of Computer-Aided Molecular Design. 2014 Mar; 28(3):135–150. doi: 10.1007/s10822-014-9718-2.

[61] Andrea A, Grinaway P, Parton D, Shirts M, Wang K, Eastman P, Friedrichs M, Pande V, Branson K, Mobley D, Chodera J. YANK: A GPU-Accelerated Platform for Alchemical Free Energy Calculations..;.

[62] Wang K, Chodera JD, Yang Y, Shirts MR. Identifying ligand binding sites and poses using GPU-accelerated Hamiltonian replica exchange molecular dynamics. Journal of computer-aided molecular design. 2013; 27(12):989–1007.

[63] Rizzi A, Chodera J, Naden L, Beauchamp K, Albanese S, Grinaway P, Rustenburg B, ajsilveira, Saladi S, Boehm K, choderalab/yank: 0.24.0 - Experimental support for online status files. Zenodo; 2019. https://doi.org/10.5281/zenodo.2577832, doi: 10.5281/zenodo.2577832.

[64] Eastman P, Pande VS. Efficient Nonbonded Interactions for Molecular Dynamics on a Graphics Processing Unit. Journal of Computational Chemistry. 2009; p. NA–NA. doi: 10.1002/jcc.21413.

[65] Eastman P, Pande V. OpenMM: A Hardware-Independent Framework for Molecular Simulations. Comput Sci Eng. 2010 Jul; 12(4):34–39. doi: 10.1109/MCSE.2010.27.

[66] Eastman P, Friedrichs MS, Chodera JD, Radmer RJ, Bruns CM, Ku JP, Beauchamp KA, Lane TJ, Wang LP, Shukla D, Tye T, Houston M, Stich T, Klein C, Shirts MR, Pande VS. OpenMM 4: A Reusable, Extensible, Hardware Independent Library for High Performance Molecular Simulation. Journal of Chemical Theory and Computation. 2013 Jan; 9(1):461–469. doi: 10.1021/ct300857j.

[67] Wang J, Wolf RM, Caldwell JW, Kollman PA, Case DA. Development and Testing of a General Amber Force Field. Journal of Computational Chemistry. 2004 Jul; 25(9):1157–1174. doi: 10.1002/jcc.20035.

[68] Mobley DL, Bannan CC, Rizzi A, Bayly CI, Chodera JD, Lim VT, Lim NM, Beauchamp KA, Shirts MR, Gilson MK, Eastman PK. Open Force Field Consortium: Escaping Atom Types Using Direct Chemical Perception with SMIRNOFF v0.1. bioRxiv. 2018 Mar; doi: 10.1101/286542.

[69] Jorgensen WL, Chandrasekhar J, Madura JD, Impey RW, Klein ML. Comparison of Simple Potential Functions for Simulating Liquid Water. The Journal of Chemical Physics. 1983 Jul; 79(2):926–935. doi: 10.1063/1.445869.

[70] Wang LP, Martinez TJ, Pande VS. Building Force Fields: An Automatic, Systematic, and Reproducible Approach. J Phys Chem Lett. 2014 Jun; 5(11):1885–1891. doi: 10.1021/jz500737m.

[71] Izadi S, Anandakrishnan R, Onufriev AV. Building Water Models: A Different Approach. The Journal of Physical Chemistry Letters. 2014 Nov; 5(21):3863–3871. doi: 10.1021/jz501780a.

[72] Shultz MD. Two Decades under the Influence of the Rule of Five and the Changing Properties of Approved Oral Drugs: Miniperspective. J Med Chem. 2019 Feb; 62(4):1701–1714. doi: 10.1021/acs.jmedchem.8b00686.

[73] Procacci P, Guarnieri G. SAMPL6 Blind Predictions of Water-Octanol Partition Coefficients Using Nonequilibrium Alchemical Approaches. Journal of Computer-Aided Molecular Design. 2019 Oct; doi: 10.1007/s10822-019-00233-9.

[74] Riquelme M, Vöhringer-Martinez E. SAMPL6 Octanol–Water Partition Coefficients from Alchemical Free Energy Calculations with MBIS Atomic Charges. Journal of Computer-Aided Molecular Design. 2020 Jan; doi: 10.1007/s10822-020-00281-6.

[75] Patel P, Kuntz DM, Jones MR, Brooks B, Wilson A. SAMPL6 LogP Challenge: Machine Learning and Quantum Mechanical Approaches. Journal of Computer-Aided Molecular Design. 2020; (SAMPL6 Part II Special Issue).

[76] Ouimet JA, Paluch AS. Predicting Octanol/Water Partition Coefficients for the SAMPL6 Challenge Using the SM12, SM8, and SMD Solvation Models. Journal of Computer-Aided Molecular Design. 2020; (SAMPL6 Part II Special Issue).

[77] Lui R, Guan D, Matthews S. A Comparison of Molecular Representations for Lipophilicity Quantitative Structure–Property Relationships with Results from the SAMPL6 logP Prediction Challenge. Journal of Computer-Aided Molecular Design. 2020 Jan; doi: 10.1007/s10822-020-00279-0.

[78] Jones MR, Brooks BR. Quantum Chemical Predictions of Water-Octanol Partition Coefficients Applied to the SAMPL6 Blind Challenge. Journal of Computer-Aided Molecular Design. 2020; (SAMPL6 Part II Special Issue).

[79] Guan D, Lui R, Matthews S. LogP Prediction Performance with the SMD Solvation Model and the M06 Density Functional Family for SAMPL6 Blind Prediction Challenge Molecules. Journal of Computer-Aided Molecular Design. 2020 Jan; doi: 10.1007/s10822-020-00278-1.

[80] Arslan E, Findik BK, Aviyente V. A Blind SAMPL6 Challenge: Insight into the Octanol-Water Partition Coefficients of Drug-like Molecules via a DFT Approach. Journal of Computer-Aided Molecular Design. 2020 Jan; doi: 10.1007/s10822-020-00284-3.

[81] Fan S, Iorga BI, Beckstein O. Prediction of Octanol-Water Partition Coefficients for the SAMPL6-$$\log P$$logP Molecules Using Molecular Dynamics Simulations with OPLS-AA, AMBER and CHARMM Force Fields. Journal of Computer-Aided Molecular Design. 2020 Jan; doi: 10.1007/s10822-019-00267-z.

[82] Wang S, Riniker S. Use of Molecular Dynamics Fingerprints (MDFPs) in SAMPL6 Octanol–Water Log P Blind Challenge. Journal of Computer-Aided Molecular Design. 2019 Nov; doi: 10.1007/s10822-019-00252-6.

[83] Krämer A, Hudson PS, Jones MR, Brooks BR. Multi-Phase Boltzmann Weighting: Accounting for Local Inhomogeneity in Molecular Simulations of Water-Octanol Partition Coefficients. Journal of Computer-Aided Molecular Design. 2020; (SAMPL6 Part II Special Issue).

[84] Nikitin A. Non-Zero Lennard-Jones Parameters for the Toukan–Rahman Water Model: More Accurate Calculations of the Solvation Free Energy of Organic Substances. Journal of Computer-Aided Molecular Design. 2019 Nov; doi: 10.1007/s10822-019-00256-2.

[85] Zamora WJ, Pinheiro S, German K, Ràfols C, Curutchet C, Luque FJ. Prediction of the N-Octanol/Water Partition Coefficients in the SAMPL6 Blind Challenge from MST Continuum Solvation Calculations. Journal of Computer-Aided Molecular Design. 2019 Nov; doi: 10.1007/s10822-019-00262-4.

[86] Nikitin A, Milchevskiy Y, Lyubartsev A. AMBER-Ii: New Combining Rules and Force Field for Perfluoroalkanes. The Journal of Physical Chemistry B. 2015 Nov; 119(46):14563–14573. doi: 10.1021/acs.jpcb.5b07233.

[87] Lyubartsev AP, Martsinovski AA, Shevkunov SV, Vorontsov-Velyaminov PN. New Approach to Monte Carlo Calculation of the Free Energy: Method of Expanded Ensembles. Journal of chemical physics. 1992; 96(3):1776–1783. doi: 10.1063/1.462133.

[88] Vanommeslaeghe K, Hatcher E, Acharya C, Kundu S, Zhong S, Shim J, Darian E, Guvench O, Lopes P, Vorobyov I, Mackerell AD. CHARMM General Force Field: A Force Field for Drug-like Molecules Compatible with the CHARMM All-Atom Additive Biological Force Fields. Journal of Computational Chemistry. 2009; p. NA–NA. doi: 10.1002/jcc.21367.

[89] Zwanzig RW. High-Temperature Equation of State by a Perturbation Method. I. Nonpolar Gases. The Journal of Chemical Physics. 1954; 22(8):1420–1426. https://doi.org/10.1063/1.1740409, doi: 10.1063/1.1740409.

[90] Bennett CH. Efficient Estimation of Free Energy Differences from Monte Carlo Data. Journal of Computational Physics. 1976 Oct; 22(2):245–268. doi: 10.1016/0021-9991(76)90078-4.

[91] Kirkwood JG. Statistical Mechanics of Fluid Mixtures. The Journal of Chemical Physics. 1935; 3(5):300–313. https://doi.org/10.1063/1.1749657, doi: 10.1063/1.1749657.

[92] Procacci P, Cardelli C. Fast Switching Alchemical Transformations in Molecular Dynamics Simulations. Journal of Chemical Theory and Computation. 2014 Jul; 10(7):2813–2823. doi: 10.1021/ct500142c.

[93] Jarzynski C. Nonequilibrium Equality for Free Energy Differences. Physical Review Letters. 1997 Apr; 78(14):2690–2693. doi: 10.1103/Phys-RevLett.78.2690.

[94] Shirts MR, Chodera JD. Statistically Optimal Analysis of Samples from Multiple Equilibrium States. The Journal of Chemical Physics. 2008 Sep; 129(12):124105. doi: 10.1063/1.2978177.

[95] Izadi S, Onufriev AV. Accuracy Limit of Rigid 3-Point Water Models. The Journal of Chemical Physics. 2016 Aug; 145(7):074501. doi: 10.1063/1.4960175.

[96] Dodda LS, Cabeza de Vaca I, Tirado-Rives J, Jorgensen WL. LigParGen Web Server: An Automatic OPLS-AA Parameter Generator for Organic Ligands. Nucleic Acids Research. 2017 Jul; 45(W1):W331–W336. doi: 10.1093/nar/gkx312.

[97] Vassetti D, Pagliai M, Procacci P. Assessment of GAFF2 and OPLS-AA General Force Fields in Combination with the Water Models TIP3P, SPCE, and OPC3 for the Solvation Free Energy of Druglike Organic Molecules. Journal of Chemical Theory and Computation. 2019 Mar; 15(3):1983–1995. doi: 10.1021/acs.jctc.8b01039.

[98] Bultinck P, Van Alsenoy C, Ayers PW, Carbo-Dorca R. Critical analysis and extension of the Hirshfeld atoms in molecules. The Journal of Chemical Physics. 2007; 126(14):144111. https://doi.org/10.1063/1.2715563, doi: 10.1063/1.2715563.

[99] Verstraelen T, Vandenbrande S, Heidar-Zadeh F, Vanduyfhuys L, Van Speybroeck V, Waroquier M, Ayers PW. Minimal Basis Iterative Stockholder: Atoms in Molecules for Force-Field Development. Journal of Chemical Theory and Computation. 2016 Aug; 12(8):3894–3912. doi: 10.1021/acs.jctc.6b00456.

[100] Kusalik PG, Svishchev IM. The Spatial Structure in Liquid Water. Science. 1994 Aug; 265(5176):1219–1221. doi: 10.1126/science.265.5176.1219.

[101] Li H, Chowdhary J, Huang L, He X, MacKerell AD, Roux B. Drude Polarizable Force Field for Molecular Dynamics Simulations of Saturated and Unsaturated Zwitterionic Lipids. Journal of Chemical Theory and Computation. 2017 Sep; 13(9):4535–4552. doi: 10.1021/acs.jctc.7b00262.

[102] Kamath G, Kurnikov I, Fain B, Leontyev I, Illarionov A, Butin O, Olevanov M, Pereyaslavets L. Prediction of Cyclohexane-Water Distribution Coefficient for SAMPL5 Drug-like Compounds with the QMPFF3 and ARROW Polarizable Force Fields. Journal of Computer-Aided Molecular Design. 2016 Nov; 30(11):977–988. doi: 10.1007/s10822-016-9958-4.

[103] Bannan CC, Calabró G, Kyu DY, Mobley DL. Calculating Partition Coefficients of Small Molecules in Octanol/Water and Cyclohexane/Water. Journal of Chemical Theory and Computation. 2016 Aug; 12(8):4015–4024. doi: 10.1021/acs.jctc.6b00449.

[104] Niemi GJ, Basak SC, Grunwald G, Veith GD. Prediction of Octanol/Water Partition Coefficient (Kow) with Algorithmically Derived Variables. Environmental Toxicology and Chemistry. 1992 Jul; 11(7):893–900. doi: 10.1002/etc.5620110703.

[105] Tetko IV, Tanchuk VY, Villa AEP. Prediction of N-Octanol/Water Partition Coefficients from PHYSPROP Database Using Artificial Neural Networks and E-State Indices. Journal of Chemical Information and Computer Sciences. 2001 Sep; 41(5):1407–1421. doi: 10.1021/ci010368v.

[106] Vraka C, Nics L, Wagner KH, Hacker M, Wadsak W, Mitterhauser M. Log P, a Yesterday’s Value? Nuclear Medicine and Biology. 2017 Jul; 50:1–10. doi: 10.1016/j.nucmedbio.2017.03.003.

[107] Glomme A, März J, Dressman JB. Comparison of a Miniaturized Shake-Flask Solubility Method with Automated Potentiometric Acid/Base Titrations and Calculated Solubilities. Journal of Pharmaceutical Sciences. 2005 Jan; 94(1):1–16. doi: 10.1002/jps.20212.

[108] Slater B, McCormack A, Avdeef A, Comer JEA. PH-Metric logP.4. Comparison of Partition Coefficients Determined by HPLC and Potentiometric Methods to Literature Values. Journal of Pharmaceutical Sciences. 1994 Sep; 83(9):1280–1283. doi: 10.1002/jps.2600830918.

[109] Mobley DL, Bayly CI, Cooper MD, Dill KA. Predictions of Hydration Free Energies from All-Atom Molecular Dynamics Simulations. J Phys Chem B. 2009 Jan; 113:4533–4537.

[110] Cournia Z, Allen B, Sherman W. Relative Binding Free Energy Calculations in Drug Discovery: Recent Advances and Practical Considerations. Journal of Chemical Information and Modeling. 2017 Dec; 57(12):2911–2937. doi: 10.1021/acs.jcim.7b00564.

[111] Case D, Berryman J, Betz R, Cerutti D, Cheatham Iii T, Darden T, Duke R, Giese T, Gohlke H, Goetz A, et al. AMBER 2015. University of California, San Francisco. 2015;.

[112] Martínez L, Andrade R, Birgin EG, Martínez JM. PACKMOL: A Package for Building Initial Configurations for Molecular Dynamics Simulations. Journal of Computational Chemistry. 2009 Oct; 30(13):2157–2164. doi: 10.1002/jcc.21224.

[113] Eastman P, Swails J, Chodera JD, McGibbon RT, Zhao Y, Beauchamp KA, Wang LP, Simmonett AC, Harrigan MP, Stern CD, Wiewiora RP, Brooks BR, Pande VS. OpenMM 7: Rapid Development of High Performance Algorithms for Molecular Dynamics. PLOS Computational Biology. 2017 Jul; 13(7):e1005659. doi: 10.1371/journal.pcbi.1005659.

[114] Hopkins CW, Le Grand S, Walker RC, Roitberg AE. Long-Time-Step Molecular Dynamics through Hydrogen Mass Repartitioning. Journal of Chemical Theory and Computation. 2015 Apr; 11(4):1864–1874. doi: 10.1021/ct5010406.

[115] Rizzi A, Chodera J, Naden L, Beauchamp K, Albanese S, Grinaway P, Rustenburg B, Saladi S, Boehm K, choderalab/yank: Bugfix release. Zenodo; 2018. https://doi.org/10.5281/zenodo.1447109, doi: 10.5281/zenodo.1447109.

[116] Gerber PR. Charge Distribution from a Simple Molecular Orbital Type Calculation and Non-Bonding Interaction Terms in the Force Field MAB. Journal of Computer-Aided Molecular Design. 1998 Jan; 12(1):37–51. doi: 10.1023/A:1007902804814.

[117] Milletti F, Storchi L, Sforna G, Cruciani G. New and Original p *K* _a_ Prediction Method Using Grid Molecular Interaction Fields. Journal of Chemical Information and Modeling. 2007 Nov; 47(6):2172–2181. doi: 10.1021/ci700018y.

