## Supplementary material for "Assessing the accuracy of octanol-water partition coefficient predictions in the SAMPL6 Part II log *P* Challenge": SAMPL6-supplementary-documents.tar.gz: logP_challenge_instructions.md

# SAMPL6 log *P* Challenge Instructions

SAMPL6 was originally announced as featuring a log *D* prediction challenge, but there were difficulties in the collection of experimental data as we explain in “Experimental Details” section below. 
We were instead able to collect experimental neutral-compound log partition coefficients (log *P*) for a subset of the SAMPL6 pKa challenge compounds. 
Thus, these form the basis of  SAMPL6 Part II -- a log *P* prediction challenge commencing immediately. 
We hope that the log *P* challenge will be useful in investigating sources of modeling errors that impact solvation, partition, and affinity predictions other than protonation state related errors that were prominent in SAMPL5 log *D* challenge.  

This challenge consists of predicting the octanol-water partition coefficients (log *P*) of 11 small molecules that resemble fragments of small molecule protein kinase inhibitors. 
Our aim is to evaluate how well current models can capture the transfer free energy of small molecules between different solvent environments through blind predictions.

log *P*<sub>oct/wat</sub> = log<sub>10</sub> ( [unionized solute]<sub>octanol</sub> / [unionized solute]<sub>water</sub> ) 

Participants are encouraged to submit articles evaluating their methods to the coming special issue or section of the Journal of Computer-Aided Molecular Design special issue targeting September 2019. 
The challenge will culminate with a joint D3R/SAMPL workshop.
The following subsections describe the molecules included in this challenge, the experimental conditions and measurements, the quantities to be predicted, and how prediction results must be submitted.

## Challenge Timeline

- Nov 1, 2018  -  SAMPL6 Part II Challenge start date
- Mar 22, 2019  -  Challenge submissions due 
- Mar 25, 2019  -  Experimental data release date 
- May 16, 2019  -  SAMPL6 log *P* challenge virtual workshop
- Aug 22-23, 2019  -  Joint D3R/SAMPL workshop, San Diego 
- Oct 15, 2019  -  JCAMD special issue submissions due 

Your predictions must be uploaded on the [D3R SAMPL6 web-page](https://drugdesigndata.org/about/sampl6) by March 22nd, 2019. 
The experimental results will be released immediately after the challenge closes. 

# Motivation

Distribution coefficients (log *D*) replaced hydration free energies in SAMPL5 challenge and provided great insight into the importance of modeling a variety of physical effects (overview doi:10.1007/s10822-016-9954-8 and experiment doi:10.1007/s10822-016-9971-7; JCAMD special issue https://link.springer.com/journal/10822/30/11/page/1). 
log *D* values capture the same properties as hydration free energies, namely, solvation in the respective solvents. 
In many SAMPL5 submissions, they were predicted as if they were partition coefficients (log *P*). 
The difference between log *D* (which reflects the transfer free energy at a given pH including the effects of accessing all equilibrium protonation states of the solute in each phase) and log *P* (which reflects the free energy of transfer for the neutral form only) proved particularly important. 
In some cases, other effects like the presence of a small amount of water in cyclohexane may also have played a role.

Because the SAMPL5 log *D* challenge highlighted the difficulty in correctly predicting transfer free energies involving protonation states (the best methods with RMSE of 2.5 log units [1]), we aimed to isolate the protonation and partition prediction components into two different challenges in SAMPL6. 
Participants are asked to predict the partition coefficient log *P* of the neutral species between octanol and water phases.

Partition coefficient prediction as a model problem embodies important elements of the physical chemistry of protein-ligand binding affinity prediction, while making it far easier to probe the accuracy of computational tools used to model protein-ligand interactions and to identify and correct sources of error. 
For physical modeling approaches, evaluation of partition coefficient predictions is a means of separating force field accuracy from sampling and protonation state modeling challenges. Protein-ligand binding equilibrium is like a partitioning between two environments: protein binding site and aqueous phase. 
Methods that employ thermodynamic cycles, such as free energy calculations, can therefore employ similar strategies as they would for calculating binding affinities.  
On the other hand, partition coefficient prediction omits the difficulties of conformational sampling of proteins and treatment of protonation states. 

We believe SAMPL6 logP challenge will benefit improvement of solvation, partition/distribution coefficient and affinity prediction methods, as well as other components of molecular modeling methods such as force fields, sampling algorithms, and prediction of prospective model inaccuracies. 
One of the goals of this challenge is to encourage prediction of model uncertainties (an estimate of the inaccuracy with which your model predicts the physical property), since the ability to tell when methods will be successful or not would be very useful for increasing the application potential and impact of computational methods.

## 11 small molecules are included in log *P* challenge

![SAMPL6_logP_compounds.jpg](images/SAMPL6_logP_compounds.jpg)

**Fig 1. SAMPL6 logP Challenge molecules.** These molecules are a subset of the SAMPL6 pKa Challenge set. 

The SAMPL6 log *P* prediction challenge is composed of a subset of kinase inhibitor fragment-like small molecules from SAMPL6 pKa challenge. We kept the Molecule IDs assigned to these molecules for the pKa challenge. 
These 11 small molecules were selected due to their suitability for potentiometric log *P* measurements. 
Six of them represent the 4-amino quinazoline scaffold. 

A list of SAMPL6 log *P* Challenge small molecules, isomeric SMILES, and molecule IDs can be found here: [`/physical_properties/logP/molecule_ID_and_SMILES.csv`](/physical_properties/logP/molecule_ID_and_SMILES.csv). 
Counterions, where present in solid formulations (see Experimental Details section below), were included in canonical isomeric SMILES for the sake of completeness, although no significant effect is expected from the presence of chloride counterions as experiments were conducted in ionic-strength adjusted medium with KCl.


## Experimental details

Experimental log *P* values were collected using potentiometric log *P* (pH-metric log *P*)[2] measurements with a [Sirius T3 instrument (Pion)](http://www.sirius-analytical.com/products/t3) by Mehtap Isik from the Chodera Lab at MSKCC with the support of the Preformulation Group, Pharmaceutical Sciences, MRL, Merck & Co., Inc, especially Dorothy Levorse, Timothy Rhodes, and Brad Sherborne.

The pH-metric log *P* measurement method of the Sirius T3 instrument [3-6] is based on determining a lipophilicity profile directly from acid-base titrations in a dual-phase water-partition solvent system. 
In this method, multiple potentiometric acid-base titrations are performed in the presence of different ratios of octanol and water to observe the ionization and partitioning behavior of the analyte. A
s the relative volume ratio of octanol to water changes, a shift in apparent pK<sub>a</sub> (p<sub>o</sub>K<sub>a</sub>) is observed due to partitioning to the octanol phase. 
Equations of partitioning and ionization are solved to determine the log *P* of the neutral and ionic species. 
To use this method, aqueous pKa value(s) must be known and analytes must be sufficiently water soluble.

Aqueous pKa values of log *P* challenge compounds were previously determined using UV-metric measurements with the Sirius T3, as described in [the SAMPL6 pKa challenge](physical_properties/pKa/). 
These values were used as input for pH-metric log *P* measurements. 
Unfortunately, only 11 of 24 compounds of pKa challenge set were found to be suitable for pH-metric log *P* measurements.

Three independent replicates of log *P* measurements in 1-octanol and water biphasic systems were performed at 25°C. 
Samples were prepared by weighing 1-3 mg of analyte in solid powder form into glass vials. 
The weights of analytes were inputted to the analysis software. 
The rest of the sample preparation was controlled by the automated titrator: addition of ionic-strength adjusted (ISA) water and partition solvent, mixing, and sonication. 
ISA water is 0.15 M KCl solution which was used to keep ionic strength constant during the experiment. 
Partition solvent is 1-octanol saturated with ISA water, which was prepared as a 5% ISA water-octanol mixture (by volume) and letting the mixture phases separate. 
In some cases to help with kinetic solubility issues of the analytes, solid samples were predosed manually with 80-100 uL octanol before the addition of ISA water and partition solvent.

![logP_experiment_design_figure.jpg](images/logP_experiment_design_figure.jpg)

**Fig 2. Mean molecular charge profiles of pH-metric titrations for log *P* measurement of a monoprotic base [2].**
Green curve represents the predicted ionization profile in aqueous solution (without octanol) calculated based on experimental pKa. 
Blue curves are titration curves shifted in the presence of different amounts of octanol. 
pK<sub>a</sub> or p<sub>o</sub>K<sub>a</sub> are determined as the pH value at which half-maximum point of mean molecular charge is reached. 
Water-octanol volume ratios of three titrations need to be selected so that they lead to distinct p<sub>o</sub>K<sub>a</sub> values for an accurate determination of log *P*. 
**(A)** Modelled ionization profile of a good experiment with well separated p<sub>o</sub>K<sub>a</sub> values. 
**(B)** An experiment with overlapping p<sub>o</sub>K<sub>a</sub> values. 
It is necessary in case **(B)** to adjust the octanol-water ratios for better, more resolved log *P* measurements. 

For each replicate of log *P* measurement, three sequential automated acid-base titrations were performed at three different volume ratios of octanol and water, using 0.5 M KOH and HCl titrants while monitoring pH with a pH electrode. 
Additional octanol volumes were dispensed before each titration to achieve target octanol-water ratios. 
Experiments were designed so that the total volume in the analysis vial did not exceed 3 mL by the end of three titrations. 

When an ionizable substance is titrated in a two-phase system, the apparent pKa observed in the titration shifts due to differential partitioning of neutral and ionized species into the nonaqueous phase. 
The p<sub>o</sub>K<sub>a</sub> value is the apparent pKa in the presence of octanol and it is dependent on the ratio of the water and octanol phases. 
The p<sub>o</sub>K<sub>a</sub> shifts up with monoprotic acids and down with monoprotic bases. 
The shift in p<sub>o</sub>K<sub>a</sub> is directly proportional to the log *P* of the compound and the ratio of octanol to water. 
For a monoprotic base, partition coefficient of neutral species (P<sup>0</sup>) relates to pKa and p<sub>o</sub>K<sub>a</sub> as follows, where r is the volume ratio of nonaqueous phase to aqueous phase [2]:

![P_and_pKa_equation_for_logP_experiments.jpg](images/P_and_pKa_equation_for_logP_experiments.jpg)

Analysis of experimental data was performed with the Sirius T3 Refinement Software. 
First, simulated titration curves were constructed using aqueous pKa values, predicted log *P* values, input analyte concentration, and volumes of aqueous and organic phases. 
Collected experimental data points were used to refine the model parameters (logP of neutral species, log *P* of ionic species, analyte concentration factor, carbonate content, acidity error) to determine the log *P* values of neutral species and ions [5]. 
pH-metric log *P* measurements have the potential to determine partition coefficient of the ions (log *P*<sup>1</sup>) in addition to logP of neutral species (log *P*<sup>0</sup>). 
It was however very challenging to design experiments to capture log *P*<sup>1</sup> values due to volumetric limitations and potentially large shifts that may move p<sub>o</sub>K<sub>a</sub> values out of the measurable pH range (2-12). 
Therefore, we designed the experiments to capture only the partition coefficient of neutral species (log *P*<sup>0</sup>) accurately. The SAMPL6 log *P* prediction challenge will only involve prediction of neutral species. 
Experimental protocols were optimized iteratively by adjusting octanol-water ratios, analyte concentration, and pH interval of the titration.  

An example of pH-metric log *P* determination results is shown in **Figure 3** for phenol (which is not part of the prediction challenge). The report of phenol log *P* measurement can be found in [/physical_properties/logP/example_experimental_data](/physical_properties/logP/example_experimental_data) directory.

![phenol_logP_figure.jpg](images/phenol_logP_figure.jpg)

**Fig 3.** Illustration of octanol-water log *P* measurement of phenol (log *P*=1.49) using the pH-metric log *P* method of the Sirius T3. Since phenol is a monoprotic acid, octanol partitioning causes higher p<sub>o</sub>K<sub>a</sub> values than aqueous pKa. 
**(A)** Triangles represent experimental data points collected during the octanol-ISA water titrations and solid lines represent the ionization and partitioning model fit to the data. 
The black line is the titration curve in aqueous media without octanol and based on the aqueous pKa. 
Blue, red, and green triangles represent three sequential titrations with increasing logR (log<sub>10</sub> of octanol to water volumetric ratio) that show shifted p<sub>o</sub>K<sub>a</sub> values. 
**(B)** Inspection of the logR profile aids experimental design to achieve 3 data points with well separated p<sub>o</sub>K<sub>a</sub> values by adjusting the octanol-water ratios.

We attempted measuring the log *P* for all 24 SAMPL6 pKa challenge compounds, but the pH-metric log *P* measurement method was suitable for only a subset of 11 molecules which were included in the study. 
Potential factors limiting the log *P* measurement of more molecules were low water solubility within the pH range of titration, limitation of total sample volume that restricts range of achievable octanol and water ratios, p<sub>o</sub>K<sub>a</sub> values shifting out of the measurable pH range of 2-12 (especially high acidic pKas and low basic pKas), and log *P* values out of the dynamic range of the experimental methodology. 
We only included small molecules with high quality pH-metric log *P* measurements in this challenge.

## Computational prediction methods

You may use any method(s) you like to generate your predictions; e.g., molecular mechanics or quantum mechanics, MD with implicit or explicit solvent, QSPR, empirical logP prediction methods etc.

## Possible factors to consider in your approach

Partition coefficients are determined by the difference in solvation free energies of the relevant species in the different phases. 
In fact, they can be estimated from gas-to-solvent transfer free energies into the different solvents. 
However, it is important to note that the experimental reality may be more complicated for several reasons.

The experiments are performed on phase-separated water and octanol.  
The aqueous phase had 0.15 M KCl to achieve constant ionic strength during experiments. 
After mixing, the water and octanol phases may no longer be pure water and octanol. 
Please consider mutual solubility of water and octanol.
Water and/or salts may be found in the octanol phase. 
Octanol may be found in the aqueous phase. 
The mole fraction of water in octanol was measured as 0.2705 ± 0.0028 at 25°C [7].

Additionally, the solute can impact the distribution of water and octanol themselves. 
For example, carboxylic acids and some other solutes strongly bind one or more water molecules even in the nonaqueous phase, at least in some cases [8]. 
Dimerization or oligomerization of solute molecules in one or more of the phases may also impact results; for example, a polar molecule might dimerize in a nonpolar phase, resulting in stabilization in that phase relative to what would be expected based on the monomer’s transfer free energy [8] .

Finally, the possibility of tautomerization should be considered. 
log *P* measurements capture partition of neutral species which can consist of multiple tautomers with significant populations or the major tautomer may not be the one given in input file ([molecule_ID_and_SMILES.csv](/physical_properties/logP/molecule_ID_and_SMILES.csv)). 
Shifts in tautomeric state populations on transferring between phases is also a possibility.

We are not aware which, if any, of these potential complications are relevant for this competition. 
We mention them only to ensure all participants are aware of them.

## Instructions and submission template 
- Participants must use the provided template file ([logP_prediction_template.csv](physical_properties/pKa/submission_template/logP_prediction_template.csv)) found in `physical_properties/pKa/submission_template/` directory to upload predictions to [the SAMPL website](https://drugdesigndata.org/about/sampl6).

- Fill one template file for each approach to create a submission that contains predictions for all molecules with predicted using one method. You may submit predictions from multiple methods, but you should fill a separate template file for each different method.

- You may report only 1 log *P* value for each molecule per method.

- It is mandatory to submit predictions for all 11 molecules. Incomplete submissions will not be accepted.

- Report log *P* values to two decimal places (e.g. 2.71).

- Report the standard error of the mean (SEM) as a measure of statistical uncertainty (imprecision) for your method.  log *P* SEM should capture variation of predicted values of the same method over repeated calculations.

- Report the model uncertainty of your log *P* prediction --- the predicted accuracy of your method [1,9]. This is not a statistical uncertainty. Rather, the model uncertainty is an estimate of how well your predicted values are expected to agree with experimental values. For example, for classical simulation approaches based on force fields, this could measure how well you expect the force field will agree with experiment for this compound. The model uncertainty could be global or different for each molecule. For example, reference calculations in SAMPL5 log *D* challenge estimated the model uncertainty as the root mean squared error (RMSE) between predicted and experimental values for a set of molecules with published cyclohexane-water partition coefficients. 

- Lines beginning with a hash-tag (#) may be included as comments. These and blank lines will be ignored during analysis.

- The file must contain the following four components in the following order: your predictions, a name for your computational protocol, a list of the major software packages used, prediction method category, and a long-form methods description. Each of these components must begin with a line containing only the corresponding keyword: `Predictions:`, `Name:`, `Software:`, `Category:`, and `Method:`, as illustrated in the example files. Example submission files can be found in [`physical_properties/logP/example_submission_file/`](physical_properties/logP/example_submission_file/) directory to illustrate expected format when filling submission templates.

- For Method Category section please state if your prediction method can be better classified as a empirical modeling method, physical modeling method, mixed (both empirical and physical) or other, using the category labels `Empirical`, `Physical`, `Mixed`, or `Other`. Empirical models are prediction methods that are trained on experimental data, such as QSPR, machine learning models, artificial neural networks etc. Physical models are prediction methods that rely on the physical principles of the system such as molecular mechanics or quantum mechanics based methods to predict molecular properties. If your method takes advantage of both kinds of approaches please report it as “Mixed”. If these categories do not match your method, report as “Other”. If you choose “Mixed” or “Other” categories,  please explain your decision in the beginning of Method Description section.

- Names of the prediction files must have three sections separated by `-`: predicted property `logP`, and your name and must end with an integer indicating the number of prediction set. For example, if you want to submit one prediction, you would name it `logP-myname-1.csv`, where `myname` is arbitrary text of your choice. If you submit three prediction files, you would name them `logP-myname-1.csv`, `logP-myname-2.csv`, and `logP-myname-3.csv`.

- Prediction files will be machine parsed, so correct formatting is essential. Files with the wrong format will not be accepted.

## Evaluation strategy for computational logP predictions

Predicted log *P* values and their statistical uncertainties will be directly compared to experimental octanol-water log *P* measurements. 
We will evaluate methods using various error metrics including, RMSE, MAE, R<sup>2</sup>, linear regression slope etc. We will give priority to RMSE and MAE over correlation based statistics due to limited number of data points and dynamic range in this experimental data set. 
We will evaluate model uncertainties in comparison to calculated RMSE between experimental and predicted values of each method to evaluate how successful each method was at estimating its inaccuracy. 
QQ-plots of model uncertainties will be plotted to understand if model uncertainty was under or overestimated. 
It is possible that we will also evaluate metrics such as weighted RMSE where predictions with low model uncertainties count more than those with high model uncertainties.

## Submission of multiple predictions

Some participants use SAMPL to help evaluate various computational methods. 
To accommodate this, multiple prediction sets from a single research group or company are allowed, even for the same type of predictions if they are made by different methods. 
If you would like to submit predictions from multiple methods, you should fill a separate submission template files for each different method. See "Uploading your predictions" section below for requirements on how to name submission files.

## Uploading your predictions

D3R website SAMPL6 page is now ready to accept your uploaded predictions. You may upload your predictions as soon as you are ready and until the submission deadline. If you want to upload predictions of more than one method, each must be uploaded as a separate file. Please use the template provided to create the submission files, as the predictions will be parsed and analyzed with automated scripts. Please include all predictions made with the same method in one file.  

**All submissions are required to contain logP predictions of all 11 compounds. Incomplete submissions - such as for a subset of compounds - will not be accepted.**   

To submit your predictions go to the [D3R SAMPL6 webpage](https://drugdesigndata.org/about/sampl6), log in to your account, and select the **logP Prediction** challenge on the left menu. Then click **Submissions** on the green ribbon.  

Add your file(s) using **Submit Your Data** section. Submitted files will be checked by a validation script for conforming to submission template rules. After clicking **Start upload**, please pay attention to any error message that may show up at the bottom of the submission page. When your files are successfully submitted you should be able to see them listed in the upper **Manage Your Data** section, even after you refresh the webpage or re-log in to your account.  

Please check that **Manage Your Data** section includes a complete list of your submission files.  

## SAMPL6 Part II Workshops

There will be two opportunities for participants to discuss their results.  **SAMPL6 log *P* Challenge Virtual Workshop** will meet on **May 16, 2019**, for sharing methods and discussion of preliminary results.  All participants are also invited to the third in-person **D3R and SAMPL Workshop** which is scheduled for **August 22-23, 2019**, in San Diego to discuss further findings and lessons-learned, as wells as D3R and SAMPL projects more broadly. Note that the workshop is right before the ACS National Meeting in San Diego with the theme of “Chemistry of Water” .


### 1. The SAMPL6 Log *P* Challenge Virtual Workshop

We had virtual pre-workshop on the SAMPL6 LogP challenge on Thursday, May 16, beginning at 7 am US Pacific time.  

#### Overview/Purpose

The purpose of this workshop was to go over a preliminary evaluation of results, begin considering analysis and lessons learned, and nucleate opportunities for follow up and additional discussion. Part of the goal is to facilitate discussion so that participants can work *together* to maximize lessons learned in the lead up to the August in-person D3R/SAMPL workshop and the JCAMD special issue.
We hope this early workshop will result in additional follow-up afterwards.

#### Agenda

A tentative agenda for this workshop is (all times US Pacific (PDT; UTC-7)):
- **7-7:10 am**: *Welcome, introductions, and purpose*; David Mobley (UCI)
- **7:10-7:50 am**: *Experiments and overview of results*, Mehtap Isik (MSKCC); 30 minute talk + 10 minutes of questions/discussion.
- **7:50-8:05 am**: Andrew Paluch (Miami University, Ohio)
- **8:05-8:35 am**: Christoph Loschen (Cosmologic), *COSMO-RS based predictions for the SAMPL6 logP challenge*; 20 minute talk + 10 minutes of questions/discussion
- **8:35-8:55 am**: Nicolas Tielker (TU Dortmund); 12 minute talk + 8 minutes of questions/discussion
- **8:55-9:15 am**: Esteban Vohringer-Martinez (Univ. Concepción); 15 minute talk + 5 minutes of questions/discussion.
- **9:15 am-9:25 am**: Alexey Nikitin; 5 minute talk + 5 minutes of discussion.
- **9:25-10 am**: Discussion and follow up opportunities.

#### Connection details

This is a Zoom meeting/workshop. You may join using [this link](https://uchealth.zoom.us/j/687748918). You should download the software in advance if you do not already have it. If you plan on attending and are not a participant, please ensure your video is turned off and your microphone is muted to improve the experience for other participants.
Dial-in options for audio are also available; contact David Mobley if you need to dial in.

#### Presentations and video record

Slide decks of talks given in SAMPL6 Log *P* Challenge virtual workshop can be found here:  
[SAMPL6-logP-challenge-virtual-workshop GitHub repository](https://github.com/choderalab/SAMPL6-logP-challenge-virtual-workshop)

[SAMPL6 logP virtual workshop video](https://www.youtube.com/watch?v=FWUPXG8U3UE) is also available.


### 2. The 2019 D3R/SAMPL Joint Workshop

The next [face-to-face workshop](https://drugdesigndata.org/about/d3r-2019-workshop) is Aug. 22-23, 2019 in San Diego, joint with D3R.
Participation is relatively limited but we are checking on whether virtual attendance is a possibility.

## Publishing in the special issue of JCAMD 

In coordination with Terry Stouch, we are planning a special issue of *J. Comp. Aided Mol. Design* (JCAMD) focused on the SAMPL6 log *P* challenge. All participants are welcome to submit manuscripts. The submission deadline for this issue is extended Oct. 15, 2019, with submissions opening prior to our August 2019 workshop for those who would like to submit early. As usual, papers are expected to appear online in advance of their publication in the special issue as soon as they are ready unless otherwise requested.

### Review criteria for special issue
In general, review criteria for SAMPL special issues are modestly different than for typical journal publications. Specifically, since these are blind prediction challenges, the community feeling has been that participants should be entitled to report what they did even if the reviewers might feel that it was ill advised, not particularly novel/exciting, or had modest had technical problems. However, papers must still:

- fairly report their results, without overselling or giving an unwarranted sales pitch
- clearly identify and discuss any technical flaws reviews or others might have highlighted
- provide adequate details (and supporting materials) so that others can reproduce the work
and otherwise ensure that they meet the standards of the journal.

Additionally, as a particular focus of SAMPL is on lessons learned, authors are urged to pay devote careful attention to highlighting what was learned from participation and how it might be of benefit to the field or to others employing similar methodologies.

###  Including submission IDs in publications
If you are preparing manuscripts reporting on your predictions in SAMPL6 Part II log *P* challenge, we kindly ask you to report the submission ID of your blind submission in your publication. Submission IDs are 5-character unique identifiers assigned to each submission and can be found in [this table](https://github.com/MobleyLab/SAMPL6/tree/master/physical_properties/logP/analysis#submission-ids-for-log-p-prediction-methods). We hope that including submission IDs in manuscripts will help to cross-reference between the papers in the special issue.

## Files provided

- `/physical_properties/logP/molecule_ID_and_SMILES.csv` - CSV file that indicates SAMPL6 logP challenge molecule IDs and isomeric SMILES.
- `/physical_properties/logP/submission_template/logP_prediction_template.csv` - An empty prediction submission template files.
-  `/physical_properties/logP/example_submission_file/logP-MehtapIsikExampleFile-1.csv` - An example submission file filled with random values to illustrate expected format.
- `/physical_properties/logP/example_experimental_data/` - This directory contains the experimental report of pH-metric log *P* measurement of phenol with Sirius T3 as an example.

## Problems, questions, and contact

If you notice any issues with any of these files, please contact us via the GitHub issue tracker. 
You are also strongly advised to both sign up for the SAMPL6 e-mail list via the D3R site and sign up for notifications on this GitHub repository in case we have updates.
Please feel free to contact us if you notice any errors in the information provided or have questions about SAMPL6; please use the issue tracker connected with this repository, or for specific questions about logP challenge, use the following email: [].

## References

[1] Bannan, Caitlin C., Kalistyn H. Burley, Michael Chiu, Michael R. Shirts, Michael K. Gilson, and David L. Mobley. “Blind Prediction of Cyclohexane–water Distribution Coefficients from the SAMPL5 Challenge.” *Journal of Computer-Aided Molecular Design* 30, no. 11 (November 2016): 927–44. 

[2] Comer, John, and Kin Tam. Lipophilicity Profiles: Theory and Measurement. Wiley-VCH: Zürich, Switzerland, 2001.

[3] “Sirius T3 User Manual, v1.1.” (Sirius Analytical Instruments Ltd, East Sussex, UK), 2008.

[4] Avdeef, Alex. “PH-Metric Log P. Part 1. Difference Plots for Determining Ion-Pair Octanol-Water Partition Coefficients of Multiprotic Substances.” *Quantitative Structure-Activity Relationships* 11, no. 4 (1992): 510–17.

[5] Avdeef, Alex. “pH-Metric Log P. II: Refinement of Partition Coefficients and Ionization Constants of Multiprotic Substances.” *Journal of Pharmaceutical Sciences* 82, no. 2 (1993): 183–90.

[6] Slater, Bryan, Ann McCormack, Alex Avdeef, and John EA Comer. “Ph-Metric Log P. 4. Comparison of Partition Coefficients Determined by HPLC and Potentiometric Methods to Literature Values.” *Journal of Pharmaceutical Sciences* 83, no. 9 (1994): 1280–83.

[7] Lang, Brian E. “Solubility of Water in Octan-1-Ol from (275 to 369) K.” *Journal of Chemical & Engineering Data* 57, no. 8 (August 9, 2012): 2221–26. https://doi.org/10.1021/je3001427.

[8] Leo, A., Hansch, C., & Elkins, D. (1971). Partition coefficients and their uses. *Chemical Reviews* , 71 (6), 525–616.

[9] Mobley, David L., Karisa L. Wymer, Nathan M. Lim, and J. Peter Guthrie. “Blind Prediction of Solvation Free Energies from the SAMPL4 Challenge.” *Journal of Computer-Aided Molecular Design* 28, no. 3 (March 2014): 135–50. https://doi.org/10.1007/s10822-014-9718-2.
